# Omicron BA.2 specifically evades broad sarbecovirus neutralizing antibodies

**DOI:** 10.1101/2022.02.07.479349

**Authors:** Yunlong Cao, Ayijiang Yisimayi, Fanchong Jian, Tianhe Xiao, Weiliang Song, Jing Wang, Shuo Du, Zhiying Zhang, Pulan Liu, Xiaohua Hao, Qianqian Li, Xiaosu Chen, Lei Wang, Peng Wang, Ran An, Yao Wang, Jing Wang, Peng Yang, Haiyan Sun, Lijuan Zhao, Wen Zhang, Dong Zhao, Jiang Zheng, Lingling Yu, Can Li, Na Zhang, Rui Wang, Xiao Niu, Sijie Yang, Xuetao Song, Linlin Zheng, Zhiqiang Li, Qingqing Gu, Fei Shao, Weijin Huang, Youchun Wang, Zhongyang Shen, Xiangxi Wang, Ronghua Jin, Junyu Xiao, Xiaoliang Sunney Xie

**Author notes:** Correspondence: Yunlong Cao; Zhongyang Shen; Xiangxi Wang; Ronghua Jin; Junyu Xiao; Xiaoliang Sunney Xie. These authors contributed equally (Yunlong Cao, Ayijiang Yisimayi, Fanchong Jian, Tianhe Xiao, Weiliang Song, Jing Wang, Shuo Du, Zhiying Zhang, Pulan Liu, Xiaohua Hao, Qianqian Li, Xiaosu Chen, Lei Wang).

## Abstract

Omicron sub-lineage BA.2 has rapidly surged globally, accounting for over 60% of recent SARS-CoV-2 infections. Newly acquired RBD mutations and high transmission advantage over BA.1 urge the investigation of BA.2’s immune evasion capability. Here, we show that BA.2 causes strong neutralization resistance, comparable to BA.1, in vaccinated individuals’ plasma. However, BA.2 displays more severe antibody evasion in BA.1 convalescents, and most prominently, in vaccinated SARS convalescents’ plasma, suggesting a substantial antigenicity difference between BA.2 and BA.1. To specify, we determined the escaping mutation profiles^1,2^ of 714 SARS-CoV-2 RBD neutralizing antibodies, including 241 broad sarbecovirus neutralizing antibodies isolated from SARS convalescents, and measured their neutralization efficacy against BA.1, BA.1.1, BA.2. Importantly, BA.2 specifically induces large-scale escape of BA.1/BA.1.1-effective broad sarbecovirus neutralizing antibodies via novel mutations T376A, D405N, and R408S. These sites were highly conserved across sarbecoviruses, suggesting that Omicron BA.2 arose from immune pressure selection instead of zoonotic spillover. Moreover, BA.2 reduces the efficacy of S309 (Sotrovimab)^3,4^ and broad sarbecovirus neutralizing antibodies targeting the similar epitope region, including BD55-5840. Structural comparisons of BD55-5840 in complexes with BA.1 and BA.2 spike suggest that BA.2 could hinder antibody binding through S371F-induced N343-glycan displacement. Intriguingly, the absence of G446S mutation in BA.2 enabled a proportion of 440-449 linear epitope targeting antibodies to retain neutralizing efficacy, including COV2-2130 (Cilgavimab)^5^. Together, we showed that BA.2 exhibits distinct antigenicity compared to BA.1 and provided a comprehensive profile of SARS-CoV-2 antibody escaping mutations. Our study offers critical insights into the humoral immune evading mechanism of current and future variants.

## Main

The recent emergence and global spreading of severe acute respiratory syndrome coronavirus 2 (SARS-CoV-2) variant Omicron (B. 1.1.529) has posed a critical challenge to the efficacy of COVID-19 vaccines and neutralizing antibody therapeutic^7–9^. Due to multiple mutations to the spike protein and its receptor-binding domain (RBD) and N-terminal domain (NTD), Omicron can cause severe neutralizing antibody evasion^1,10–13^. The continuous evolution of Omicron BA.1, such as BA.1.1, BA.2 and BA.3, may lead to an even stronger immune escape^7,13,14^.

Omicron sub-lineage BA.2 has rapidly surged worldwide, out-competing BA.1 and accounting for over 60% of recent COVID-19 cases. Compared to the RBD of BA.1, BA.2 contains three additional mutations, including T376A, D405N and R408S, and lacks the G446S and G496S harbored by BA.1 (Extended Data Fig. 1). The S371F on BA.1 is also substituted with S371L in BA.2. Moreover, mutations carried by BA.2 on the NTD largely differ from BA.1, with only one identical mutation, G142D, shared; however, G142D alone is sufficient to induce NTD neutralization antibody escape^13^. Given BA.2’s newly acquired mutations, higher transmissibility, and the ability to cause reinfections among BA.1 convalescents^15^, the immune escape capability and antigenicity of BA.2 require immediate investigation.

### BA.2 shows distinct antigenicity compared to BA.1

To probe the neutralization resistance degree of BA.2, we first performed pseudovirus neutralization assays using D614G and Omicron variants, BA.1, BA.1.1 and BA.2, against the plasma from vaccinated individuals, BA.1 convalescents, and SARS convalescents (Supplementary Table 1). Plasma samples of the volunteers were obtained 4 weeks after the booster shot, or 4 weeks after discharge from hospitalization. In vaccine-boosted individuals, BA.1, BA.1.1 and BA.2 showed no significant difference in plasma neutralization resistance (Fig. 1a-b), which is concordant with several recent research^4,16^. However, this does not necessarily mean BA.2 displays similar antigenicity with BA.1 and BA.1.1. We found that BA.2 caused stronger immune evasion than BA.1 and BA.1.1 in vaccinated individuals infected with Omicron BA.1, and most prominently in vaccinated SARS convalescents (Fig. 1c-d). This implies that BA.2 may specifically evade broad sarbecovirus neutralizing antibodies, which are substantially enriched in vaccinated SARS convalescents (Fig. 1e)^17^. Indeed, stronger SARS-CoV-1 neutralizing ability tends to be associated with a sharper drop in neutralizing potency against BA.2.

**Fig. 1.**
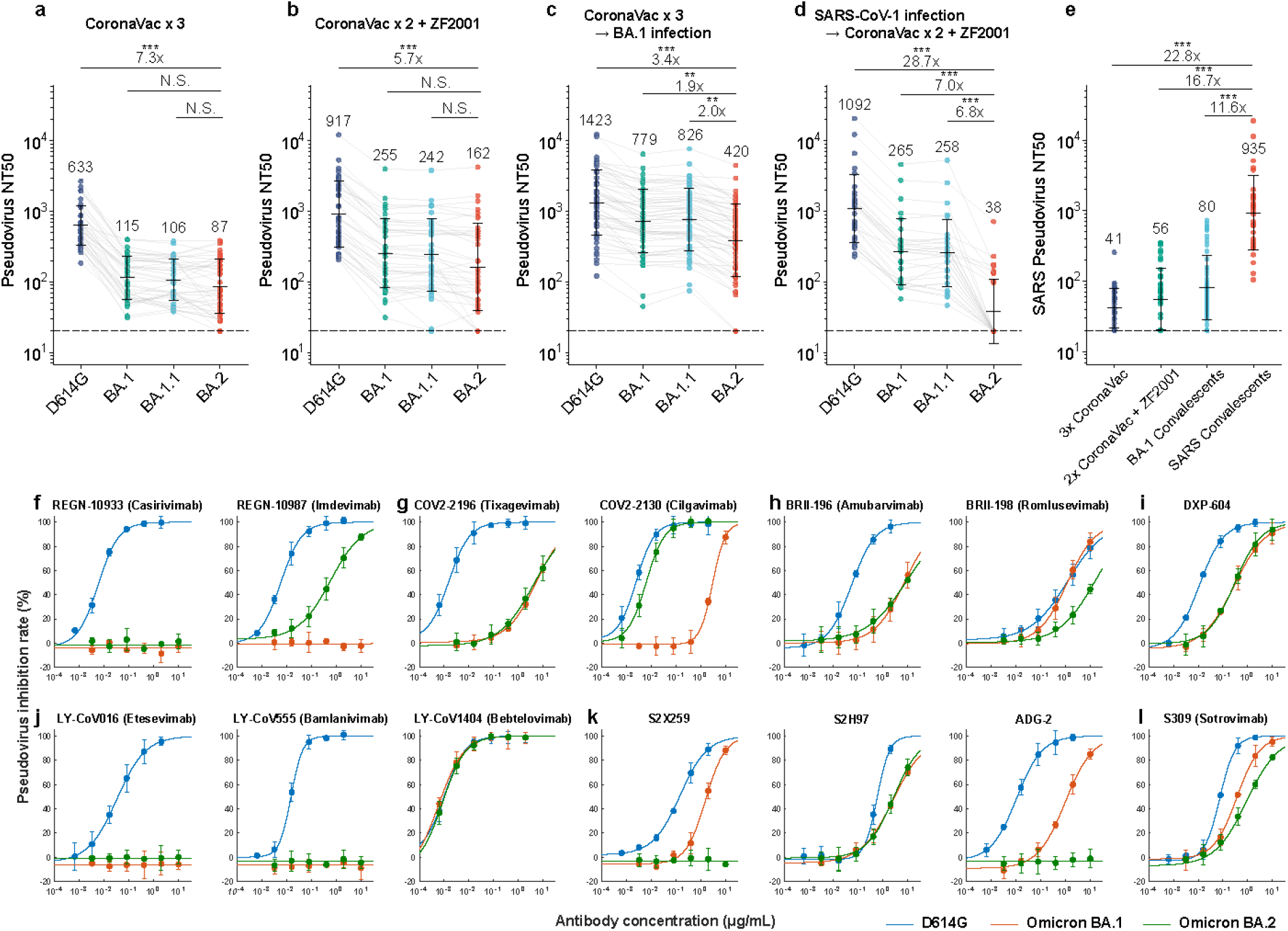
BA.2 shows distinct antigenicity to plasma and therapeutic antibodies compared to BA.1. **a**-**d**, SARS-CoV-2 variants pseudovirus neutralization by plasma from vaccinated and convalescent individuals. **a**, Individuals received 3 doses of CoronaVac (n=40). **b**, Individuals received 2 doses of CoronaVac and 1 dose of ZF2001 (n=39). **c**, BA.1 convalescents who had received 3 doses of CoronaVac before infection (n=54). **d**, SARS convalescents who received 2 doses of CoronaVac and 1 dose of ZF2001 (n=30). **e**, SARS-CoV-1 pseudovirus neutralization by plasma from individuals mentioned above. Average NT50 are annotated above the points. **f-k**, Neutralizing activities against SARS-CoV-2 D614G, BA.1 and BA.2 spike-pseudotyped VSV of RBD neutralizing antibodies. **f-j**, Therapeutic antibodies specific to SARS-CoV-2. **k-l**, Broad sarbecovirus antibodies. P-values are calculated using two-sided Wilcoxon test of paired samples. ***: p<0.001, **: p<0.01, *: p<0.05. All pseudovirus neutralization assays were conducted in biological duplicates.

Next, we examined the reaction difference between BA.1 and BA.2 against therapeutic antibodies and published broad sarbecovirus neutralizing antibodies (Fig. 1f-l). BA.2 and BA.1 displayed similar reactivity against Class 1 and 2 antibodies, such that REGN-10933^6^, LY-CoV016^18^ and LY-CoV555^19^ are evaded by both BA.1 and BA.2, while COV2-2196^5^, BRII-196^20^ and DXP-604^21,22^ exhibited comparable efficacy. This is reasonable since BA.2 and BA.1 do not show mutational differences in the receptor-binding motif. However, two major antigenicity differences were observed between BA.1 and BA.2. First, neutralizing antibodies targeting the linear epitope 440-449^1^, such as REGN-10987^6^ and COV2-2130^5^, retained neutralizing potency against BA.2. Second, BA.2 escapes or greatly reduces the efficacy of BA.1 effective broad sarbecovirus neutralizing antibodies, including S2X259^23^, ADG-2^24^ and S309^3^. Notably, DXP-604^21^ and especially LY-CoV1404^25^ demonstrated high potency against both BA.1 and BA.2. Together, these observations support that BA.2 displays distinct antigenicity from BA.1.

### BA.2 evades BA.1 effective broad sarbecovirus neutralizing antibodies

To specify how BA.2 affects SARS-CoV-2 antibody neutralization, we determined the escaping mutation profiles of 714 SARS-CoV-2 RBD neutralizing antibodies using high-throughput yeast display mutant screening that covers all possible single residue substitutions in the wildtype RBD background^1,26–29^ (Supplementary Data 1). The antibody library mostly contains neutralizing antibodies isolated from SARS-CoV-2 convalescents and vaccinees, and also includes 241 broad sarbecovirus neutralizing antibodies isolated from SARS convalescents by performing high-throughput single-cell VDJ sequencing on memory B cells that could cross-bind to both SARS-CoV-1 and SARS-CoV-2 RBD (Extended Data Fig. 2, Supplementary Table 2) ^21,30^. Based on the escaping mutation profiles, 714 antibodies could be unsupervised clustered into 10 epitope groups (Fig. 2, Extended Data Fig. 3, Supplementary Data 2), as described previously^1^. Group A-D recapitulates our previous antibody clustering^1^, in which the members mainly target the ACE2-binding motif^31–35^. Here, the larger collection of bivalent binding antibodies expands the previous Group E and F into E1-E3 and F1-F3, which respectively target the front and backside of RBD (Fig. 2b).

**Fig. 2.**
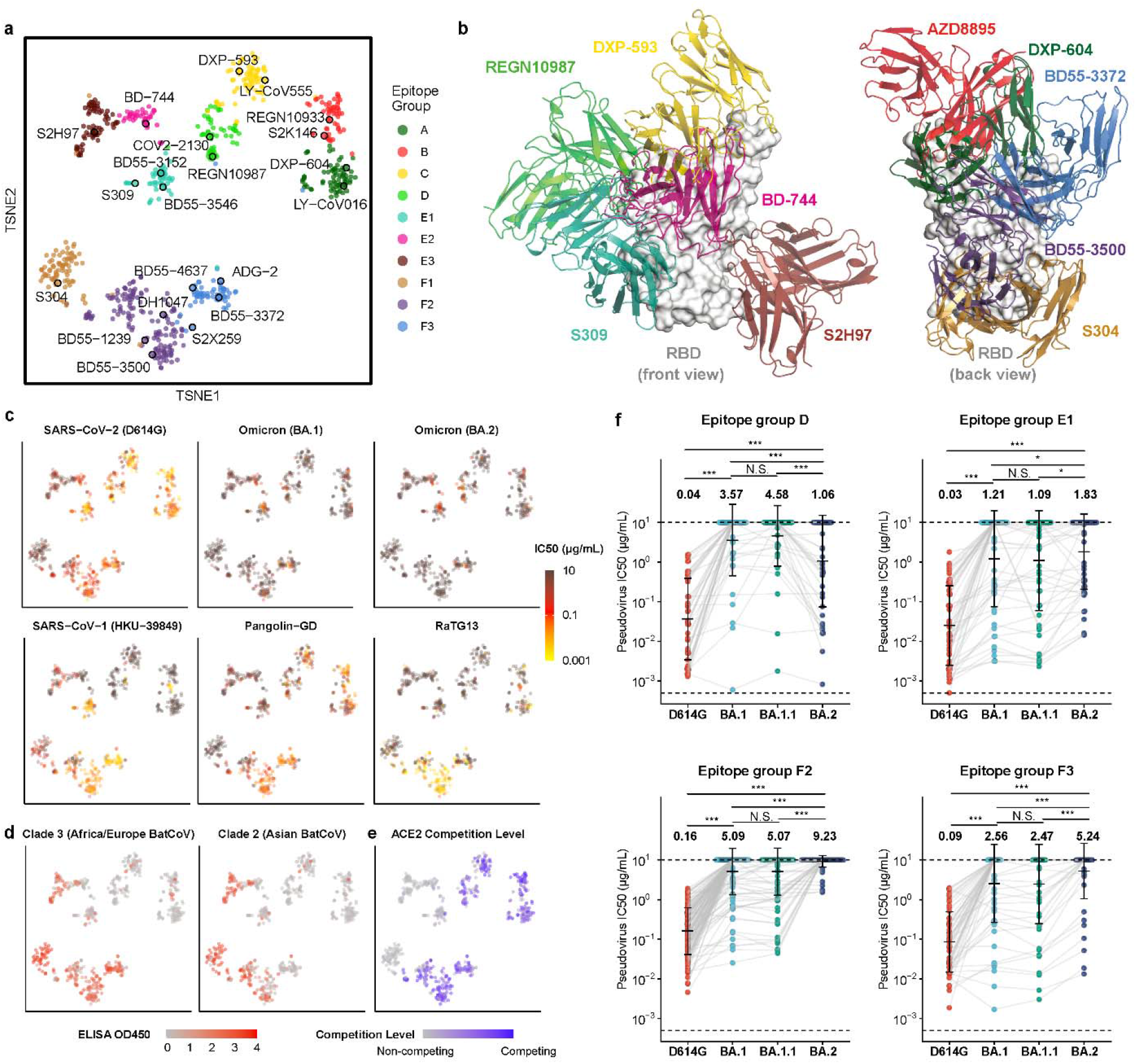
BA.2 evades BA.1 effective broad sarbecovirus neutralizing antibodies. **a**, t-distributed stochastic neighbor embedding (t-SNE) and unsupervised k-means clustering of neutralizing antibodies against SARS-CoV-2 based on antibody escaping profiles. **b**, Representative antibody structures of ten epitope groups. **c**, Neutralization against SARS-CoV-2 (D614G), SARS-CoV-1 (HKU-39849), Pangolin-GD and RaTG13 spike-pseudotyped VSV by 715 RBD antibodies. Color bars indicate IC50 values (μg/mL). **d**, ELISA reactivity to various sarbecovirus RBD of different clades of 715 RBD antibodies. ELISA OD450 is measured using 0.3 μg/mL antigen and 1 μg/mL antibody. Shades of red show the average OD450 of BM48-31 and BtKY72 (clade 3, left), and YN2013, Shaanxi2011, Rs4247, SC2018, Rp3, ZXC21, ZC45 and Anlong112 (clade 2, right). **e**, The ACE2 blocking activity for RBD antibodies. Shades of blue show the competition level measured through competing ELISA. **f**, Distribution of IC50 against SARS-CoV-2 D614G, BA.1, BA. 1.1 and BA.2 of group D, E1, F2, F3 antibodies. Antibodies of which D614G IC50 >2μg/mL are excluded. Average IC50 values are annotated above the bars. P-values are calculated using two-sided Wilcoxon test of paired samples. Antibodies that showed no neutralizing activities against both variants are excluded from calculation of p-values. ***: p<0.001, **: p<0.01, *: p<0.05. All pseudovirus neutralization assays and ELISA measurements were conducted in biological duplicates.

Pseuodovirus neutralizing efficacy of these antibodies against D614G and Omicron variants (BA.1, BA. 1.1, BA.2), SARS-CoV-1, RATG13 and Pangolin-GD is tested, and their binding capability to 21 sarbecovirus RBD is measured through ELISA (Supplementary Table 2, Supplementary Table 3). By projecting each antibody’s experimental measurements against each sarbecovirus onto the t-distributed stochastic neighbor embedding (t-SNE) dimension, we found that antibodies of the same cluster have unified sarbecovirus neutralization potency and binding spectra (Fig. 2c-d, Extended Data Fig. 4). Moreover, the antibodies’ neutralization mechanism also tends to cluster based on the epitope distribution. Antibodies of group E1-E3 and F1 do not compete with ACE2 (Fig. 2e), which results in relatively weaker neutralizing potency compared to ACE2-blocking antibody (Extended Data Fig. 5a), except that group E1 antibodies (S309) still possess high neutralization potency (Fig. 2c), suggesting a rather unique neutralization mechanism. In total, five clusters of antibodies were found to exhibit broad sarbecovirus neutralizing ability with diverse breadth, namely Group E1, E3, F1, F2 and F3 (Fig. 2a-c).

Importantly, four groups of antibodies, including epitope group D, E1, F2 and F3, showed significant reactivity differences against BA.1 and BA.2 (Fig. 2f). Specifically, antibodies of epitope group D, which were largely escaped by BA.1, retained neutralizing potency against BA.2, similar to the behavior of COV2-2130. The reason is clear, which is that BA.2’s absence of G446S enabled a proportion of these antibodies to maintain their binding abilities. Contrarily, BA.1 effective E1 antibodies displayed a systematic reduction in neutralization activity against BA.2, just like S309. Similarly, BA.1 effective F2 and F3 antibodies also showed large-scale escapes caused by BA.2. The mechanisms behind the neutralization loss of those broad sarbecovirus antibodies are unclear. Nevertheless, the comparable neutralization resistance of BA.1 and BA.2 in vaccinated individuals could now be rationalized as the balanced results from the neutralization loss of E1, F2 and F3 antibodies and the retained neutralization efficacy of group D antibodies (Fig. 1a-b). And the increased immune evasion of BA.2 observed in BA.1 convalescents and SARS convalescents are essentially due to the enriched ratio of E1, F2 and F3 broad sarbecovirus antibodies contained in the plasma(Fig. 1c-e).

### S371F on BA.2 induces N343-glycan displacement that hinders antibody binding

To study why BA.2 systematically reduces the neutralization efficacy of E1 antibodies, we need to examine the epitope features shared among E1 antibodies. Thus, we solved the cryo-EM structures of three BA.1 neutralizing E1 antibodies, BD55-3152, BD55-3546 and BD55-5840, in complex with spike proteins using cryo-electron microscopy (cryo-EM) (Extended Data Fig. 6). Like S309, E1 antibodies’ epitope involves an N-linked glycan on N343 (Fig. 3a-d). Besides, members of Group E1 are generally sensitive to the changes of G339, E340, T345 and especially R346, revealed by their escaping mutation profiles (Fig. 3e). Thus, most E1 antibodies could not bind to clade 2 and 3 sarbecoviruses because of the changes of RBD antigenic sites corresponding to G339, E340 and R346, as calculated by multiple sequence alignment (MSA) (Fig. 2d, Fig. 3e). Importantly, BA.1 causes considerable antibody escaping for Group E1 antibodies (Fig. 2f); however, a proportion of E1 antibodies that could tolerate G339D and N440K mutations could retain potent neutralizing ability against BA.1 (Extended Data Fig. 7). Interestingly, despite the importance of R346 to E1 antibodies, the additional R346K carried by BA. 1.1 does not readily affect their efficacy (Fig. 2f). This is foreseeable, as Arg and Lys possess similar chemical properties. In fact, SARS-CoV-1 features a Lys at the corresponding site (K333) (Fig. 3e). The structure of BD55-3152 in complex with the Omicron spike reveals that a CDRL2 Asp (D50) interacts with R346 (Fig. 3a), whereas the structure of BD55-3152 complexed with the SARS-CoV-1 Spike shows that the same Asp also coordinates K333 (Fig. 3b). In contrast to R346K, R346S/T would greatly compromise the binding activities of E1 antibodies.

**Fig. 3.**
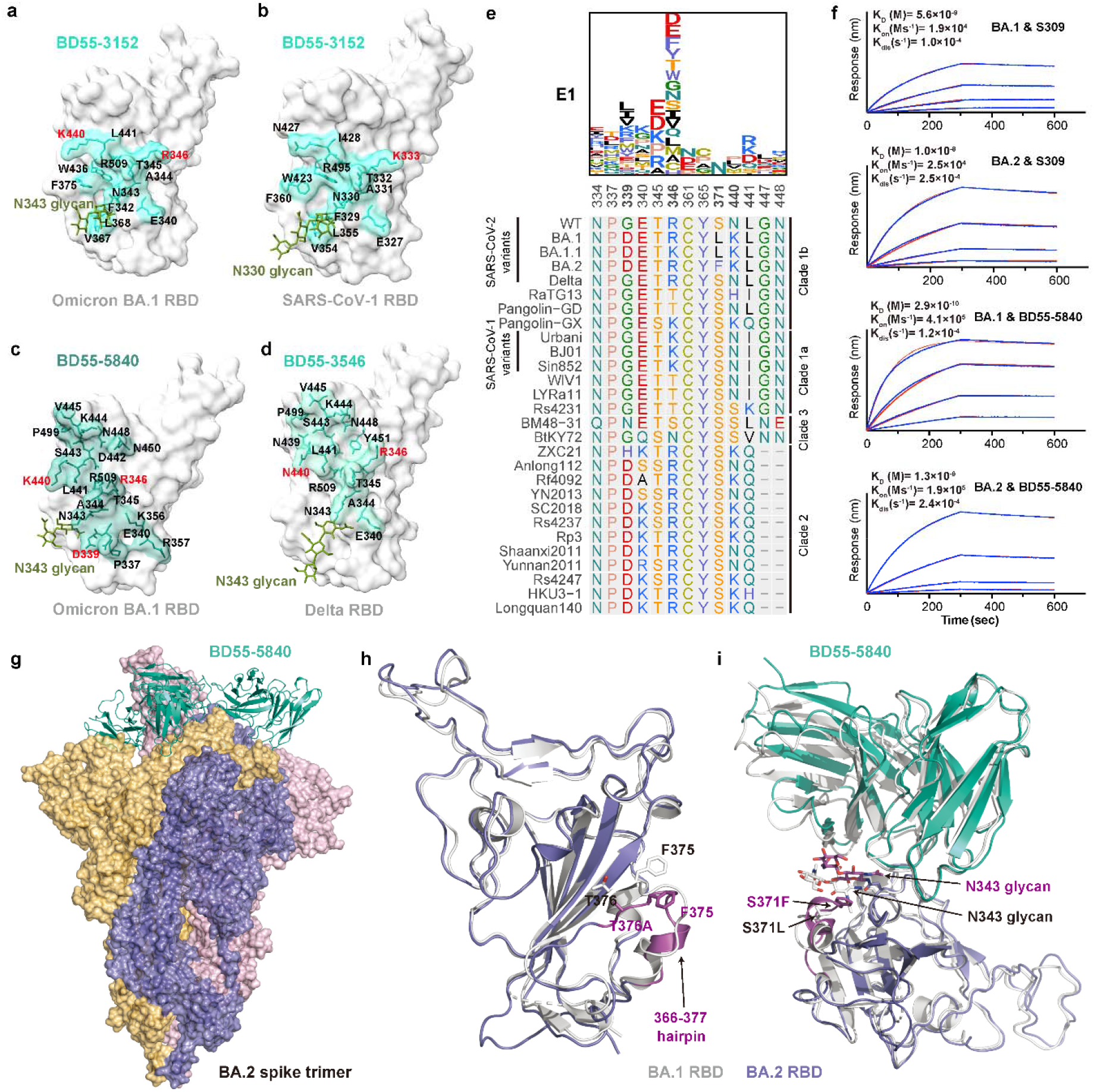
BA.2 reduces E1 antibodies’ potency through S371F-induced glycan moiety hindrance. **a-d,** Epitope of Group E1 broad neutralizing antibodies on SARS-CoV-2 RBD determined by Cryo-EM structures. **a,** BD55-3152 in complex of Omicron BA.1 RBD. **b,** BD55-3152 in complex of SARS-CoV-1 RBD. **c,** BD55-5840 in complex of Omicron BA.1 RBD. **d,** BD55-3546 in complex of Delta RBD. Residues on the binding interface are marked. Residues highlighted in red indicate mutated sites in Omicron variants. **e,** Averaged escape maps at escape hotspots of antibodies in epitope group E1, and corresponding multiple sequence alignment (MSA) of various sarbecovirus RBDs. Height of each amino acid in the escape maps represents its mutation escape score. Mutated sites in Omicron variants are marked in bold. **f,** BLI sensorgrams of Group E1 antibodies S309 and BD55-5840 binding to Omicron BA.1 and BA.2 Spike trimer. **g,** The BA.2 spike trimer (blue) in complex with three BD55-5840 Fabs (cyan). **h,** Structural overlay of the BA.2 (blue) and BA.1 (white) RBDs. The 366-377 hairpin in BA.2 RBD is highlighted in magenta. **i,** Structural overlay of the BA.2 RBD (blue)/BD55-5840 (cyan) and BA.1 RBD (white)/BD55-5840 (white) complexes. The 366-377 hairpin and N343 glycan in BA.2 RBD are highlighted in magenta.

Intriguingly, the newly acquired mutations of BA.2 do not overlap with the shared epitope of E1 antibodies, suggesting that the systematic reduction of neutralization is not caused by amino-acid substitution, but potentially due to structural alteration. To probe the molecular framework of the BA.2 spike, we determined the cryo-EM structure of the prefusion-stabilized BA.2 spike in complex with the BD55-5840 Fab (Fig. 3g). In this structure, the BA.2 spike exhibits a typical one-RBD-up asymmetric conformation, with all three RBDs bound to the BD55-5840 Fabs (Fig. 3g). A structural comparison with the BA.1 RBD bound to BD55-5840 described above suggests that the general structure of BA.2 RBD is similar to that of the BA.1 (Fig. 3h). However, the 366-377 hairpin loop that harbors the S371F and T376A BA.2-specific mutations display significant conformational differences. As a result, the bulky Phe resulting from the S371F mutation interferes with the positioning of the glycan moiety attached to Asn343, which in turn budges the heavy chain of BD55-5840 upward (Fig. 3i). This may explain the reduction of the binding of BD55-5840 and S309, rationalizing their reduced neutralizing activity (Fig. 3f). Importantly, the Asn343 glycan is critically recognized by almost all E1 neutralizing antibodies, including S309. Thus, this group of broad and potent neutralizing antibodies is likely affected by the S371F mutation in a systematic manner through N343-glycan displacement.

### BA.2 escapes neutralizing antibodies through T376A, D405N and R408S

The epitopes of group F2 and F3 antibodies cover a continuous surface on the backside of RBD and can only bind to the up RBDs (Fig. 2b). To probe how BA.2 escapes antibodies of group F2 and F3, we solved the cryo-EM structure of four representative BA.1 effective antibodies in these group, BD55-1239, BD55-3500 from group F2, and BD55-4637, BD55-3372 from group F3, in complex with the spike protein (Fig. 4a,c, Extended Data Fig. 6). Group F2 antibodies can be escaped by RBD mutation involving T376, K378, and R408 (Fig. 3e). Indeed, these residues are all at the heart of BD55-1239’s and BD55-3500’s epitopes. These sites are fairly conserved across sarbecoviruses (Fig. 4e), and the neutralization breadth of group F2 antibodies is extraordinarily broad (Fig. 2c,d). However, group F2 antibodies suffer great neutralization efficacy reduction against Omicron BA.1, mostly due to the triple mutations on S371, S373, S375 (Fig. 2f). Importantly, mutation T376A and R408S harbored by Omicron BA.2 are the major escaping mutations of group F2 antibodies (Fig. 4e). T376A and R408S could alter the antigenic surface that disrupts the binding of both the heavy-chain and light-chain of F2 antibodies (Fig. 4b), hence completely abolishing the neutralizing capacity of F2 antibodies (Fig. 2f).

**Fig. 4.**
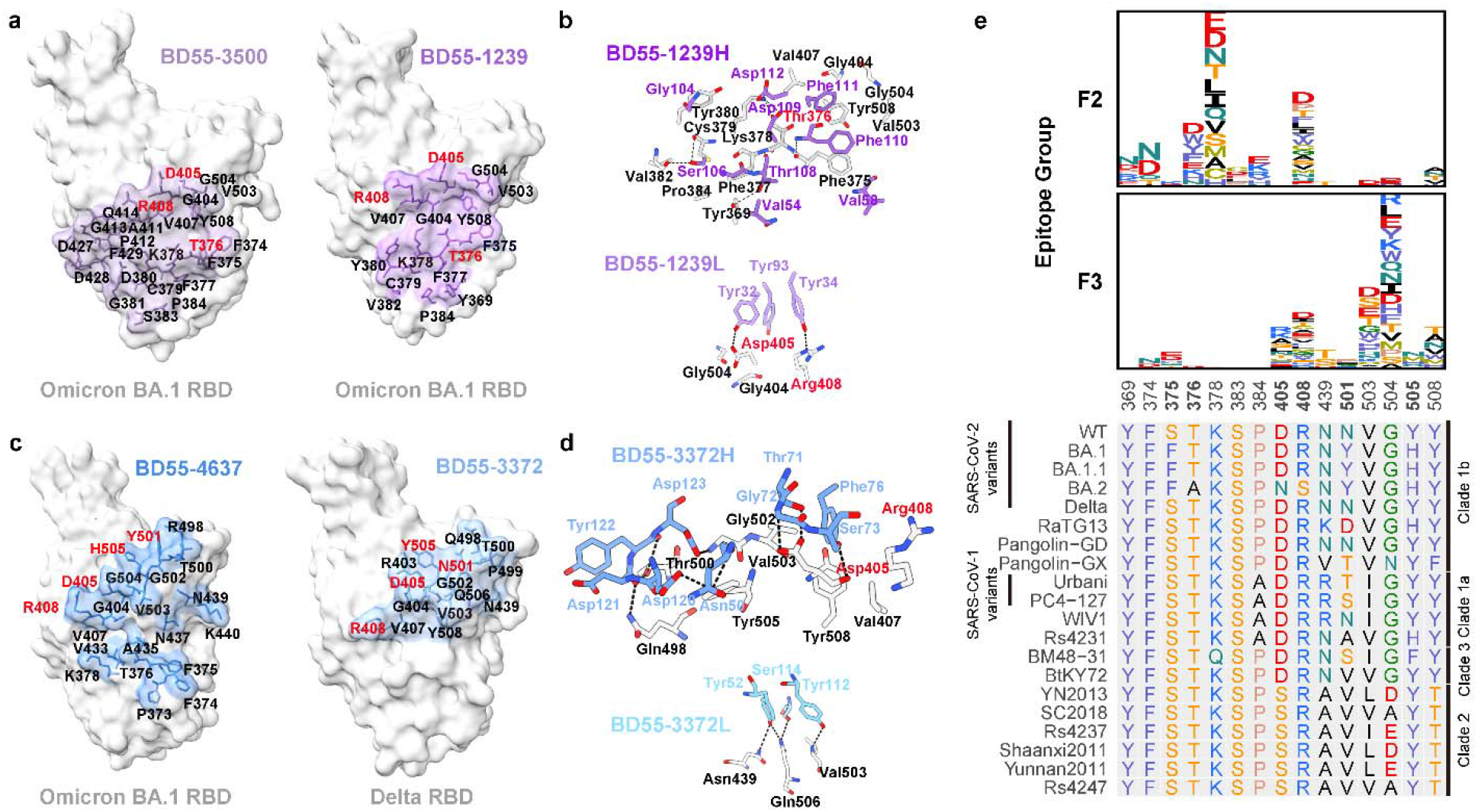
BA.2 escapes BA.1 effective F2 and F3 antibodies through T376A, D405N and R408S. **a,** Epitope of Group F2 antibodies BD55-3500 and BD55-1239 on SARS-CoV-2 Omicron BA.1 RBD. **b,** Interactions of residues on the binding interface of BD55-1239 and BA.1 RBD complex. Residues of the antibody are purple, and RBD residues are black or red. **c,** Epitope of Group F3 antibodies BD55-4637 on Delta RBD and BD55-3372 on BA.1 RBD. **d,** Interactions of residues on the binding interface of BD55-3372 and Delta RBD complex. Residues of the antibody are blue, and RBD residues are black or red. **e,** Averaged escape maps of antibodies in epitope group F2 and F3, and corresponding multiple sequence alignment (MSA) of various sarbecovirus RBDs. Height of each amino acid in the escape maps represents its mutation escape score. Residues are colored corresponding to their chemical properties. Mutated sites in Omicron variants are marked in bold.

Major escape sites for group F3 antibodies include D405, R408, V503, G504, and Y508 (Fig. 4e). Given the fact that D405, G504, and Y508 are not conserved in clade 2 sarbecoviruses, F3 antibodies cannot bind to non-ACE2 utilizing clade 2 sarbecovirus (Fig. 2d), but show good neutralization breadth against clade 1a/b/2 sarbecovirus (Fig. 2c). Similar to group E1, a proportion of F3 antibodies showed potent neutralization against Omicron BA.1, despite the Y505H and N501Y mutations carried by Omicron (Fig. 2f). However, the D405N and R408S mutations harbored by BA.2 could interrupt the heavy-chain binding of F3 antibodies, causing large-scale escapes of BA.1-effective F3 neutralizing antibodies (Fig. 4d). Nevertheless, several members of F3 antibodies are not sensitive to the D405N and R408S mutations of BA.2, making them good therapeutic drug candidates (Extended Data Fig. 7). In sum, we revealed that BA.2 could induce large-scale escapes of broad sarbecovirus neutralizing antibodies through T376A, D405N and R408S mutations.

In this study, we reported the discovery of five broad sarbecovirus neutralizing antibody groups, where BA.2 could cause large-scale escapes or neutralization reduction of the top three most potent groups, E1, F2 and F3. The remaining F1 group, which contains CR3022^21^ and S304^22^, and E3 group, which includes S2H97^20^, mainly target the RBD regions that are deeply buried within the spike trimer (Extended Data Fig. 8b-c), and are generally weak in neutralization potency (Extended Data Fig. 5). The major escaping mutation sites of E1 and F3 are highly conserved across sarbecovirus (Extended Data Fig. 8d); hence members of these groups usually display great neutralization breadth (Fig. 2d). However, since group F1 and E3 antibodies achieve neutralization capability through the disruption of spike trimer’s prefusion state, they demand the extensive opening of the RBD to engage their respective binding sites^20,44–46^. Thus, F1 and E3 virtually have no neutralization power against Omicron variants (Fig. 2c), which should be because that the Omicron spike displays a more stabilized prefusion structure compared to wildtype^47^. Notably, a unique subcluster of Group B antibodies also showed broadspectrum sarbecovirus neutralizing capability (Fig. 2c). Most Group B antibodies are SARS-CoV-2 specific since their major escaping mutations consist of E484 and F486, which are not conserved in sarbecovirus clade (Supplementary Data 3); however, the rare sub-cluster B’, featured by S2K146^39^, displayed skewed escaping mutation profiles toward N487 and Y489, which are highly conserved in clade 1a/1b/3 (Extended Data Fig. 7), making members of B’ exhibit similar breadth as F3 antibodies. Sadly, most B’ antibodies failed to neutralize Omicron variants, except for S2K146^6^ (Supplementary Table 2).

Together, we showed that BA.2 exhibits distinct antigenicity compared to BA.1, where BA.2 evolved multiple mutations on the sarbecovirus conserved regions (S371F, D405N, R408S and T376A) that could cause pinpoint escapes of broad sarbecovirus neutralizing antibodies. This supports the speculation that Omicron does not originate from zoonotic spillover but immune selection pressure. Nevertheless, BA.2 lacks the G446S mutation, and thus part of group D antibodies that target the linear 440-449 loop retained their neutralization capability against BA.2, which compensates for the loss of broad sarbecovirus neutralizing antibodies. Concerningly, BA.3 harbors G446S, S371F and R405N (Extended Data Fig. 1), making it potentially the heaviest antibody evading SARS-CoV-2 variants to date. In sum, our results provide a comprehensive profile of SARS-CoV-2 antibody escaping mutations and offer critical insights into the humoral immune evading mechanism of current and future variants, which is crucial to sarbecovirus antibody therapeutics development and broad-spectrum vaccines design.

## Supporting information

Supplementary Guide

## Acknowledgments

We thank J. Bloom for his generous gift of the yeast SARS-CoV-2 RBD libraries. We thank Sino Biological for the technical assistance on mAbs and RBD expression. We thank J. Luo and H. Lv for the help in flow cytometry. This project is financially supported by the Ministry of Science and Technology of China (CPL-1233).

## Author contributions

Y.C. and X.S.X designed the study. Y.C., J.X., X.S.X wrote the manuscript with inputs from all authors. Y.C. and F.S. coordinated the expression and characterization of the neutralizing antibodies. J.W. (BIOPIC), F.J., L.Z., H.S. performed and analyzed the yeast display screening experiments. T.X., P.W., J.W. (Changping Laboratory), R.A., Y.W., J.Z., N.Z., R.W., X.N., L.Y., C.L., X. S. L.Z., F.S. performed the neutralizing antibody expression and characterization, including pseudovirus neutralization, authentic virus neutralization, BLI, and ELISA. W.H., Q.L., Y.W. prepared the VSV-based SARS-CoV-2 pseudovirus. A.Y., Y.W., S.Y., R.A., W.S. performed and analyzed the antigen-specific single B cell VDJ sequencing. S.D., P.L., Z.Z., L.W., P.Y., X.W., J.X. performed the antibody structural analyses. X.H., W.Z., D.Z., and R.J. recruited the SARS convalescents and SARS-CoV-2 vaccinee. X.C. and Z.S. recruited the Omicron BA.1 convalescents. Q.G. proofed the manuscript.

## Data availability

Logo plots of escape maps for antibodies in this study are available in Supplementary Data 1. Processed mutation escape scores can be downloaded at https://github.com/jianfcpku/SARS-CoV-2-RBD-DMS-broad. Raw Illumina and PacBio sequencing data are available on NCBI Sequence Read Archive BioProject PRJNA787091. We used vdj_GRCh38_alts_ensembl-5.0.0 as the reference of V(D)J alignment, which can be obtained from https://support.10xgenomics.com/single-cell-vdj/software/downloads/latest.

IMGT/DomainGapAlign is based on the built-in lastest IMGT antibody database, and we let the “Species” parameter as “Homo sapiens” while kept the others as default. Public deep mutational scanning datasets involved in the study from literature could be downloaded at https://media.githubusercontent.com/media/jbloomlab/SARS2_RBD_Ab_escape_maps/main/processed_data/escape_data.csv.

Cryo-EM density maps have been deposited in the Electron Microscopy Data Bank with accession codes EMD-32732, EMD-32737, EMD-32738, EMD-32753, EMD-32734, EMD-32728, EMD-32718, EMD-32719, and EMD-33019, respectively. Structural coordinates have been deposited in the Protein Data Bank with accession codes 7WRL, 7WRY, 7WRZ, 7WSC, 7WRO, 7WRJ, 7WR8, 7WR9, and 7X6A.

## Code availability

Codes for analyzing SARS-CoV-2 escaping mutation profile data are available at https://github.com/sunneyxielab/SARS-CoV-2-RBD-Abs-HTDMS. R and Python scripts for reproducing figures in this manuscript are available at https://github.com/jianfcpku/SARS-CoV-2-RBD-DMS-broad.

## Materials and Methods

### Plasma and PBMC isolation

Blood samples were obtained from 40 volunteers who received 3 doses of CoronaVac, 39 individuals who received 2 doses of CoronaVac and 1 booster dose of ZF2001, 54 BA.1 convalescents who had received 3 doses of CoronaVac before BA.1 infection, and 30 SARS convalescents who received 2 doses of CoronaVac and 1 dose of ZF2001. The volunteers’ blood samples were obtained 4 weeks after the booster shot or 4 weeks after discharge from the hospital after BA.1 infection. Relevant experiments regarding SARS convalescents and SARS-CoV-2 vaccinees were approved by the Beijing Ditan Hospital Capital Medical University (Ethics committee archiving No. LL-2021-024-02). Written informed consent was obtained from each participant in accordance with the Declaration of Helsinki. All participants provided written informed consent for the collection of information, and that their clinical samples were stored and used for research. Data generated from the research were agreed to be published.

Whole blood samples were mixed and subjected to Ficoll (Cytiva, 17-1440-03) gradient centrifugation after 1:1 dilution in PBS+2% FBS to isolate plasma and peripheral blood mononuclear cells (PBMC). After centrifugation, plasma was collected from upper layer and cells were harvested at the interface, respectively. PBMCs were further prepared through centrifugation, red blood cells lysis (InvitrogenTM eBioscienceTM 1X RBC Lysis Buffer, 004333-57) and washing steps. Samples were stored in FBS (Gibco) with 10% DMSO (Sigma) in liquid nitrogen if not used for downstream process immediately. Cryopreserved PBMCs were thawed in DPBS+2% FBS (Stemcell, 07905).

### Antibody isolation and recombinant production

SARS-CoV-1 and SARS-CoV-2 RBD cross-binding memory B cells were isolated from PBMC of SARS-CoV-1 convalescents who received SARS-CoV-2 vaccine. Briefly, CD19+ B cells were isolated from PBMC with EasySep™ Human CD19 Positive Selection Kit II (STEMCELL, 17854). B cells were then stained with FITC anti-human CD19 antibody (BioLegend, 392508), FITC anti-human CD20 antibody (BioLegend, 302304), Brilliant Violet 421™ anti-human CD27 antibody (BioLegend, 302824), PE/Cyanine7 anti-human IgM antibody (BioLegend, 314532), biotinylated Ovalbumin (SinoBiological) conjugated with Brilliant Violet 605™ Streptavidin (BioLegend, 405229), SARS-CoV-1 biotinylated RBD protein (His & AVI Tag) (SinoBiological, 40634-V27H-B) conjugated with PE-streptavidin (BioLegend, 405204), SARS-CoV-2 biotinylated RBD protein (His & AVI Tag) (SinoBiological, 40592-V27H-B) conjugated with APC-streptavidin (BioLegend, 405207), and 7-AAD (Invitrogen, 00-6993-50). CD19/CD20+, CD27+, IgM-, OVA-, SARS-COV-1 RBD+, and SARS-CoV-2 RBD+ were sorted with MoFlo Astrios EQ Cell Sorter (Beckman Coulter). FACS data were analyzed using FlowJo™ v10.8 (BD Biosciences).

Sorted SARS-CoV-1 and SARS-CoV-2 RBD cross-binding B cells were then processed with Chromium Next GEM Single Cell V(D)J Reagent Kits v1.1 following the manufacturer’s user guide (10x Genomics, CG000208). Briefly, Cells sorted were resuspend in PBS after centrifugation. Gel beads-in-emulsion (GEMs) were obtained with 10X Chromium controller and then subjected to reverse transcription (RT). After GEM-RT clean up, RT products were subject to preamplification. After amplification and purification with SPRIselect Reagent Kit (Beckman Coulter, B23318) of RT products, B cell receptor (BCR) sequence (paired V(D)J) were enriched with 10X BCR primers. After library preparation, libraries were sequenced by Novaseq 6000 platform running Novaseq 6000 S4 Reagent Kit v1.5 300 cycles (Illumina, 20028312) or NovaSeq XP 4-Lane Kit v1.5 (Illumina, 20043131).

After sequencing and data processing, monoclonal antibodies were expressed as recombinant human IgG1. Briefly, HEK293F cells (Thermo Fisher, R79007) were transiently transfected with heavy and light chain expression vectors. The secreted monoclonal antibodies from cultured cells were purified by protein A affinity chromatography. The specificities of these antibodies were determined by ELISA.

### Antibody sequence analysis

The antibody sequences obtained from 10X Genomics V(D)J sequencing were aligned to GRCh38 reference and assembled as immunoglobulin contigs by the Cell Ranger (v6.1.1) pipeline. Non-productive contigs and B cells that had multiple heavy chain or light chain contigs were filtered out of the analysis. V(D)J gene annotation was performed using NCBI IgBlast (v1.17.1) with the IMGT reference. Mutations on V(D)J nucleotide sequences were calculated by using the igpipeline, which compared the sequences to the closest germline genes and counted the number of different nucleotides. For antibodies from public sources whose original sequencing nucleotide sequences were not all accessible, the antibody amino acid sequences were annotated by IMGT/DomainGapAlign^49^ (v4.10.2) with default parameters. The V-J pairs were visualized by R package circlize (v0.4.10).

### High-throughput antibody-escape mutation profiling

The previously described high-throughput MACS (magnetic-activated cell sorting)-based antibody-escape mutation profiling system was used to characterize RBD escaping mutation profile for neutralizing antibodies. Briefly, duplicate RBD mutant libraries were constructed based on Wuhan-Hu-1 RBD sequence (GenBank: MN908947, residues N331-T531), theoretically containing 3819 possible amino acid mutations. Each RBD mutant was barcoded with a unique 26-neuclotide (N26) and only ACE2 binding variants were enriched for downstream experiment.

For antibody escape profiling, yeast libraries were induced overnight for RBD expression and washed followed by two rounds of Protein A antibody based negative selection and MYC-tag based positive selection to enrich RBD expressing cells. Protein A antibody conjugated products were prepared following the protocol of Dynabeads Protein A (Thermo Fisher, 10008D) and incubated with induced yeast libraries at room temperature. MYC-tag based positive selection was performed according to the manufacturer’s protocol (Thermo Fisher, 88843).

After three rounds of sequential cell sorting, the obtained yeast cells were recovered overnight. Plasmids were extracted from pre- and post-sort yeast populations by 96-Well Plate Yeast Plasmid Preps Kit (Coolaber, PE053). The extracted plasmids were then used to amplify N26 barcode sequences by PCR. The final PCR products were purified with AMPure XP magnetic beads (Beckman Coulter, A63882) and 75bp single-end sequencing was performed on an Illumina Nextseq 500 platform.

### Processing of deep mutational scanning data

Single-end Illumina sequencing reads were processed as previously described. Briefly, reads were trimmed into 16 or 26 bp and aligned to the reference barcode-variant dictionary with dms_variants package (v0.8.9). Escape scores of variants were calculated as F×(n_X,ab_ / N_ab_) / (n_X,ref_ / N_ref_), where n_X,ab_ and n_X,ref_ is the number of reads representing variant X, and N_ab_ and N_ref_ are the total number of valid reads in antibody-selected (ab) and reference (ref) library, respectively. F is a scale factor defined as the 99th percentiles of escape fraction ratios. Variants detected by less than 6 reads in the reference library were removed to avoid sampling noise. Variants containing mutations with ACE2 binding below −2.35 or RBD expression below −1 were removed as well, according to data previously reported. Finally, global epistasis models were built using dms_variants package to estimate mutation escape scores. For most antibodies, at least two independent assays are conducted and single mutation escape scores are averaged across all experiments that pass quality control.

### Antibody clustering and visualization

Site total escape scores, defined as the sum of escape scores of all mutations at a particular site on RBD, were used to evaluate the impact of mutations on each site for each antibody. Each of these scores is considered as a feature of a certain antibody and used to construct a feature matrix **A**_N_×_M_ for downstream analysis, where N is the number of antibodies and M is the number of features (valid sites). Informative sites were selected using sklearn.feature_selection.VarianceThreshold of scikit-learn Python package (v0.24.2) with the variance threshold as 0.1. Then, the selected features were L2-normalized across antibodies using sklearn.preprocessing.normalize. The resulting matrix is referred as **A’N×M’**, where M’ is the number of selected features. The dissimilarity of two antibodies i, j is defined as 1-Corr(**A’**_i_,**A’**_j_), where Corr(**x**,**y**) is the Pearson’s correlation coefficient of vector **x** and **y**, i.e. Corr(**x**,**y**)= **x**·**y**/|**x**||**y**|. We used sklearn.manifold.MDS to reduce the number of features from M’ to D=20 with multidimensional scaling under the above metric. Antibodies are clustered into 10 epitope groups using sklearn.cluster.KMeans of scikit-learn in the resulting D-dimensional feature space. Finally, these D-dimensional representations of antibodies were further embedded into two-dimensional space for visualization with t-SNE using sklearn.manifold.TSNE of scikit-learn. All t-SNE plots were generated by R package ggplot2 (v3.3.3).

### Pseudovirus neutralization assay

SARS-CoV-2 spike (GenBank: MN908947), Pangolin-GD spike (GISAID: EPI_ISL_410721), RaTG13 spike (GISAID: EPI_ISL_402131), SARS-COV-1 spike (GenBank: AY278491), Omicron BA.1 spike (A67V, H69del, V70del, T95I, G142D, V143del, Y144del, Y145del, N211del, L212I, ins214EPE, G339D, S371L, S373P, S375F, K417N, N440K, G446S, S477N, T478K, E484A, Q493R, G496S, Q498R, N501Y, Y505H, T547K, D614G, H655Y, N679K, P681H, N764K, D796Y, N856K, Q954H, N969K, L981F), BA.2 spike (GISAID: EPI_ISL_7580387, T19I, L24S, del25-27, G142D, V213G, G339D, S371F, S373P, S375F, T376A, D405N, R408S, K417N, N440K, G446S, S477N, T478K, E484A, Q493R, Q498R, N501Y, Y505H, D614G, H655Y, N679K, P681H, N764K, D796Y, Q954H, N969K), BA.1.1 spike (BA.1+R346K), plasmid is constructed into pcDNA3.1 vector. G*ΔG-VSV virus (VSV G pseudotyped virus, Kerafast) is used to infect 293T cells (American Type Culture Collection [ATCC], CRL-3216), and spike protein expressing plasmid was used for transfection at the same time. After culture, the supernatant containing pseudovirus was harvested, filtered, aliquoted, and frozen at −80°C for further use.

Pseudovirus detection of PCoV-GD and RaTG13 was performed in 293T cells overexpressing human angiotensin-converting enzyme 2 (293T-hACE2 cells). Other pseudovirus neutralization assays were performed using the Huh-7 cell line (Japanese Collection of Research Bioresources [JCRB], 0403).

Monoclonal antibodies or plasma were serially diluted (5-fold or 3-fold) in DMEM (Hyclone, SH30243.01) and mixed with pseudovirus in 96-well plates. After incubation at 5% CO_2_ and 37□ for 1 h, digested Huh-7 cell (Japanese Collection of Research Bioresources [JCRB], 0403) or 293T-hACE2 cells (AmericanTypeCultureCollection[ATCC],CRL-3216) were seeded. After 24 hours of culture, supernatant was discarded and D-luciferin reagent (PerkinElmer, 6066769) was added to react in the dark, and the luminescence value was detected using a microplate spectrophotometer (PerkinElmer, HH3400). IC50 was determined by a four-parameter logistic regression model.

### ELISA

To detect the broad-spectrum binding of the antibodies among Sarbecovirus, we entrusted SinoBiological Technology Co., Ltd. to synthesize a panel of 20 sarbecovirus RBDs (Supplementary Table 3). According to the sequence of 20 RBDs, a set of nested primers was designed. The coding sequences were obtained by the overlap-PCR with a 6xHis tag sequence to facilitate protein purification. The purified PCR products were ligated to the secretory expression vector pCMV3 with CMV promoter, and then transformed into E. coli competent cells XL1-blue. Monoclones with correct transformation were cultured and expanded, and plasmids were extracted. Healthy HEK293 cells were passaged into a new cell culture and grown in suspension at 37 °C, 120 RPM, 8% CO2 to logarithmic growth phase and transfected with the recombinant constructs by using liposomal vesicles as DNA carrier. After transfection, the cell cultures were followed to assess the kinetics of cell growth and viability for 7 days. The cell expression supernatant was collected, and after centrifugation, passed through a Ni column for affinity purification. The molecular size and purity of eluted protein was confirmed by SDS-PAGE. Production lot numbers and concentration information of the 20 Sarbecovirus proteins are shown in Suppplemenatary Table 4. The WT RBD in the article is SARS-CoV-2 (2019-nCoV) Spike RBD-His Recombinant Protein (SinoBiological, 40592-V08H).

A panel of 21 sarbecovirus RBDs (supplementary table3) in PBS was pre-coated onto ELISA Plates (NEST, 514201) at 4□ overnight. The plates were washed and blocked. Then 1μg/ml purified antibodies or serially diluted antibodies were added and incubated at room temperature (RT) for 20min. Next, Peroxidase-conjugated AffiniPure Goat Anti-Human IgG (H+L) (JACKSON, 109-035-003) was applied and incubated at RT for 15min. Tetramethylbenzidine (TMB) (Solarbio, 54827-17-7) was added onto the plates. The reaction was terminated by 2 M H_2_SO_4_ after 10min incubation. Absorbance was measured at 450 nm using Ensight Multimode Plate Reader (PerkinElmer, HH3400). ELISA OD450 measurements at different antibody concentration for a particular antibody-antigen pair are fit to the model y=Ac^n^/(c^n^ + E^n^) using R package mosaic (v1.8.3), where y is OD450 values and c is corresponding antibody concentration. A, E, n are parameters, where E is the desired EC_50_ value for the specific antibody and antigen.

### Antibody-ACE2 competition for RBD

mFC-WT-RBD (Sino Biological, 40592-V05H) protein in PBS was immobilized on the ELISA plates at 4□ overnight. The coating solution was removed and washed three times by PBST and the plates were then blocked for 2 h. After blocking, the plates were washed five times, and the mixture of ACE2-his (Sino Biological, 10108-105H) and serially diluted competitor antibodies was added followed by 30min incubation at RT. Then anti-his-HRP (Proteintech, HRP-66005) was added into each well for another 20min incubation at RT. After washing the plates for five times, Tetramethylbenzidine (TMB) (Solarbio, 54827-17-7) was added into each well. After 10 min, the reaction was terminated by 2M H2SO4. Absorbance was measured at 450 nm using Ensight Multimode Plate Reader ( PerkinElmer, HH3400). The ACE2 competition coefficient is calculated as (B-A)/B, where B is the OD450 value under 0.15ug/ml antibody concentration and A is the OD450 value under 3ug/ml antibody concentration.

### Biolayer Interferometry

Biolayer interferometry assays were performed on Octet^®^ RED 384 Protein Analysis System (Fortebio) according to the manufacturer’s instruction. To measure the binding affinities, monoclonal antibodies were immobilized onto Protein A biosensors (Fortebio) and the fourfold serial dilutions of Omicron S trimer (BA.1 and BA.2) in PBS were used as analytes. Data were collected with Octet Acquisition 9.0 (Fortebio) and analyzed by Octet Analysis 9.0 (Fortebio) and Octet Analysis Studio 12.2 (Fortebio).

### Protein expression and purification for cryo-EM study

The S6P expression construct encoding the SARS-CoV-2 spike ectodomain (residues 1-1208) with six stabilizing Pro substitutions (F817P, A892P, A899P, A942P, K986P, and V987P) and a “GSAS” substitution for the furin cleavage site (residues 682–685) was previously described^22^. The Delta specific mutations (T19R, G142D, 156del, 157del, R158G, L452R, T478K, D614G, P681R, D950N) were introduced into this construct using site-directed mutagenesis. The S6P expression construct containing the Omicron BA.1 mutations (A67V, H69del, V70del, T95I, G142D, V143del, Y144del, Y145del, N211del, L212I, ins214EPE, G339D, S371L, S373P, S375F, K417N, N440K, G446S, S477N, T478K, E484A, Q493R, G496S, Q498R, N501Y, Y505H, T547K, D614G, H655Y, N679K, P681H, N764K, D796Y, N856K, Q954H, N969K, L981F) were assembled from three synthesized DNA fragments. The S6P expression construct containing the Omicron BA.2 mutations (T19I, L24S, del25-27, G142D, V213G, G339D, S371F, S373P, S375F, T376A, D405N, R408S, K417N, N440K, G446S, S477N, T478K, E484A, Q493R, Q498R, N501Y, Y505H, D614G, H655Y, N679K, P681H, N764K, D796Y, Q954H, N969K) were assembled from three synthesized DNA fragments. The expression constructs encoding the SARS-CoV spike ectodomain (residues 1-1195)^50^ was kindly provided by Prof. X. Wang (Tsinghua University), and two stabilizing Pro substitutions (K968P, V969P) was engineered into this construct using mutagenesis. For protein production, these expression plasmids, as well as the plasmids encoding the antigen-binding fragments (Fabs) of the antibodies described in this paper, were transfected into the HEK293F cells using polyethylenimine (Polysciences). The conditioned media were harvested and concentrated using a Hydrosart ultrafilter (Sartorius), and exchanged into the binding buffer (25 mM Tris, pH 8.0, and 200 mM NaCl). Protein purifications were performed using the Ni-NTA affinity method, followed by gel filtration chromatographies using either a Superose 6 increase column (for the spike proteins) or a Superose 200 increase column (for the Fabs). The final buffer used for all proteins is 20 mM HEPES, pH 7.2, and 150 mM NaCl.

### Cryo-EM data collection, processing, and structure building

The samples for cryo-EM study were prepared essentially as previously described^22,51^ ( Supplementary Table 4, Extended Data Fig. 9, and Extended Data Fig. 10). All EM grids were evacuated for 2 min and glow-discharged for 30 s using a plasma cleaner (Harrick PDC-32G-2). Four microliters of spike protein (0.8 mg/mL) was mixed with the same volume of Fabs (1 mg/mL each), and the mixture was immediately applied to glow-discharged holy-carbon gold grids (Quantifoil, R1.2/1.3) in an FEI Vitrobot IV (4 °C and 100% humidity). Data collection was performed using either a Titan Krios G3 equipped with a K3 direct detection camera, or a Titan Krios G2 with a K2 camera, both operating at 300 kV. Data processing was carried out using cryoSPARC^52^. After 2D classification, particles with good qualities were selected for global 3D reconstruction and then subjected to homogeneous refinement. To improve the density surrounding the RBD-Fab region, UCSF Chimera^53^ and Relion^54^ were used to generate the masks, and local refinement was then performed using cryoSPARC. Coot^55^ and Phenix^56^ were used for structural modeling and refinement. Figures were prepared using USCF ChimeraX^57^ and Pymol (Schrödinger, LLC.).

**Extended Data Fig. 1.**
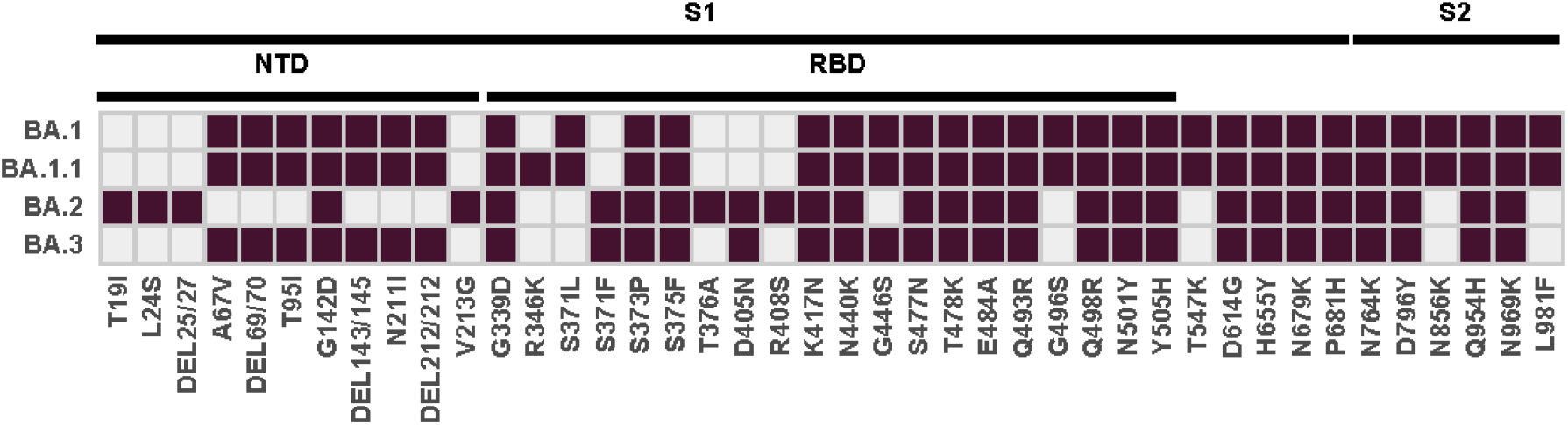
Spike protein mutation differences of Omicron variants. **a,** Mutations on the spike glycoprotein of SARS-CoV-2 Omicron BA.1, BA.1.1, BA.2 and BA.3. Grids are painted in a dark color if >50% of the corresponding sublineage sequences have the corresponding mutation.

**Extended Data Fig. 2.**
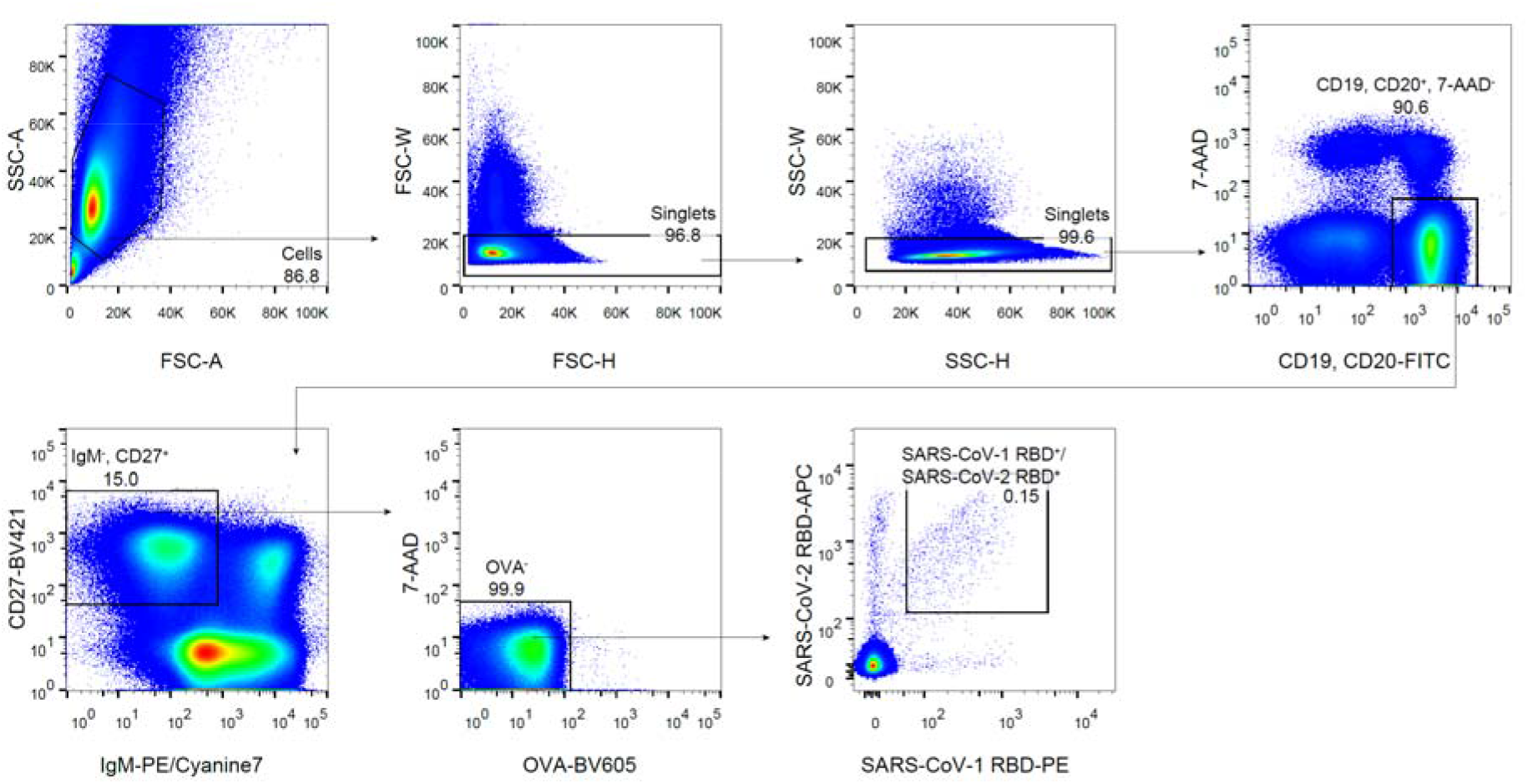
FACS of SARS-CoV-1 RBD and SARS-CoV-2 RBD cross-binding B cells. The gating strategy for sorting SARS-CoV-1-RBD^+^/SARS-CoV-2-RBD^+^ single memory B cells. Numbers next to outlined areas indicate the percentage of cells in the gate. Sorting of the PBMCs from SARS convalescents that received 3 doses of the SARS-CoV-2 vaccine is shown.

**Extended Data Fig. 3.**
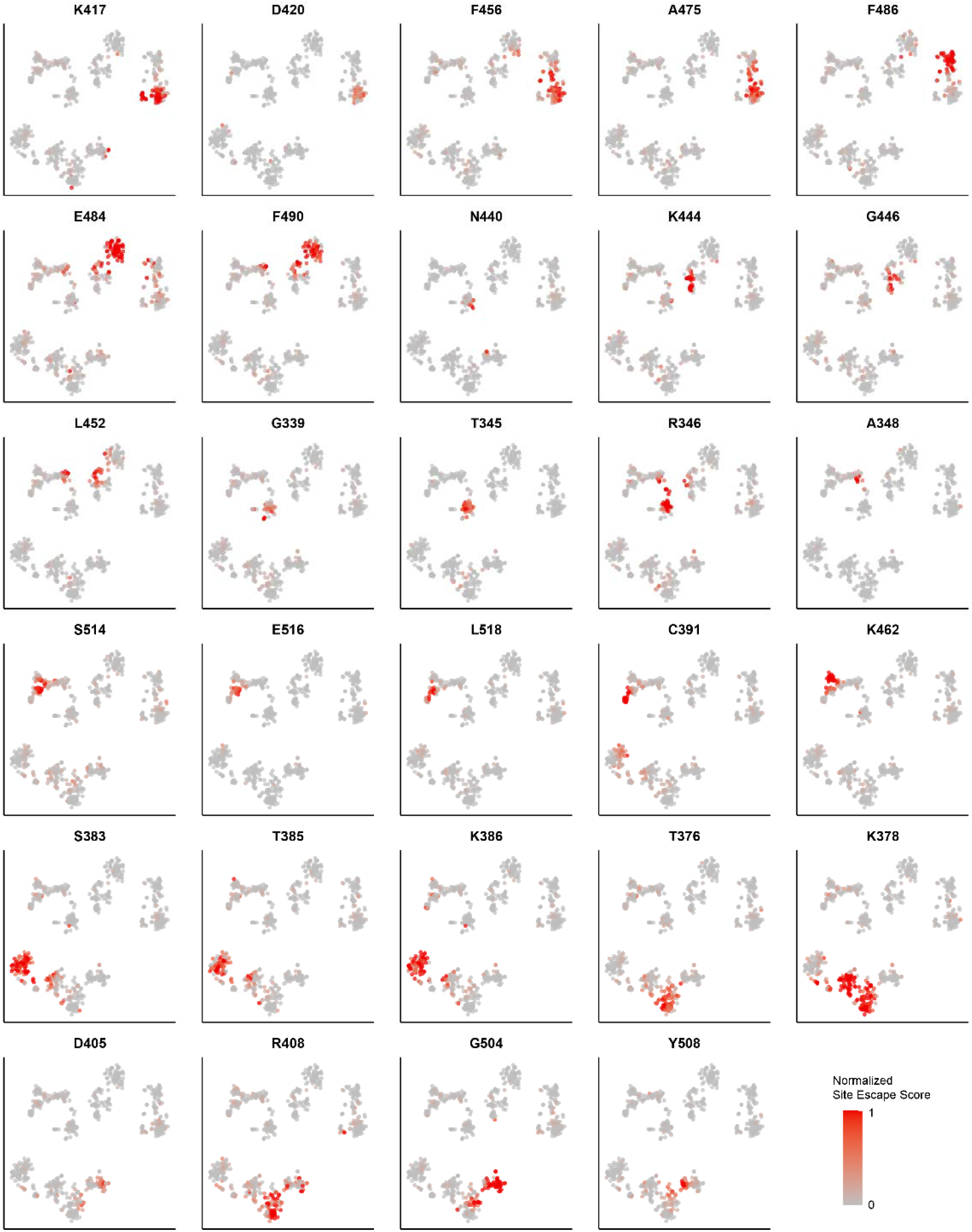
Escape scores projection of top RBD escaping hotspots. Shades of red indicate normalized site total escape scores of the representative residues for each antibody inferred from yeast display deep mutational screening.

**Extended Data Fig. 4.**
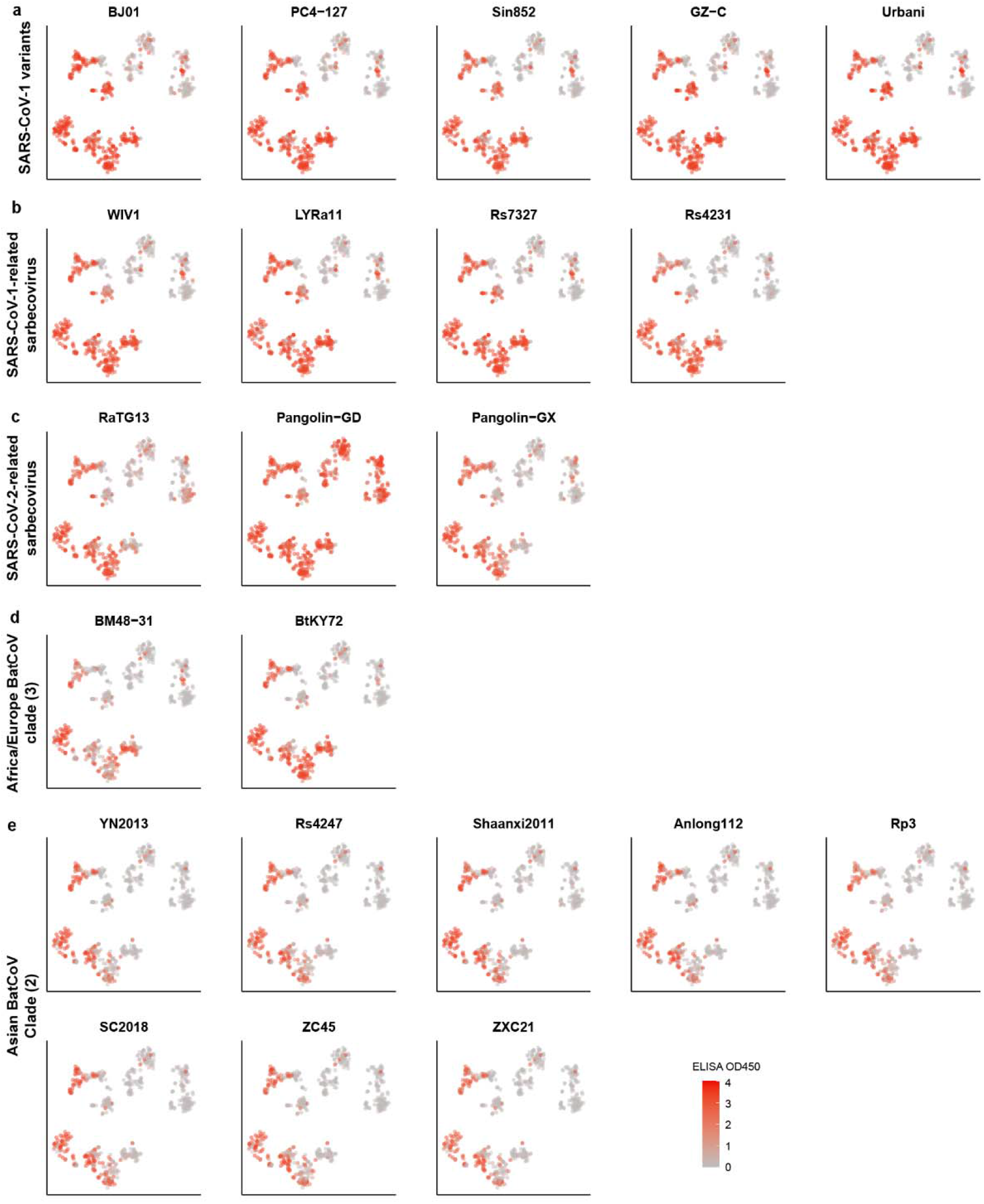
Projection of ELISA reactivity against 22 sarbecovirus RBD. **a-e**, Shades of red indicate ELISA OD450 for each antibody against various sarbecovirus clades. **a**, SARS-CoV-1 variants. **b**, SARS-CoV-1-related sarbecovirus. **c**, SARS-CoV-2 related sarbecovirus. **d**, Africa/Europe batcoronavirus. **e**, Asian non-ACE2-utilizing batcoronavirus.

**Extended Data Fig. 5.**
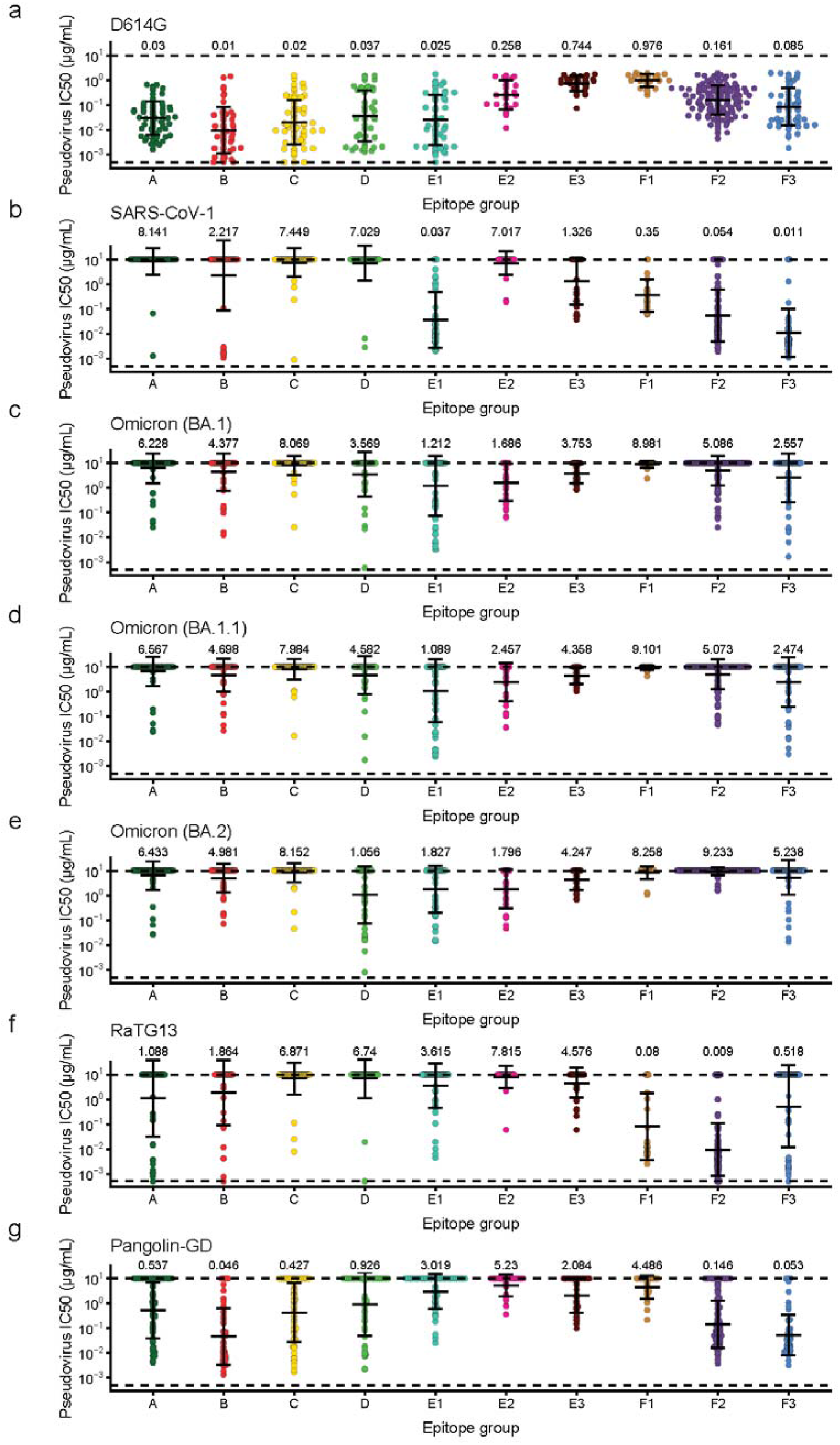
Neutralization potency for neutralizing antibodies of each epitope group. **a-g**, Half maximal inhibitory concentration (IC50) of antibodies against different sarbecovirus determined by pseudovirus neutralization assay. The IC50 values are shown as geometric mean ± s. d. in log10 scale. Dashed lines show the detection limit, which is from 0.0005 μg/mL to 10μg/mL. Number of antibodies n = 68, 49, 61, 45, 53, 27, 31, 23, 126, 57 for epitope group A, B, C, D, E1, E2, E3, F1, F2, F3, respectively (except for **f**). **a**, SARS-CoV-2 with D614G mutation. **b**, SARS-CoV-1 (HKU-39849). **c**, SARS-CoV-2 Omicron (BA.1). **d**, SARS-CoV-2 Omicron (BA.1.1). **e**, SARS-CoV-2 Omicron (BA.2). **f**, RaTG13, n = 50, 38, 47, 41, 52, 27, 31, 23, 125, 56 for epitope group A, B, C, D, E1, E2, E3, F1, F2, F3, respectively. **g**, Pangolin-GD.

**Extended Data Fig. 6.**
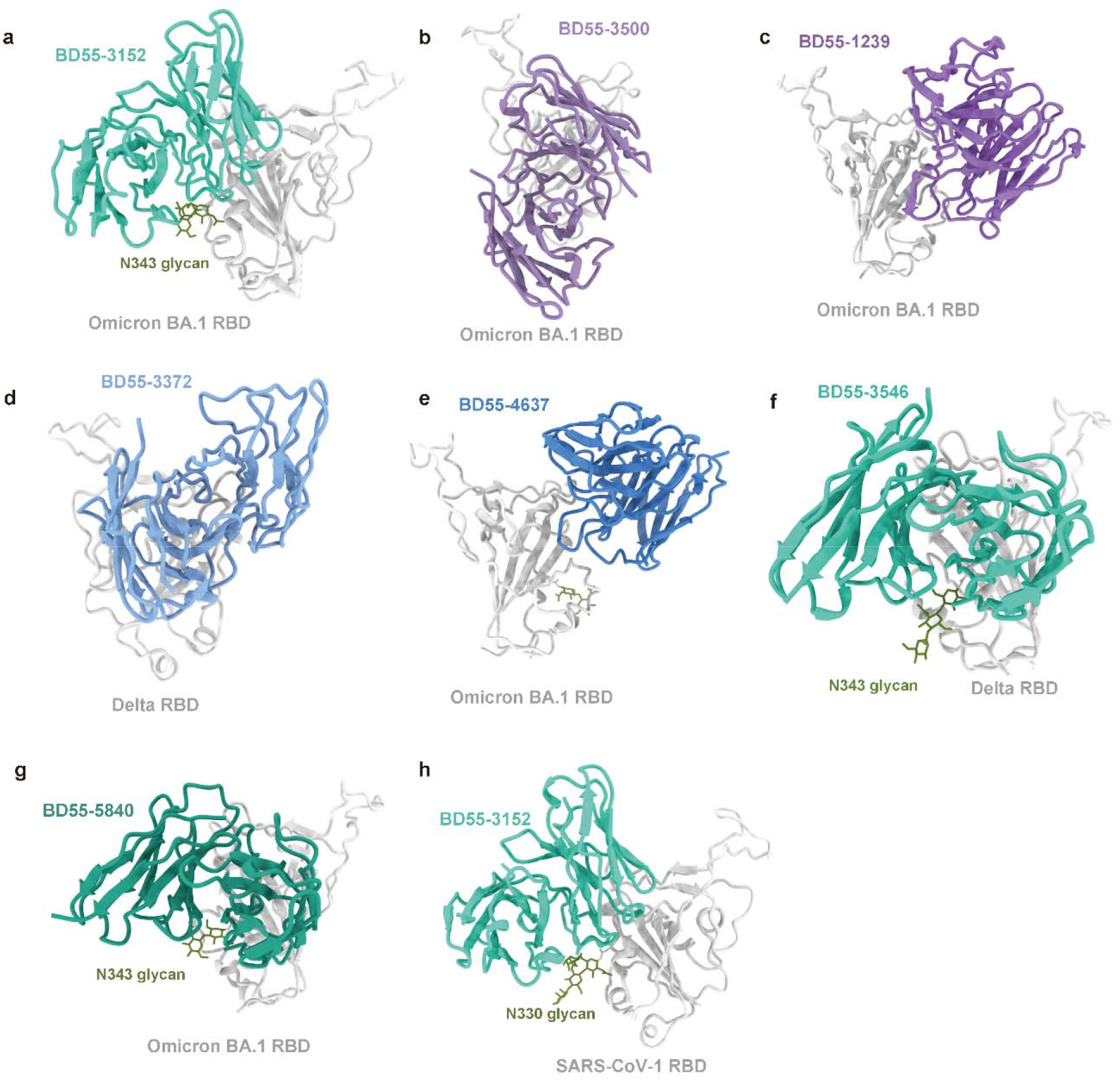
Cryo-EM structures of broad sarbecovirus neutralizing antibodies in complex of RBD. **a,** BD55-3152 (Group E1) in complex of Omicron BA.1 RBD. **b,** BD55-3500 (Group F2) in complex of Omicron BA.1 RBD. **c,** BD55-1239 (Group F2) in complex of Omicron BA.1 RBD. **d,** BD55-3372 (Group F3) in complex of Delta RBD. **e,** BD55-4637 (Group F3) in complex of Omicron BA.1 RBD. **f,** BD55-3546 (Group E1) in complex of Delta RBD. **g,** BD55-5840 (Group E1) in complex of Omicron BA.1 RBD. **h,** BD55-3152 (Group E1) in complex of SARS-CoV-1 RBD.

**Extended Data Fig. 7.**
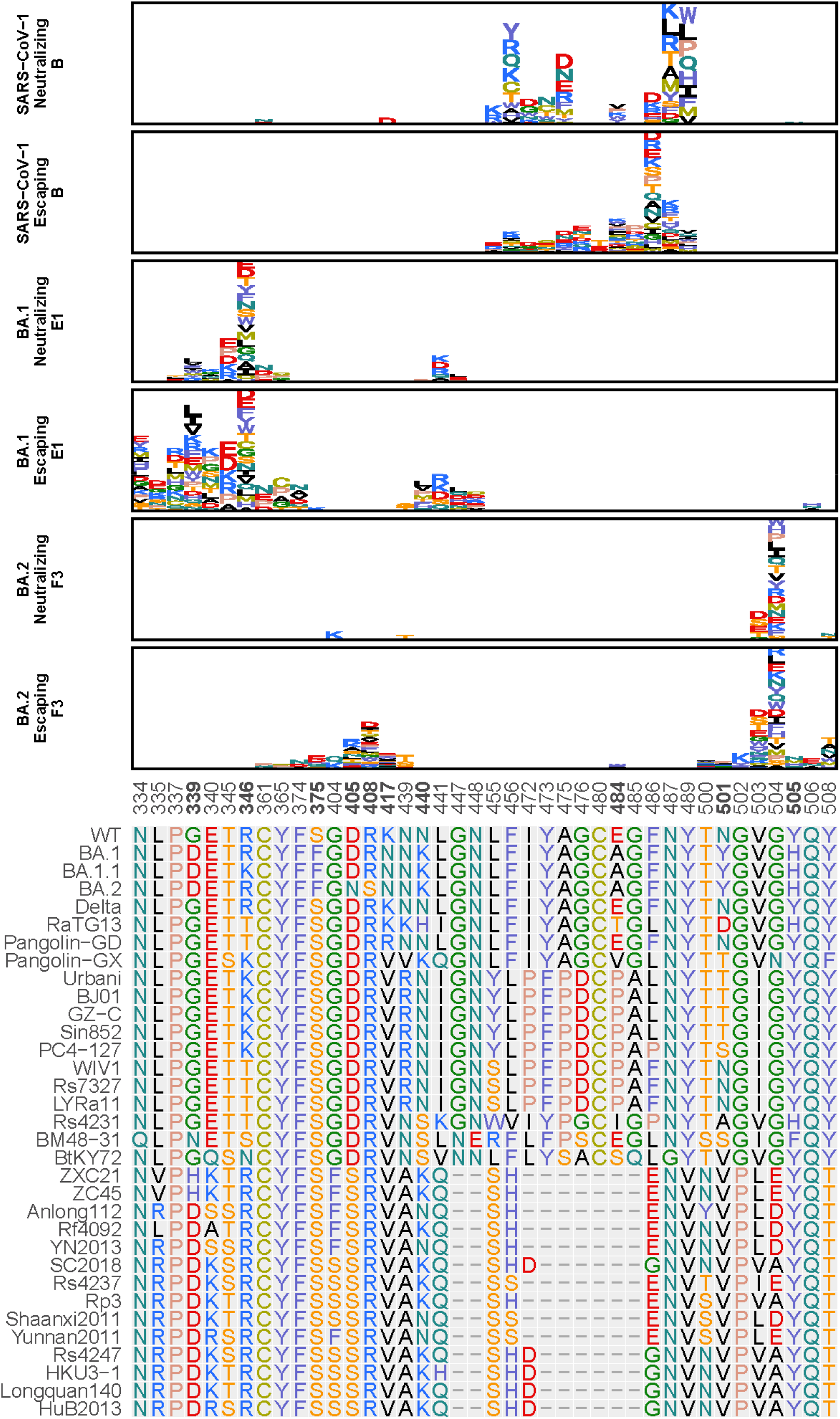
Average escape maps indicate non-homogeneity in function within single epitope groups. Average escape maps of SARS-CoV-1-neutralizing and SARS-CoV-2 Non-neutralizing Group B antibodies, BA.1-neutralizing and escaping Group E1 antibodies, and BA.2-neutralizing and escaping Group F3 antibodies are shown. Height of each amino acid in the escape maps represents its mutation escape score. Residues are colored corresponding to their chemical properties. Mutated sites in Omicron variants (including BA.1, BA.1.1 and BA.2) are marked in bold.

**Extended Data Fig. 8.**
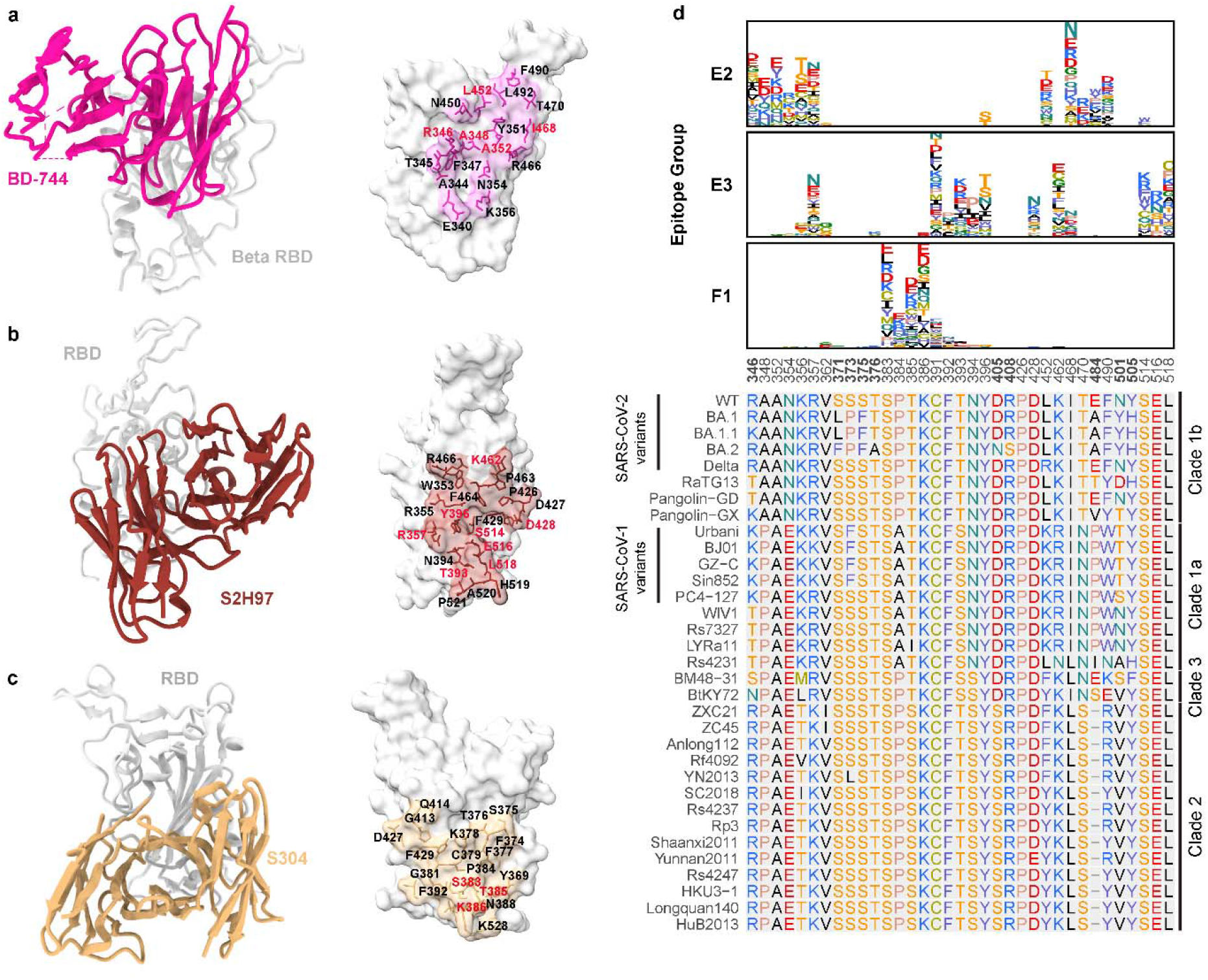
Structural and escaping mutation analyses of group E2, E3, F1 antibodies. **a-c,** High-resolution cryo-electron microscopy antibody structures of representative epitope group E2, E3, F1 neutralizing antibodies. **a,** BD-744 (group E2) in complex of SARS-CoV-2 Beta RBD (PDB: 7EY0). **b,** S2H97 (group E3) in complex of SARS-CoV-2 RBD (PDB: 7M7W). **c,** S304 (group F1) in complex of SARS-CoV-2 RBD (PDB: 7JW0). Residues on the binding interface are marked. Residues highlighted in red indicate featuring escaping hotspots of the representative epitope groups. **d,** Averaged escape maps of antibodies in epitope group E2, E3, F1, and corresponding multiple sequence alignment (MSA) of various sarbecovirus RBDs. Height of each amino acid in the escape maps represents its mutation escape score. Residues are colored corresponding to their chemical properties. Mutated sites in Omicron variants are marked in bold.

**Extended Data Fig. 9.**
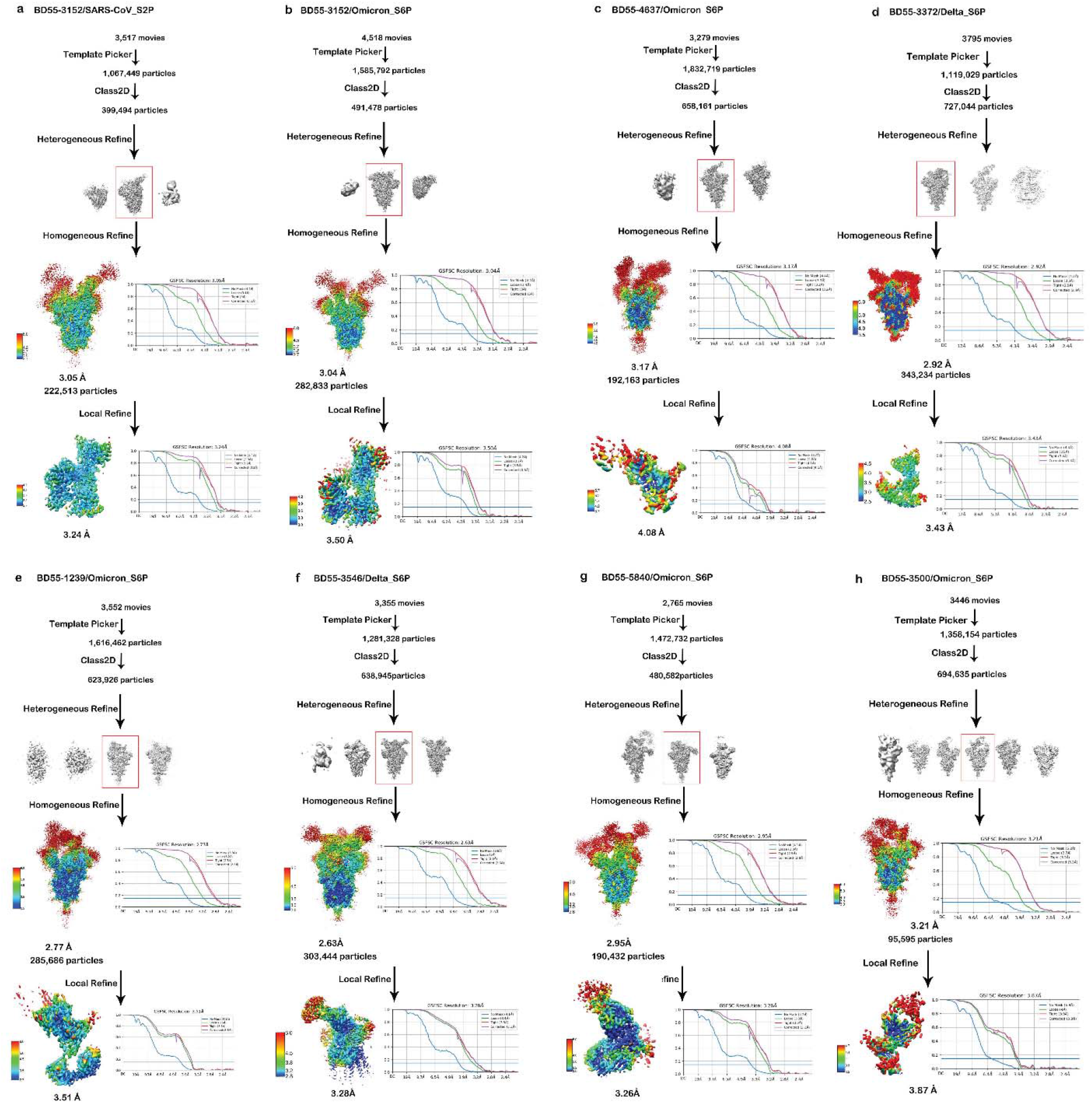
Workflow for the 3D reconstruction of the cryo-EM structures for antibody epitope studies. Workflow for the 3D reconstruction of the cryo-EM structures of the SARS-CoV-1 S2P, Omicron BA.1 S6P, and Delta S6P trimers in complex with RBD neutralizing antibodies **a-b,** BD55-3152 **c,** BD55-4637 **d,** BD55-3372 **e,** BD55-1239 **f,** BD55-3546 **g,** BD55-5840 **h,** and BD55-3500.

**Extended Data Fig. 10.**
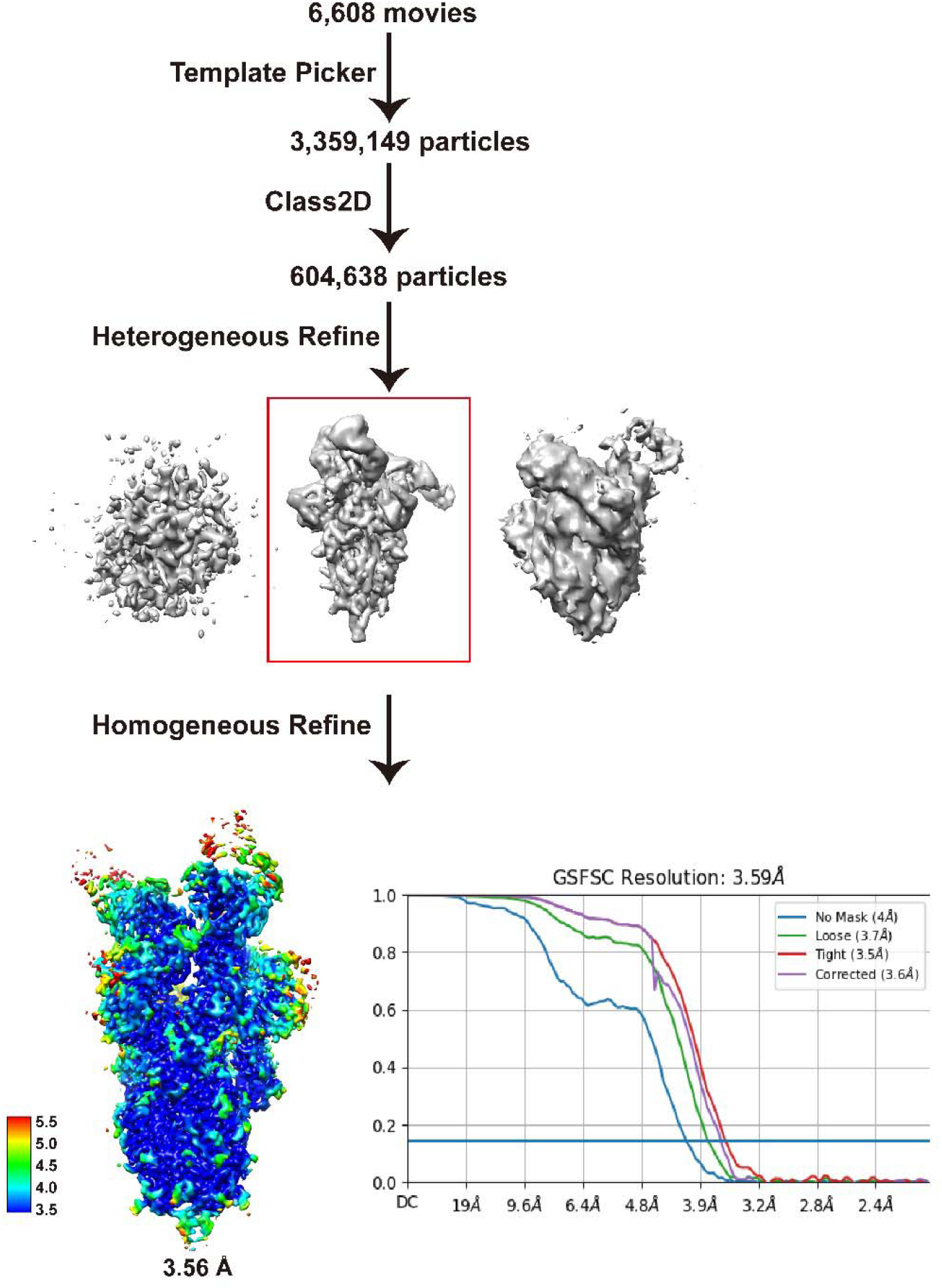
Workflow for the 3D reconstruction of the cryo-EM structures of BA.2 spike in complex with BD55-5840. Workflow for the 3D reconstruction of the cryo-EM structures of the Omicron BA.1 S6P trimers in complex with BD55-5840.

