## Supplementary figures and images for "Omicron BA.2 specifically evades broad sarbecovirus neutralizing antibodies"

### Supplementary Data 2

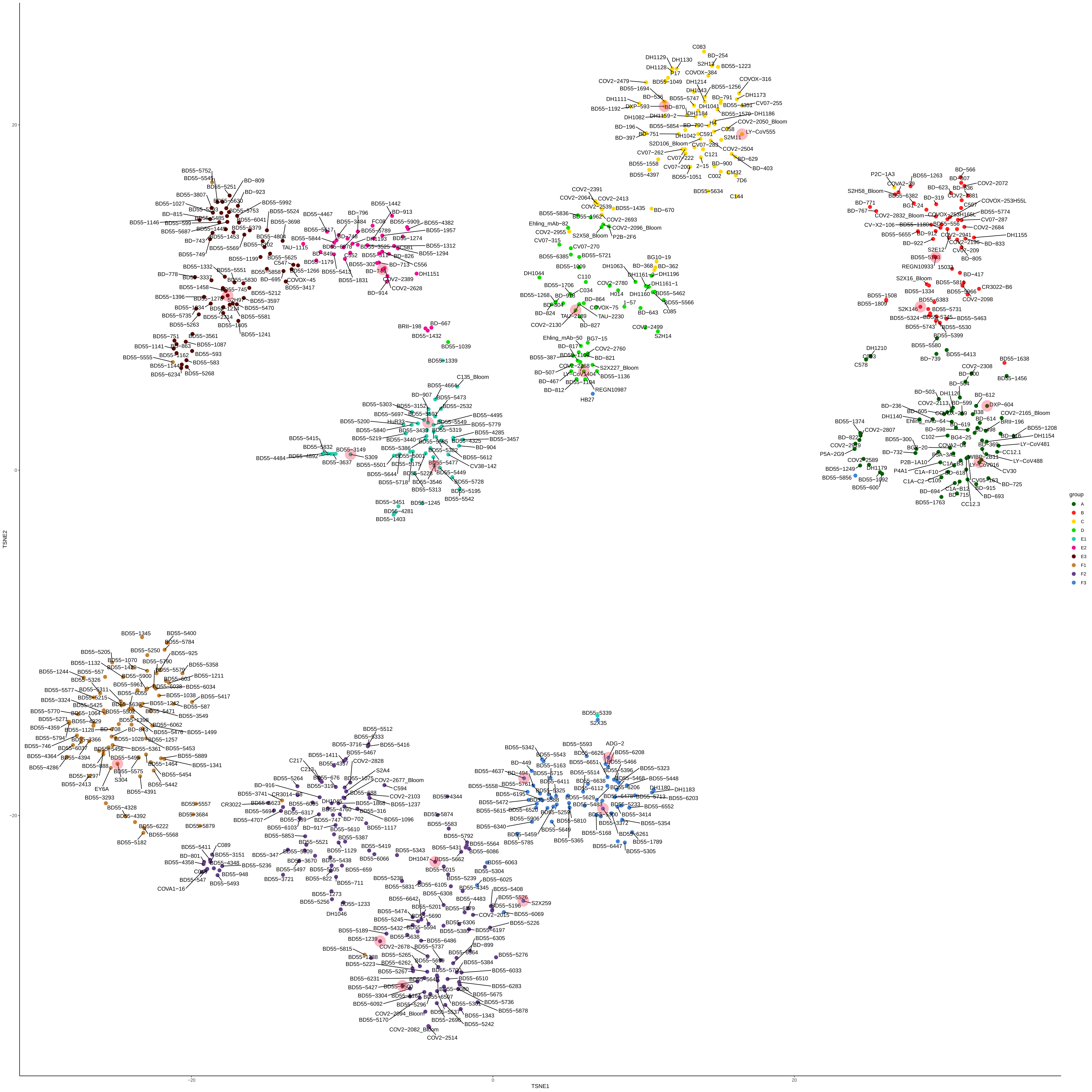

### Supplementary Data 3

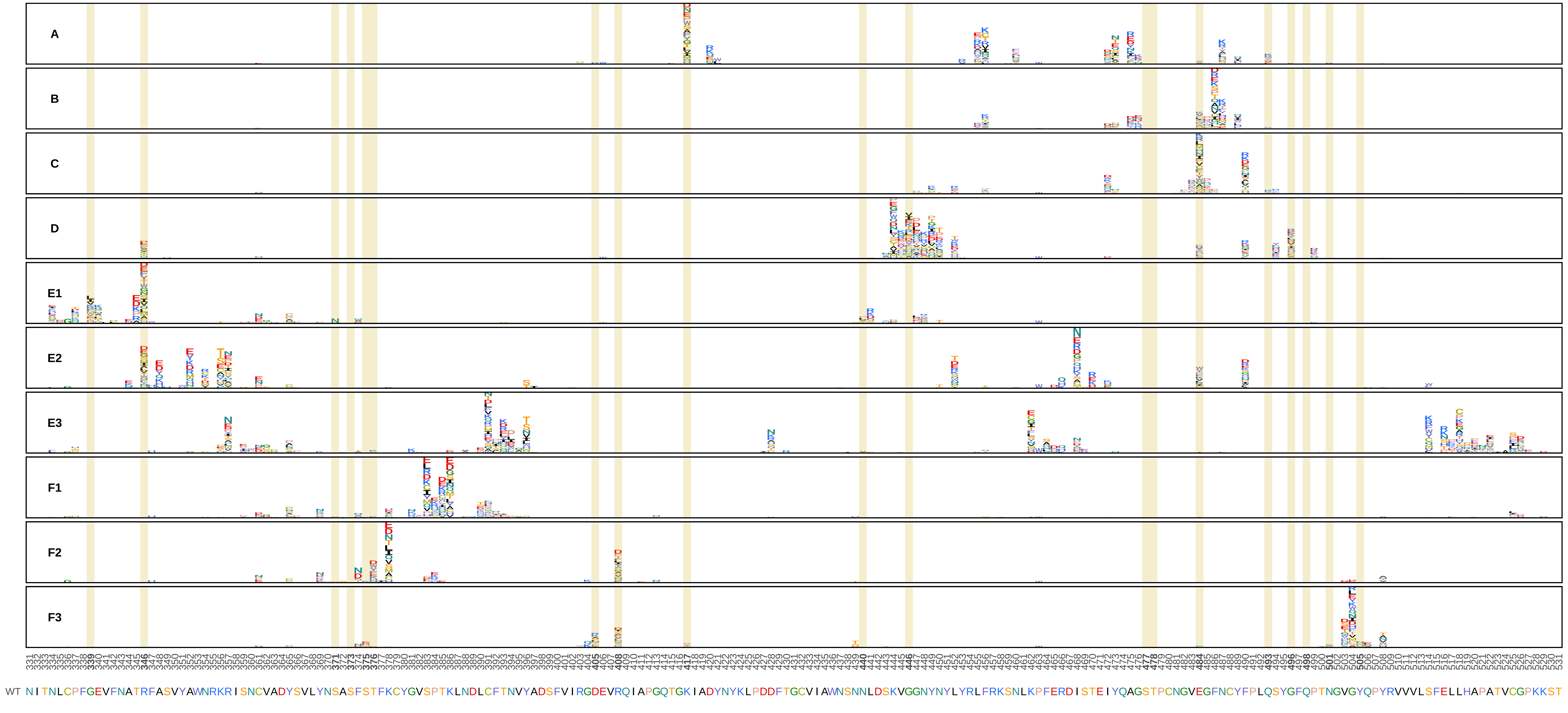
