## Supplementary Guide for "Omicron BA.2 specifically evades broad sarbecovirus neutralizing antibodies"

**Supplementary Table 1**

Summarized information of plasma donating volunteers, including their age, sex, types, doses, the interval of the vaccines received, date of BA.1 infection onset and hospitalization discharge, and the date of blood sampling.

**Supplementary Table 2**

Summarized information and experiment results of 714 RBD antibodies involved in this study, including their sources, epitope groups, pseudovirus neutralizing IC50 and ELISA OD450 for sarbecovirus, ACE2 competition levels, and heavy/light chain sequences.

**Supplementary Table 3**

Accession numbers of sequences of sarbecovirus used in the study.

**Supplementary Table 4**

Cryo-EM data collection and refinement statistics of the eight neutralizing antibodies in complex with SARS-CoV-2 or SARS-CoV-1 spike glycoprotein.

**Supplementary Data 1**

Logo plots of escape maps of SARS-CoV-2 RBD antibodies of 10 epitope groups. Sites mutated frequently in Omicron variants are highlighted in gold. R346 is highlighted in red, and mutated sites in BA.2 are highlighted in blue.

**Supplementary Data 2**

Unsupervised clustering of the 714 antibodies with each antibody’s ID shown.

**Supplementary Data 3**

Averaged escape maps of antibodies in all epitope groups displayed in the linear order. Residues are colored corresponding to the chemical properties. Sites mutated frequently in Omicron variants are highlighted in gold. R346 is highlighted in red, and mutated sites in BA.2 are highlighted in blue.
