## Supplementary Data 1 for "Omicron BA.2 specifically evades broad sarbecovirus neutralizing antibodies"

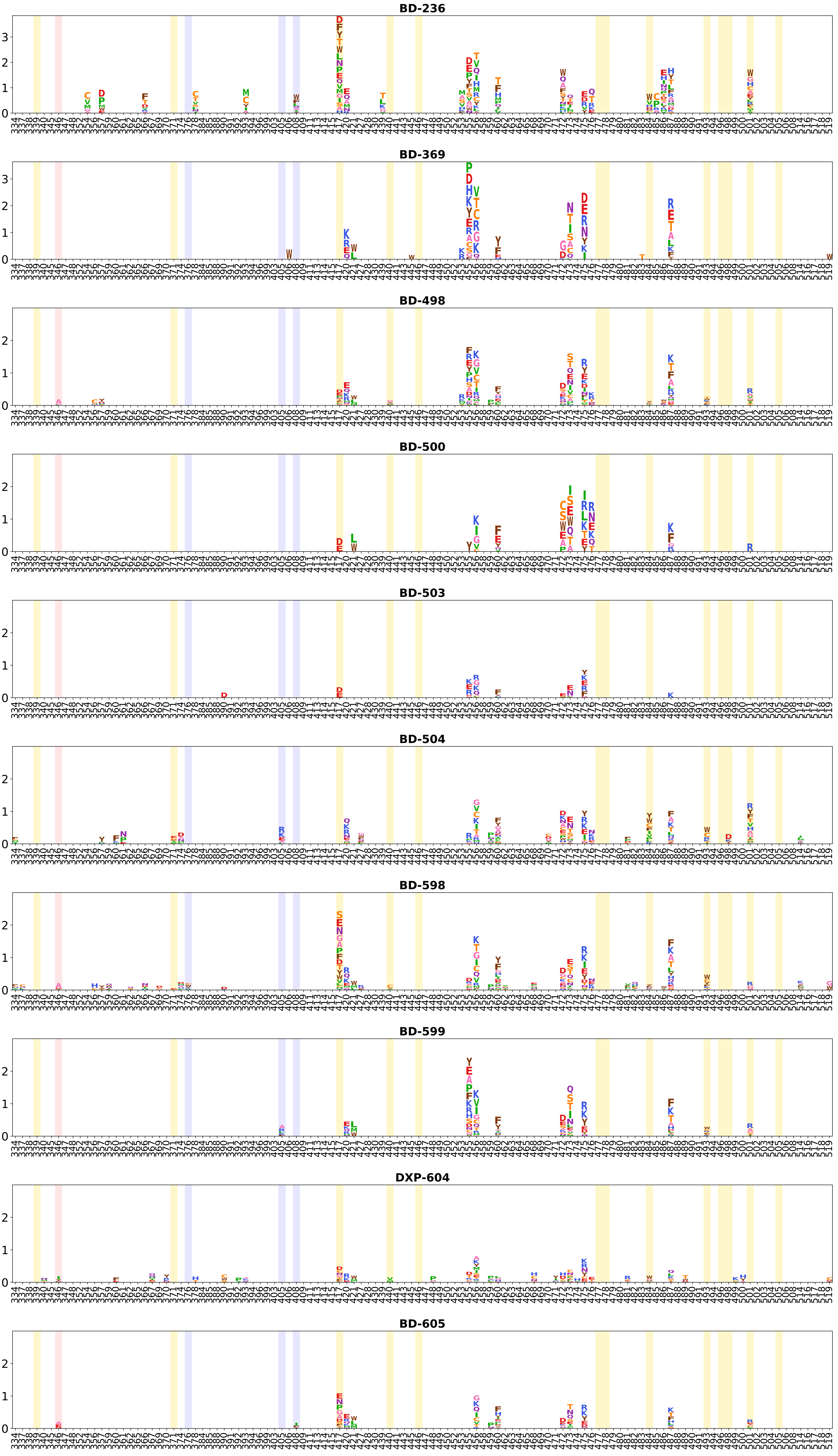
BD-236
BD-369
BD-498
BD-500
BD-503
BD-504
BD-598
BD-599
DXP-604
BD-605

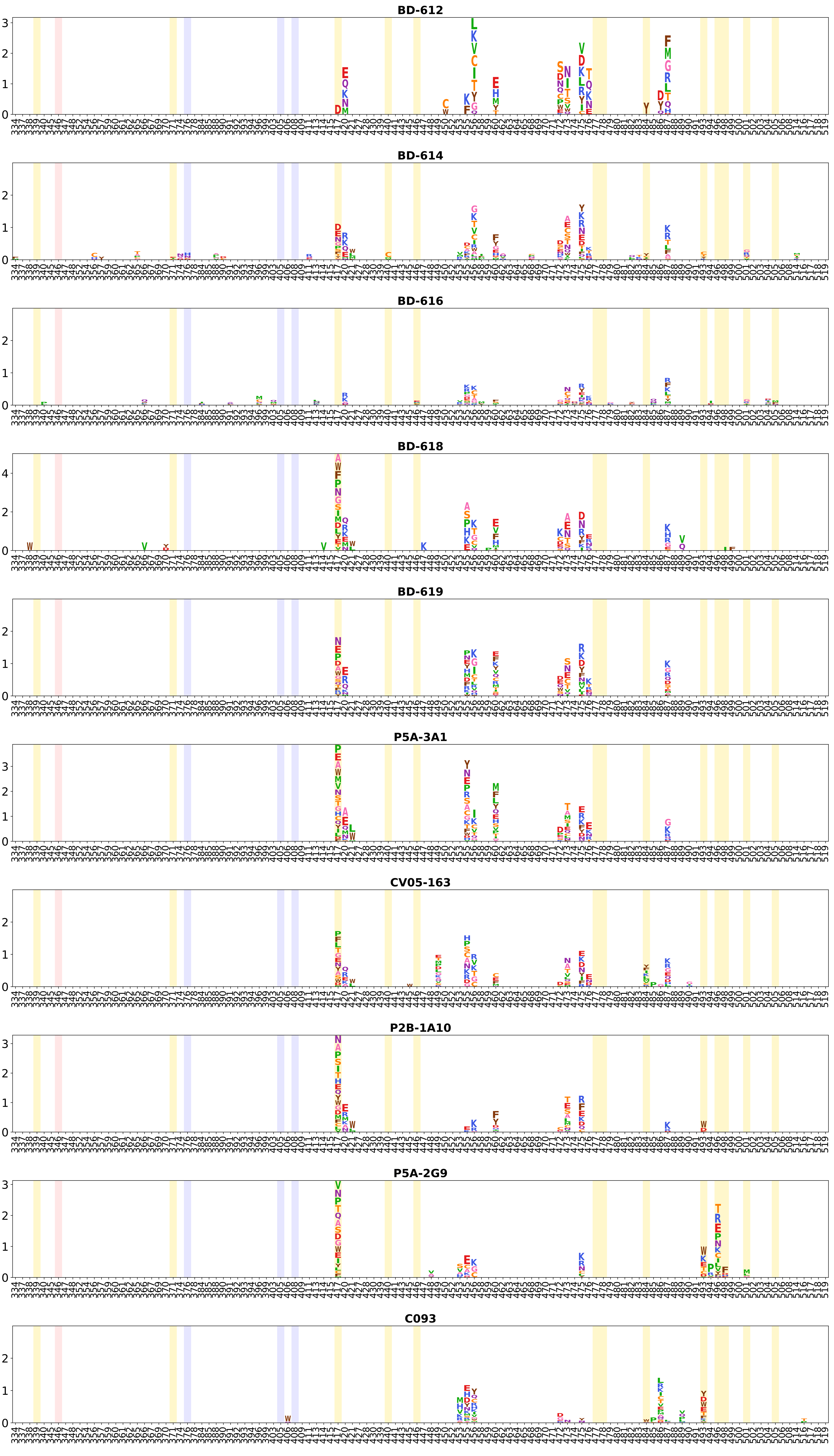
BD-612
BD-614
BD-616
BD-618
BD-619
P5A-3A1
CV05-163
P2B-1A10
P5A-2G9
C093

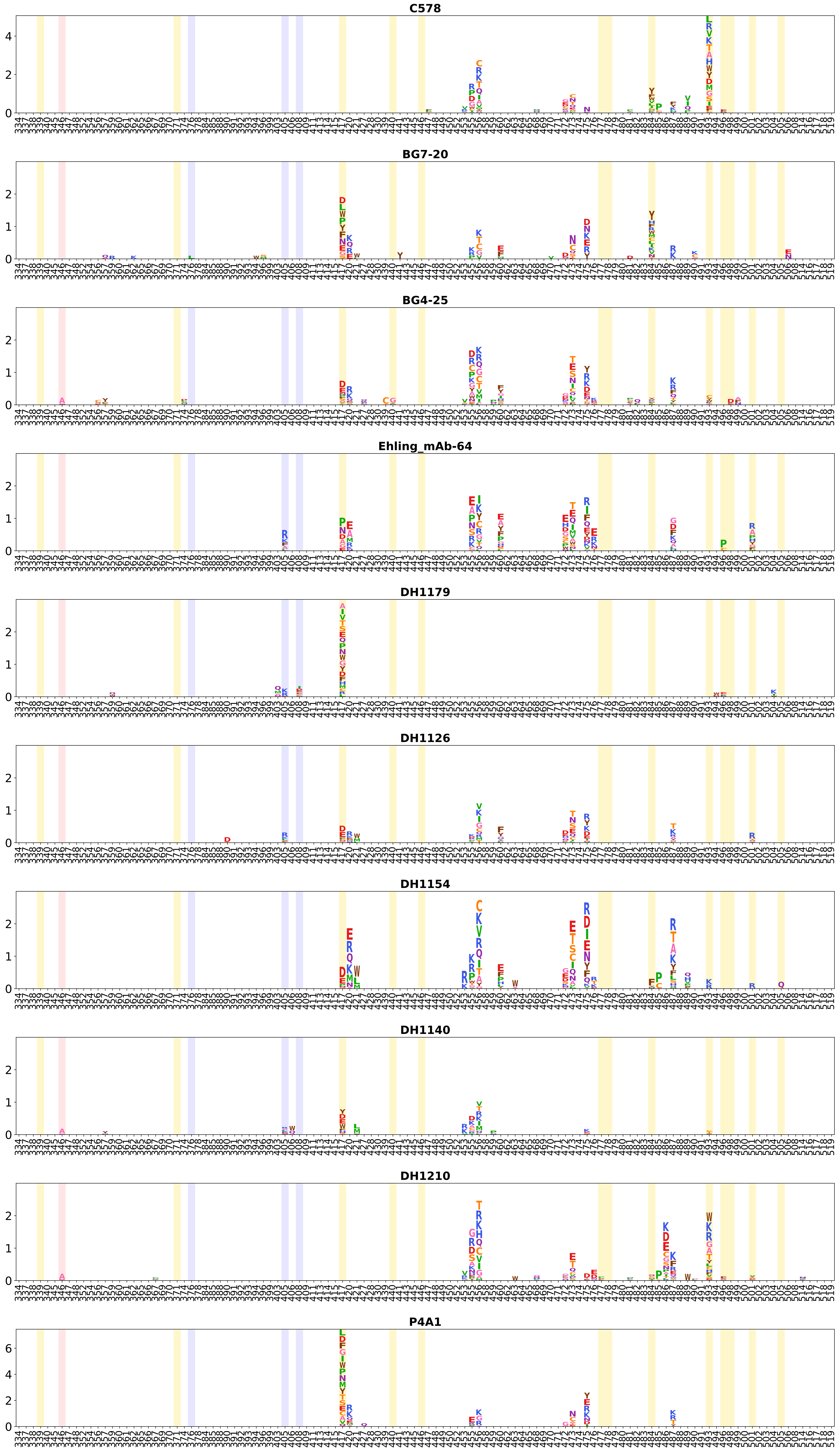
C578
BG7-20
BG4-25
Ehling_mAb-64
DH1179
DH1126
DH1154
DH1140
DH1210
P4A1

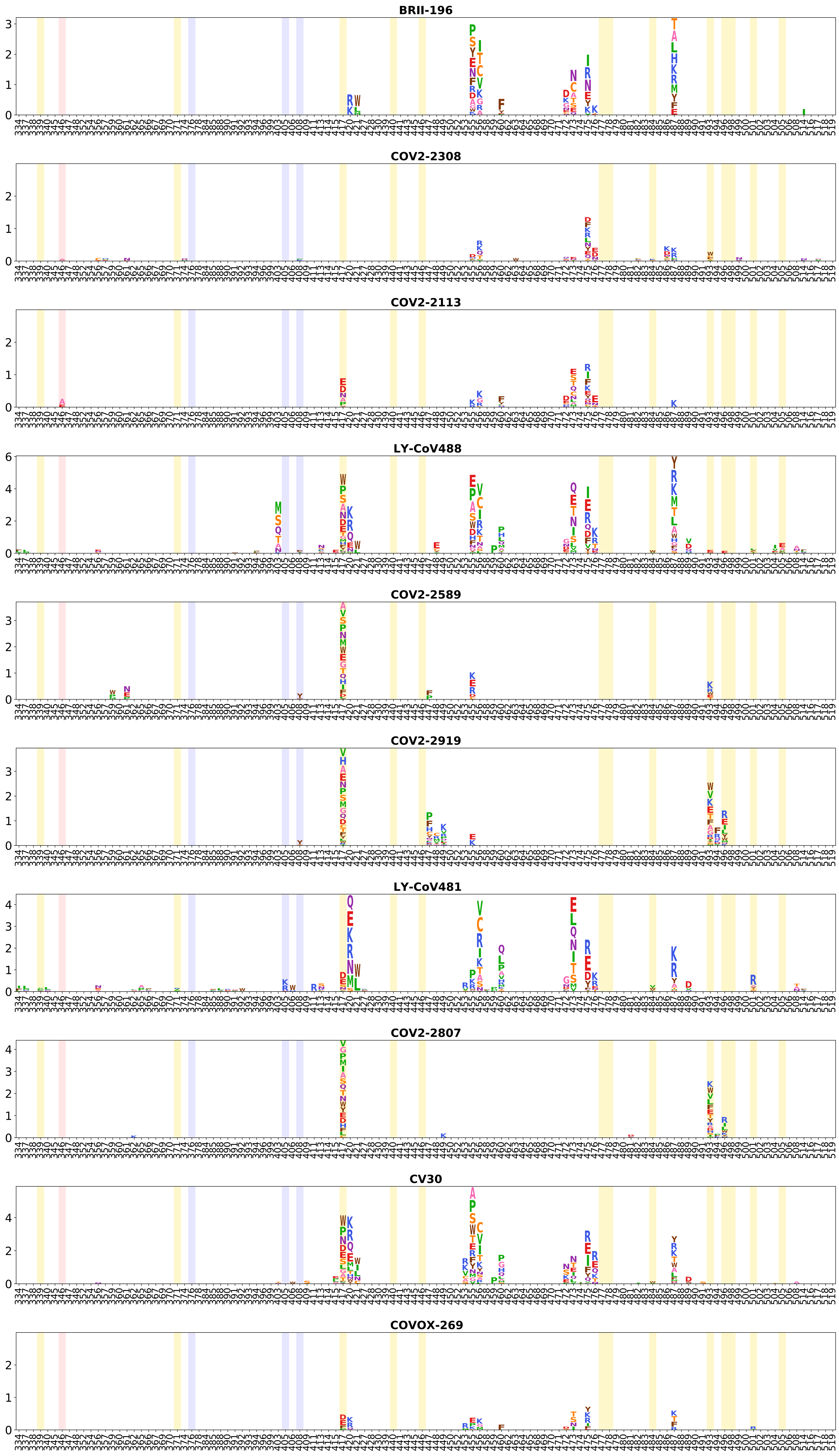
BRII-196
COV2-2308
COV2-2113
LY-CoV488
COV2-2589
COV2-2919
LY-CoV481
COV2-2807
CV30
COVOX-269

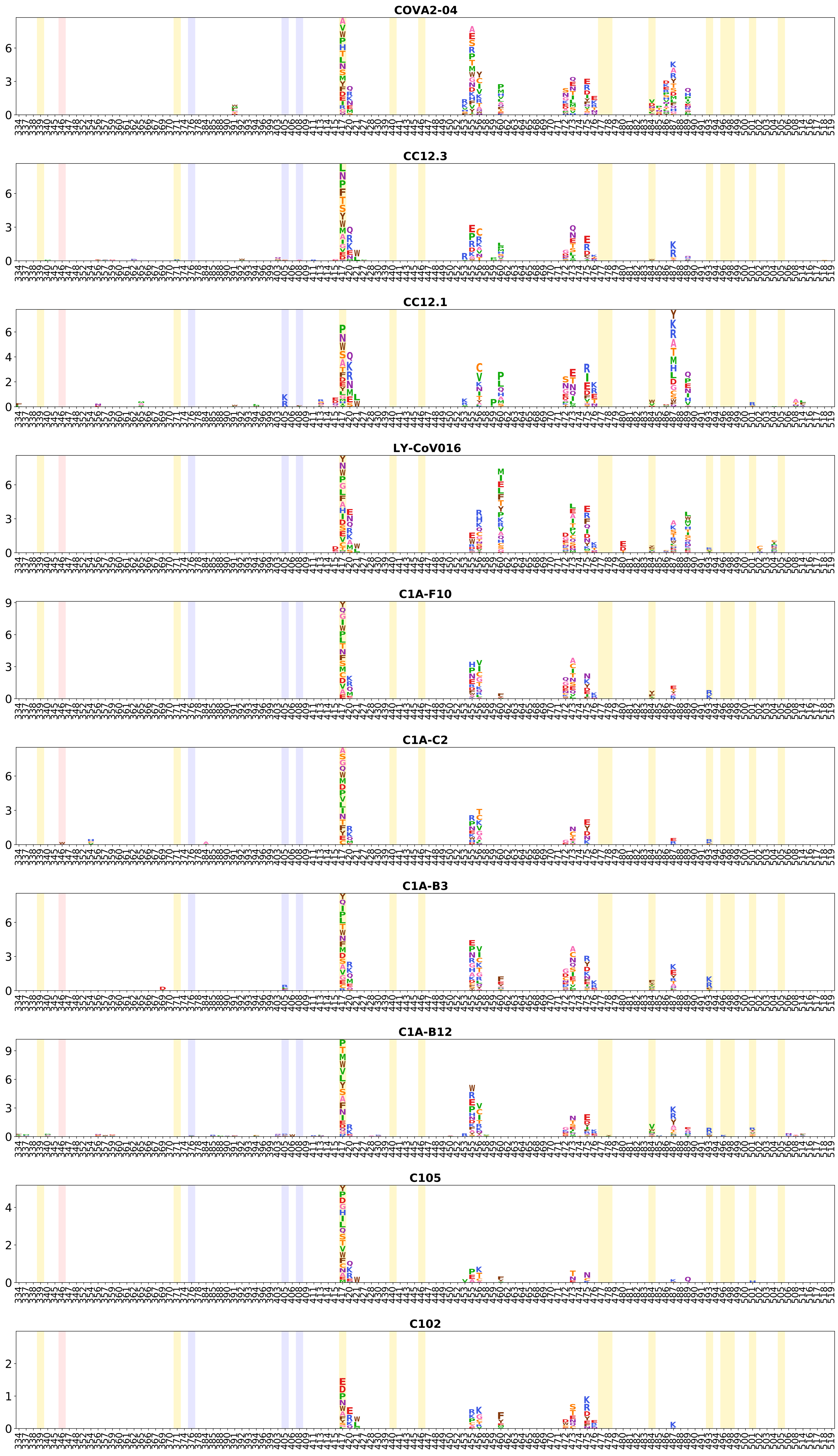
COVA2-04
CC12.3
CC12.1
LY-CoV016
C1A-F10
C1A-C2
C1A-B3
C1A-B12
C105
C102

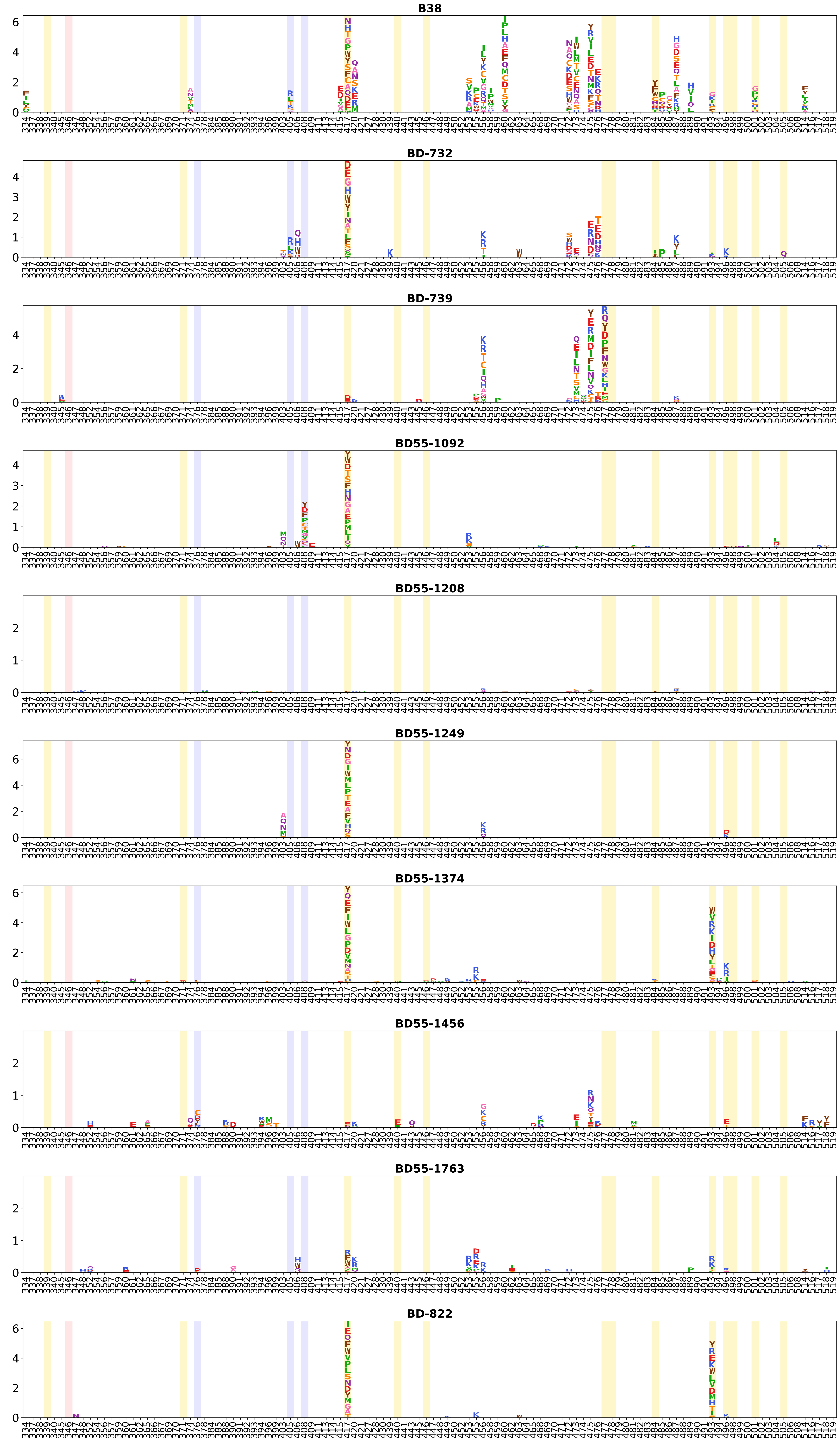
B38
BD-732
BD-739
BD55-1092
BD55-1208
BD55-1249
BD55-1374
BD55-1456
BD55-1763
BD-822

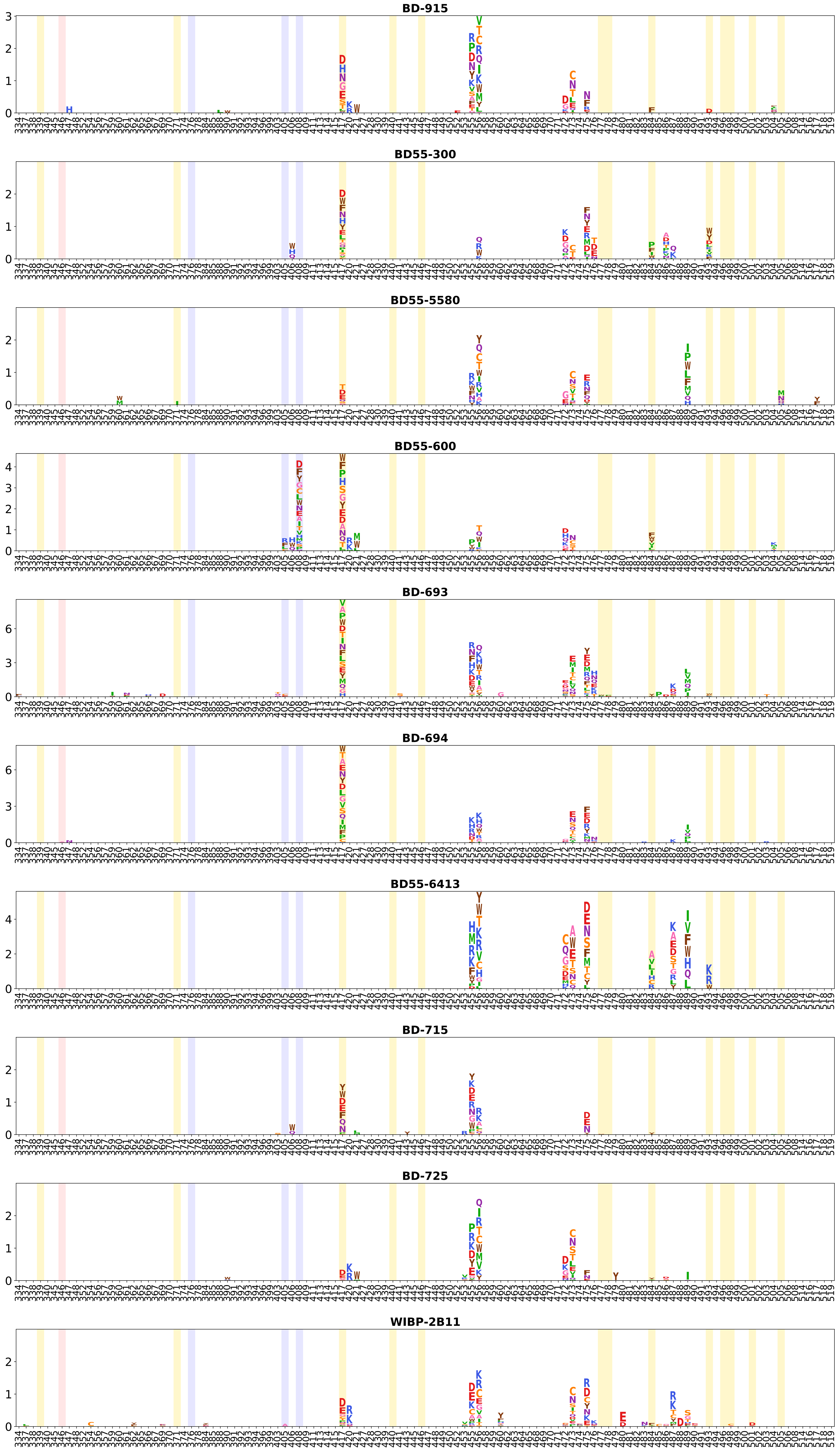
BD-915
BD55-300
BD55-5580
BD55-600
BD-693
BD-694
BD55-6413
BD-715
BD-725
WIBP-2B11

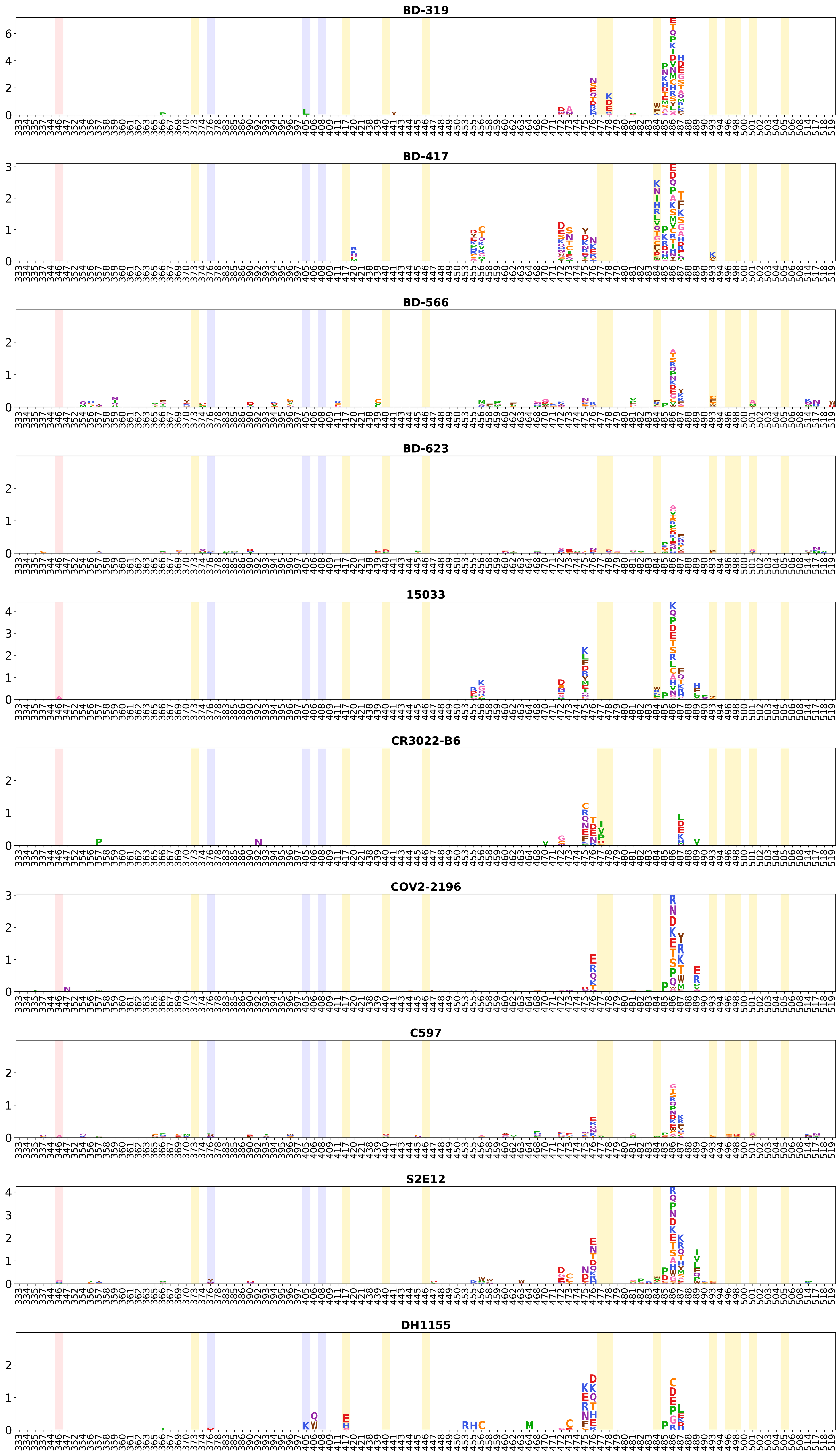

BD-319
BD-417
BD-566
BD-623
15033
CR3022-B6
COV2-2196
C597
S2E12
DH1155

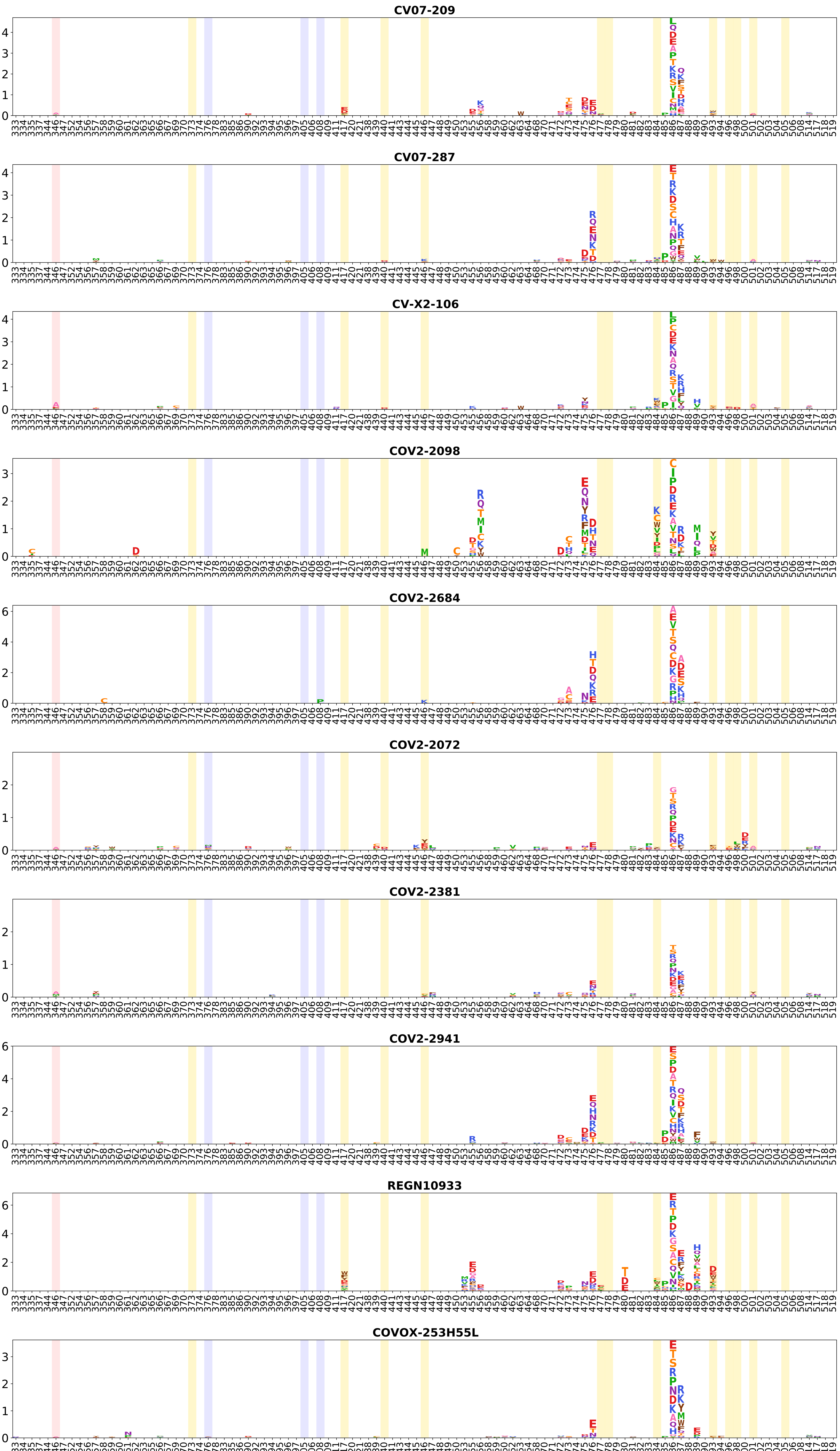
CV07-209
CV07-287
CV-X2-106
COV2-2098
COV2-2684
COV2-2072
COV2-2381
COV2-2941
REGN10933
COVOX-253H55L

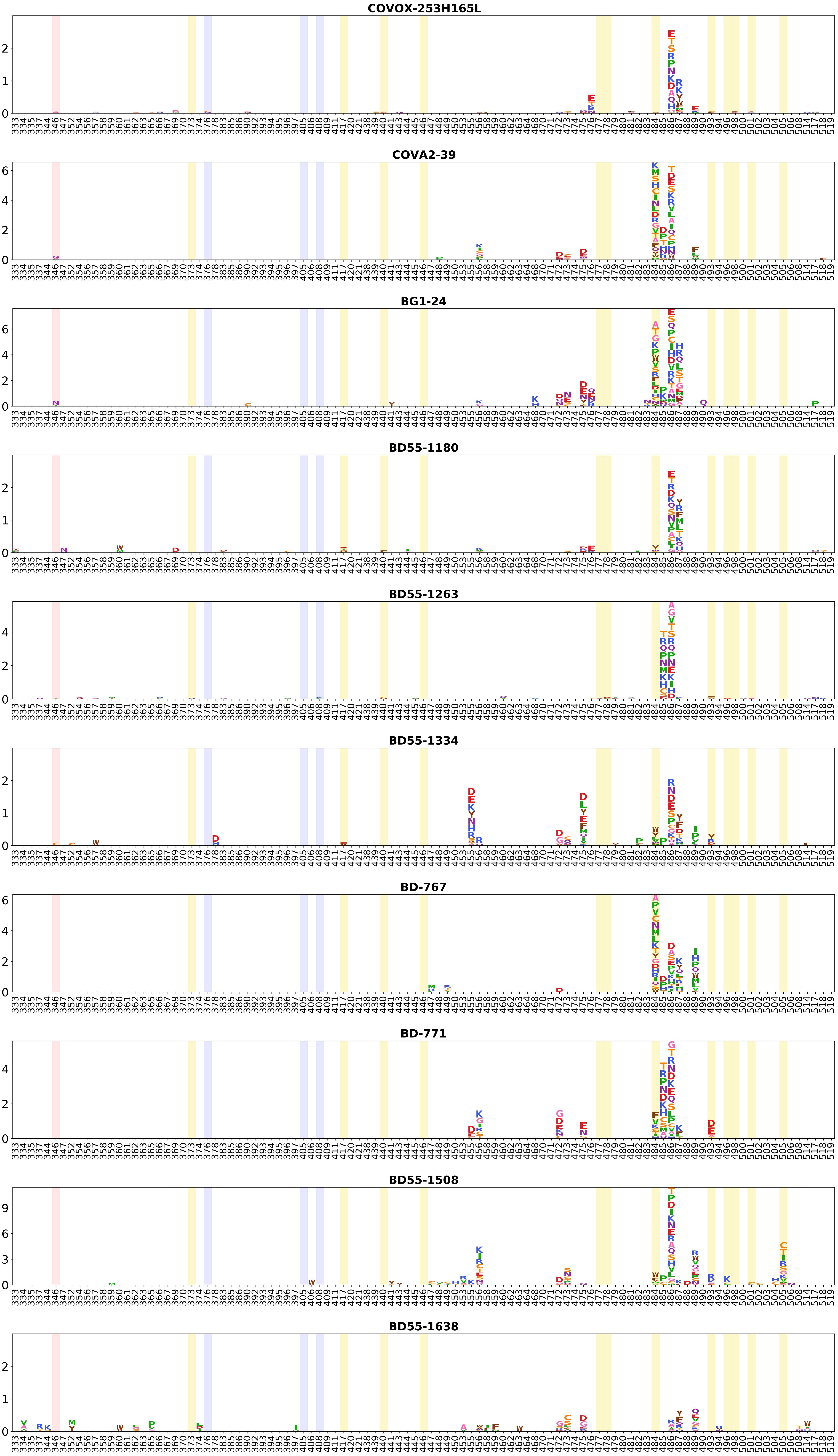
COVOX-253H165L
COVA2-39
BG1-24
BD55-1180
BD55-1263
BD55-1334
BD-767
BD-771
BD55-1508
BD55-1638

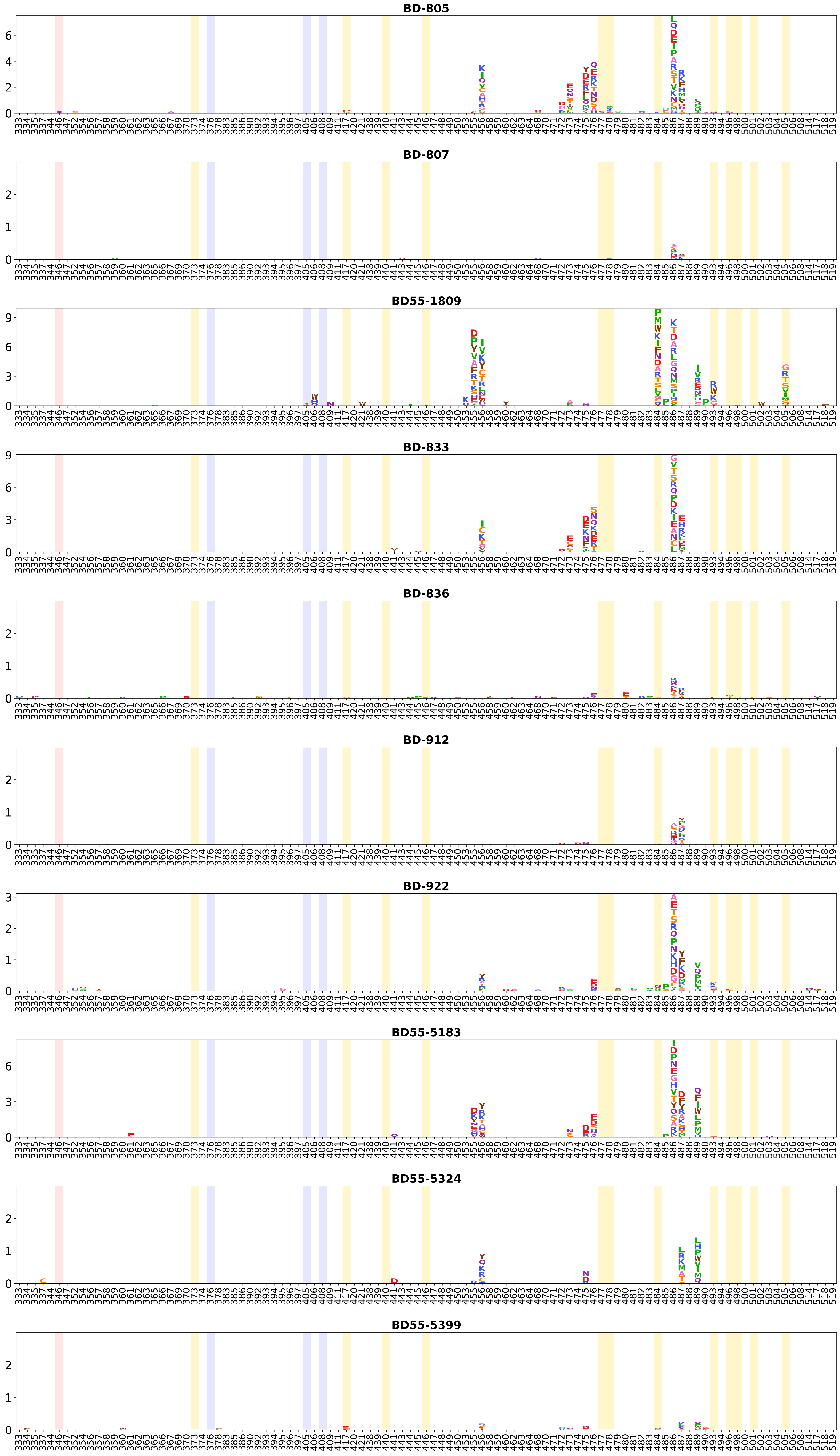
BD-805
BD-807
BD55-1809
BD-833
BD-836
BD-912
BD-922
BD55-5183
BD55-5324
BD55-5399

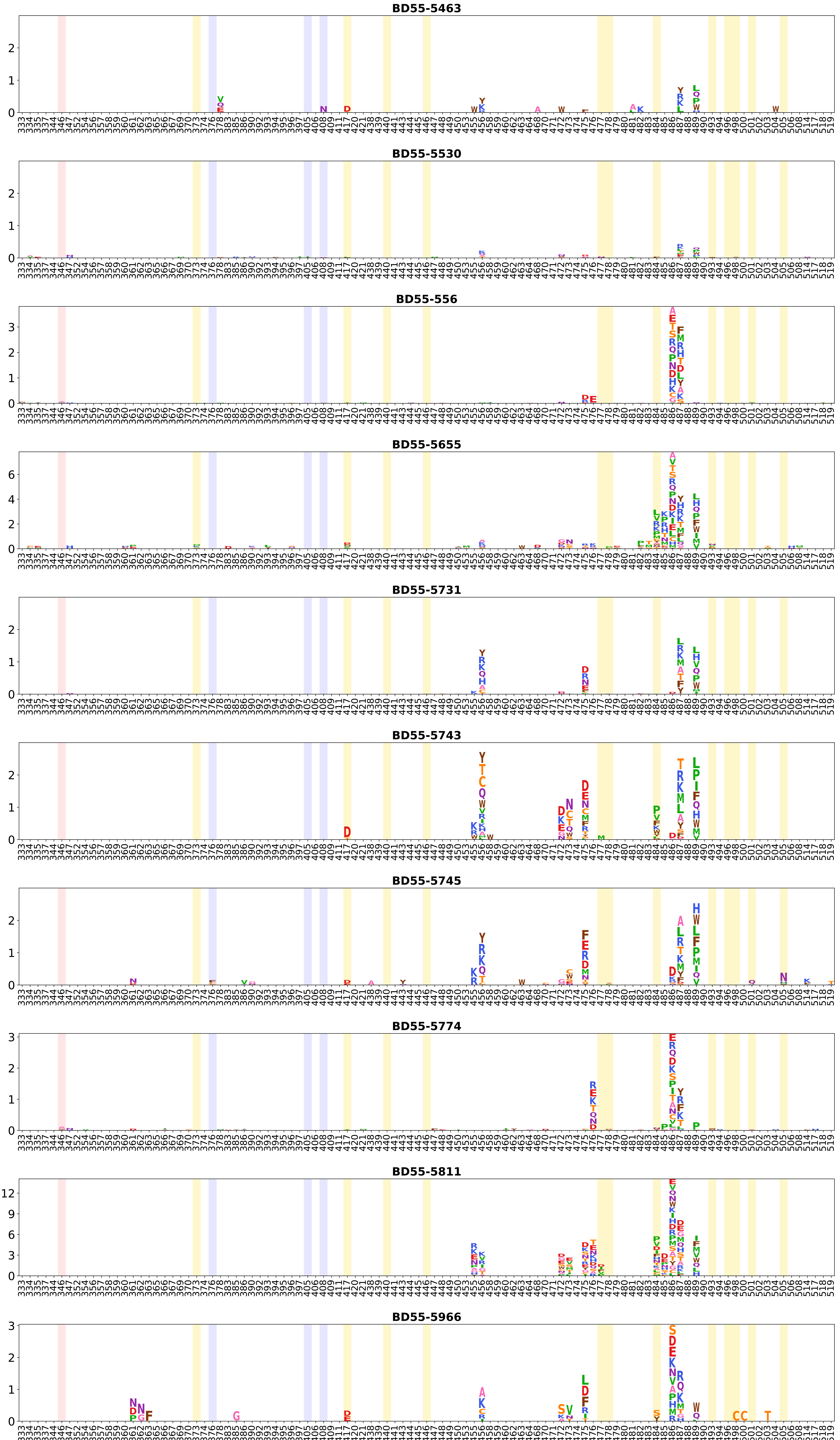
BD55-5463
BD55-5530
BD55-556
BD55-5655
BD55-5731
BD55-5743
BD55-5745
BD55-5774
BD55-5811
BD55-5966

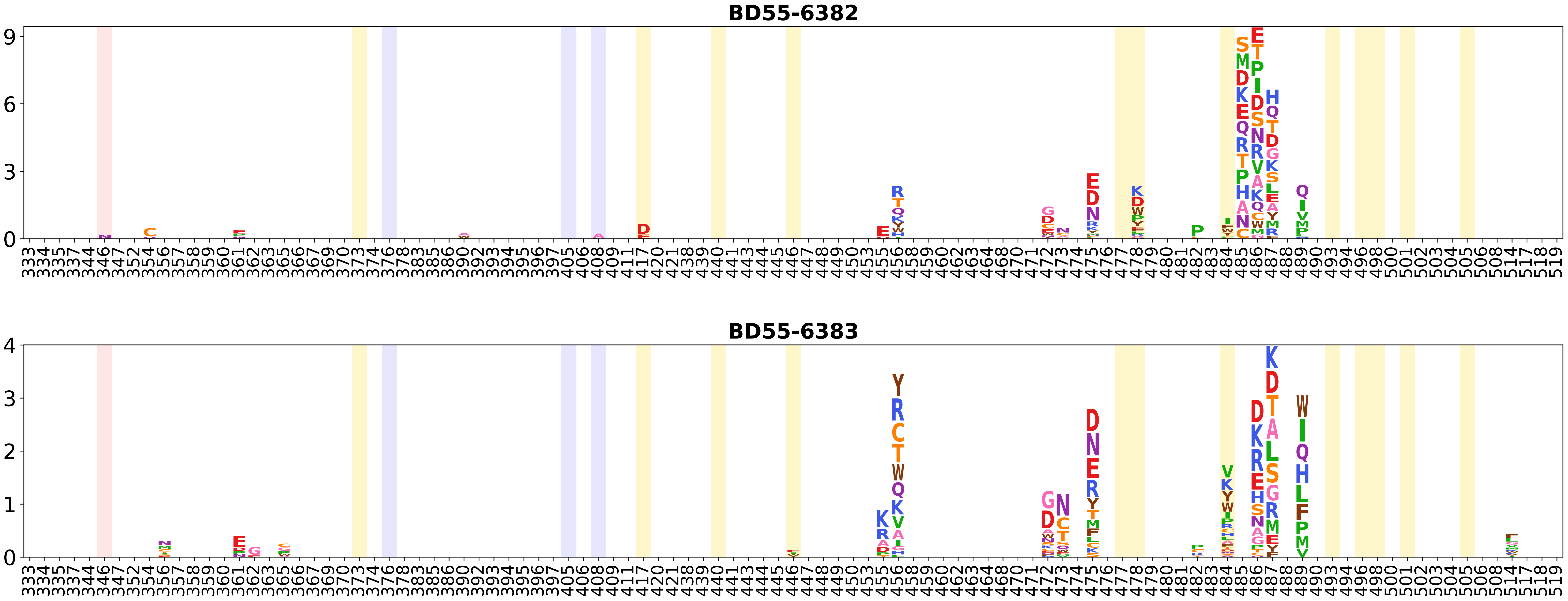
BD55-6382
BD55-6383

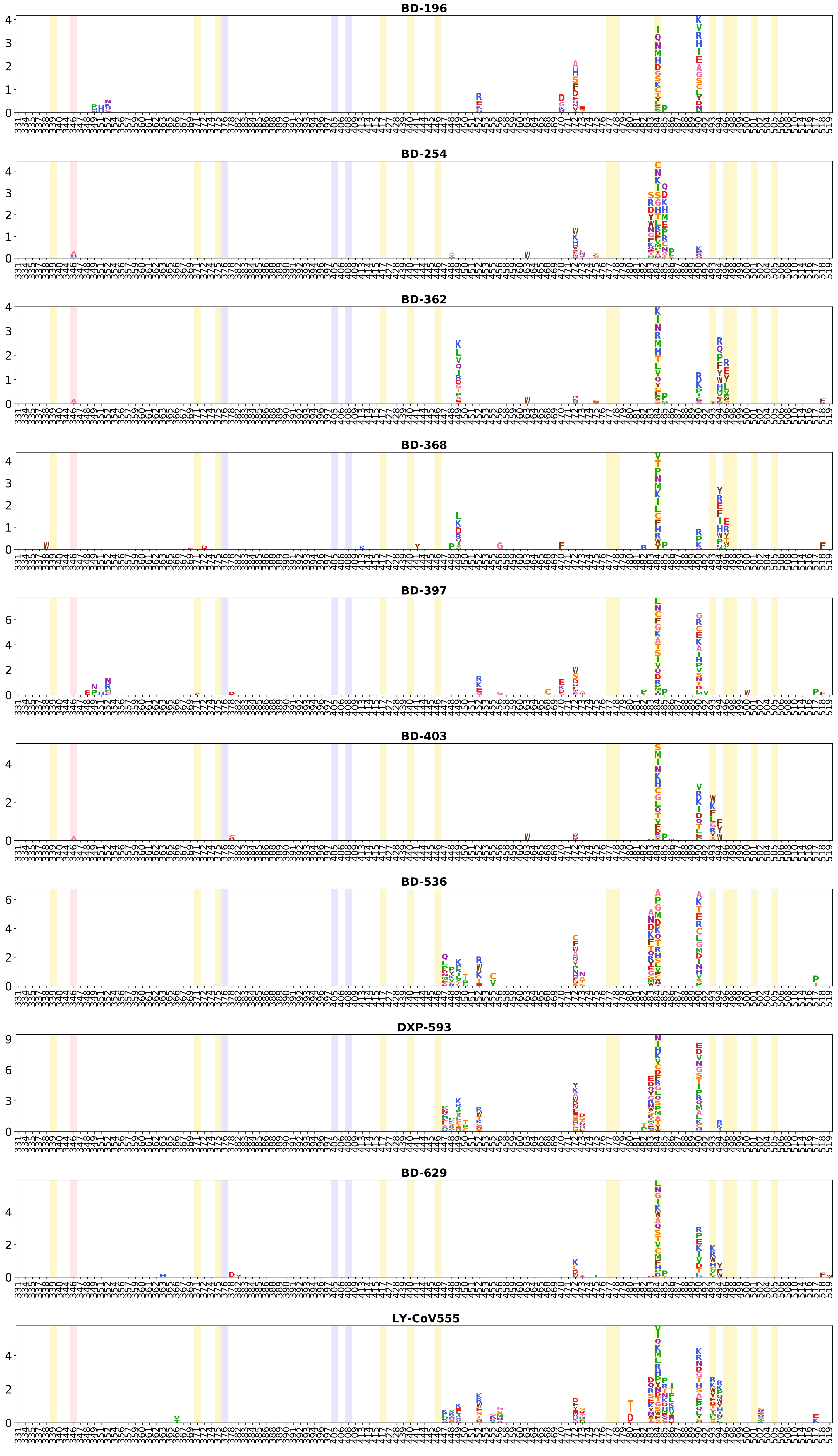
BD-196
BD-254
BD-362
BD-368
BD-397
BD-403
BD-536
DXP-593
BD-629
LY-CoV555

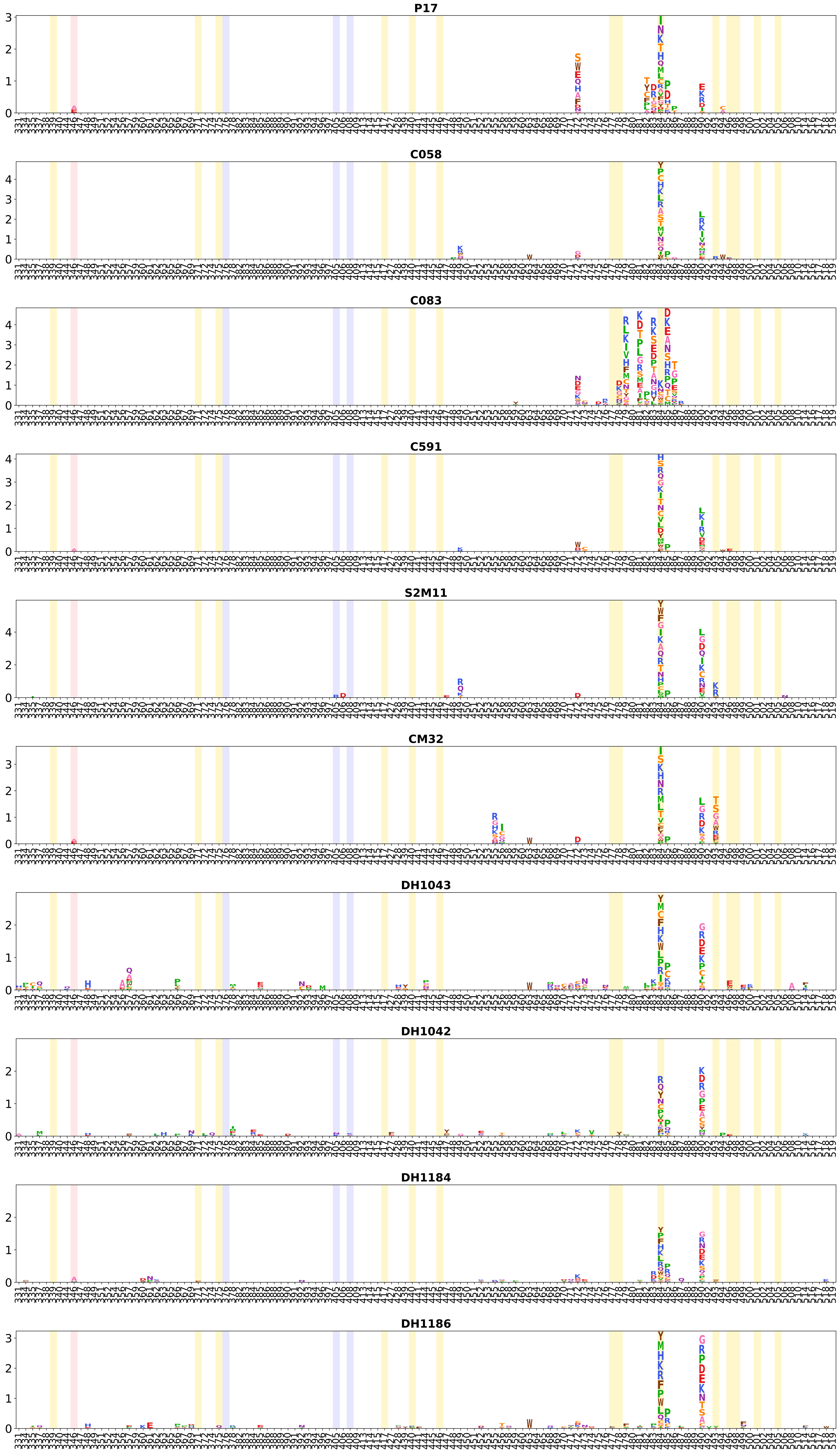

P17
C058
C083
C591
S2M11
CM32
DH1043
DH1042
DH1184
DH1186

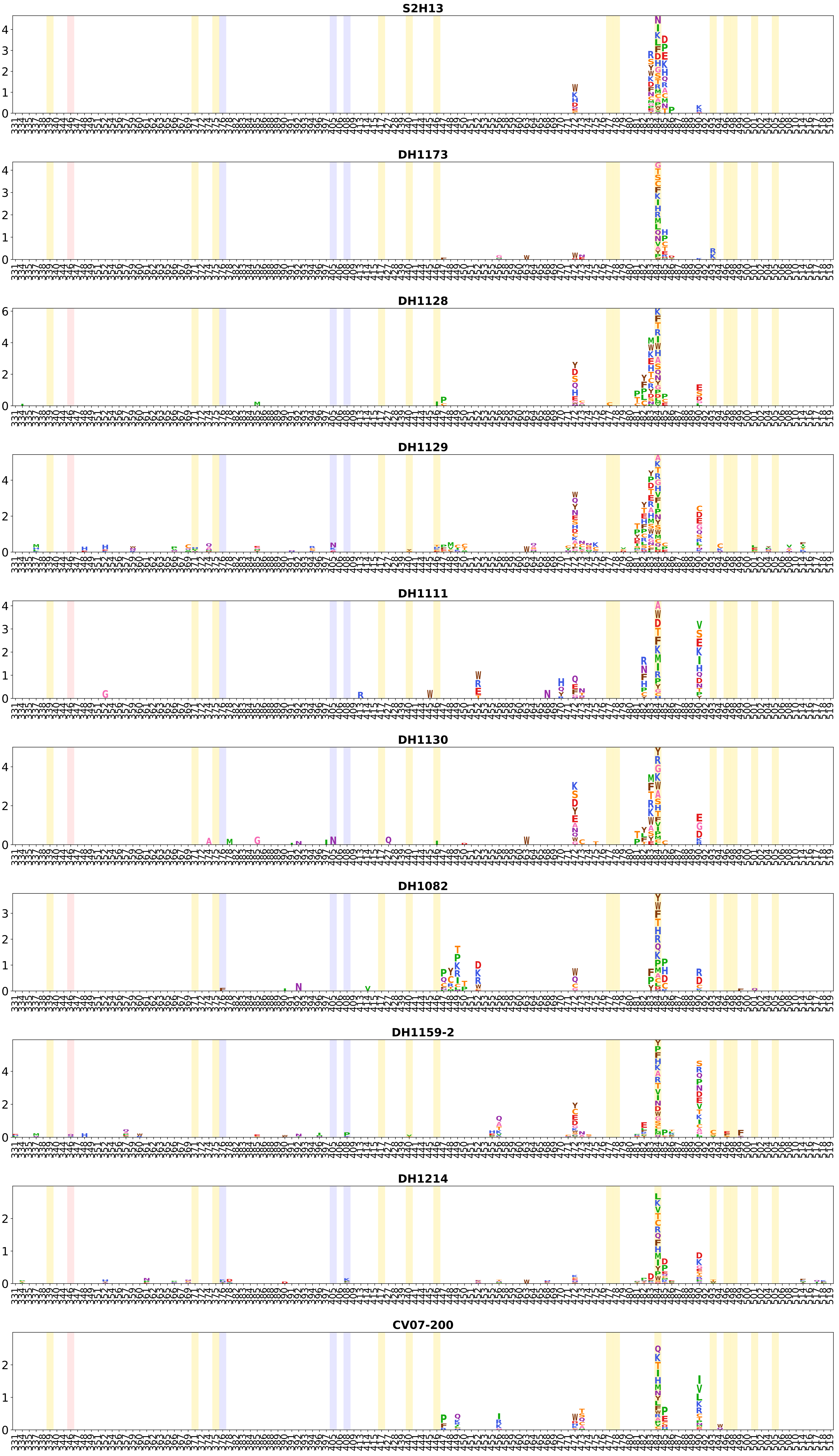
S2H13
DH1173
DH1128
DH1129
DH1111
DH1130
DH1082
DH1159-2
DH1214
CV07-200

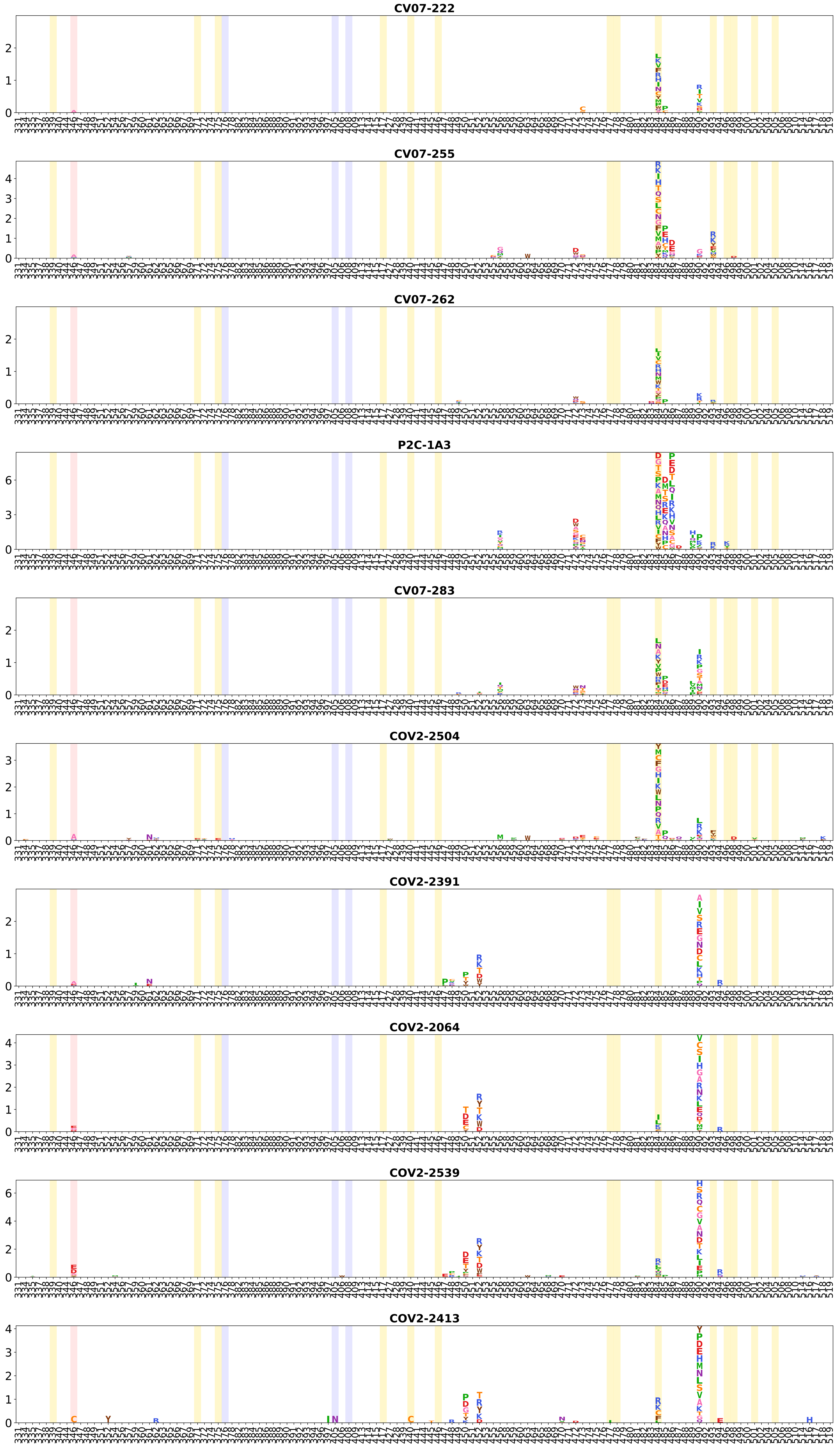
CV07-222
CV07-255
CV07-262
P2C-1A3
CV07-283
COV2-2504
COV2-2391
COV2-2064
COV2-2539
COV2-2413

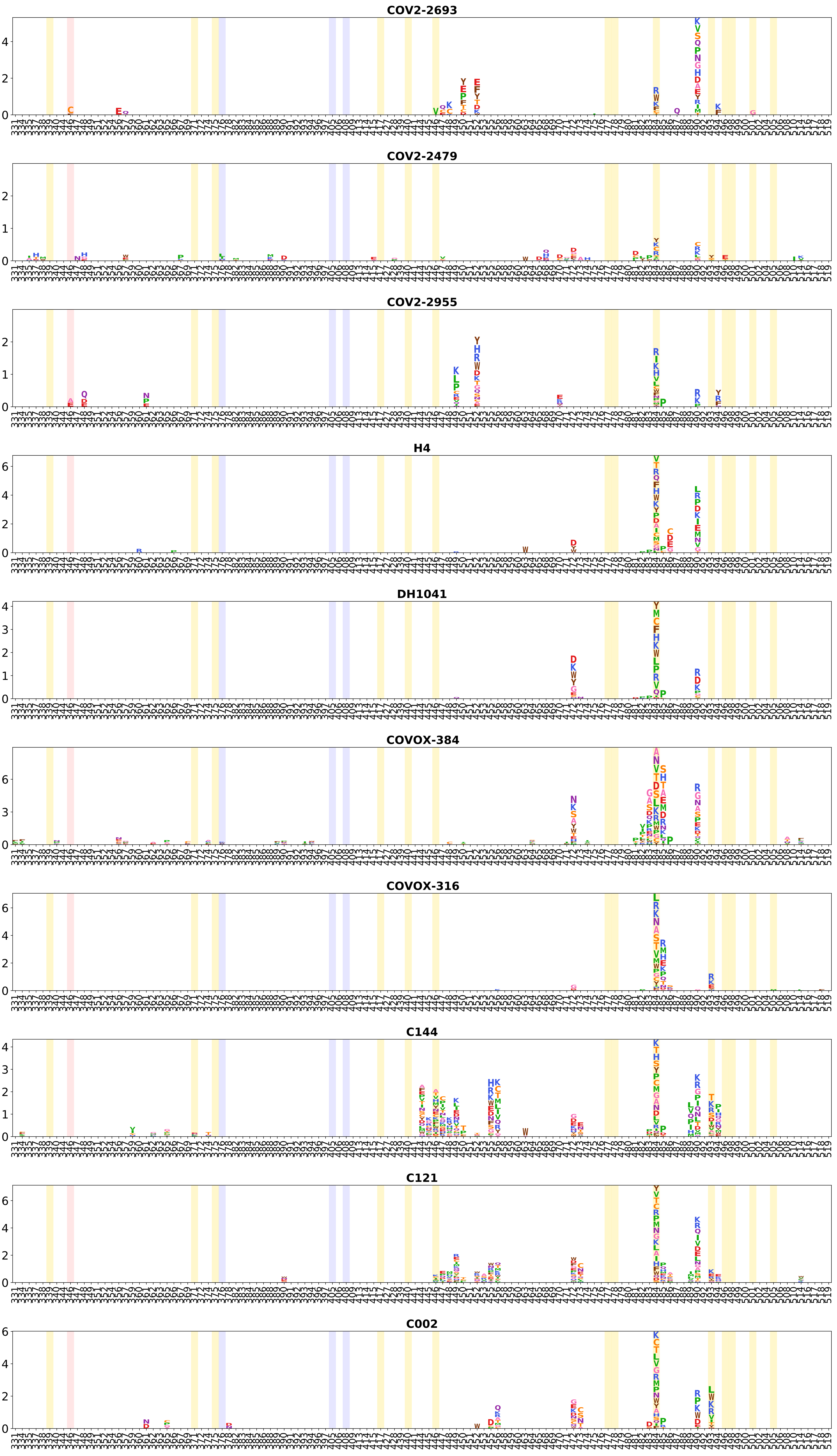
COV2-2693
COV2-2479
COV2-2955
H4
DH1041
COVOX-384
COVOX-316
C144
C121
C002

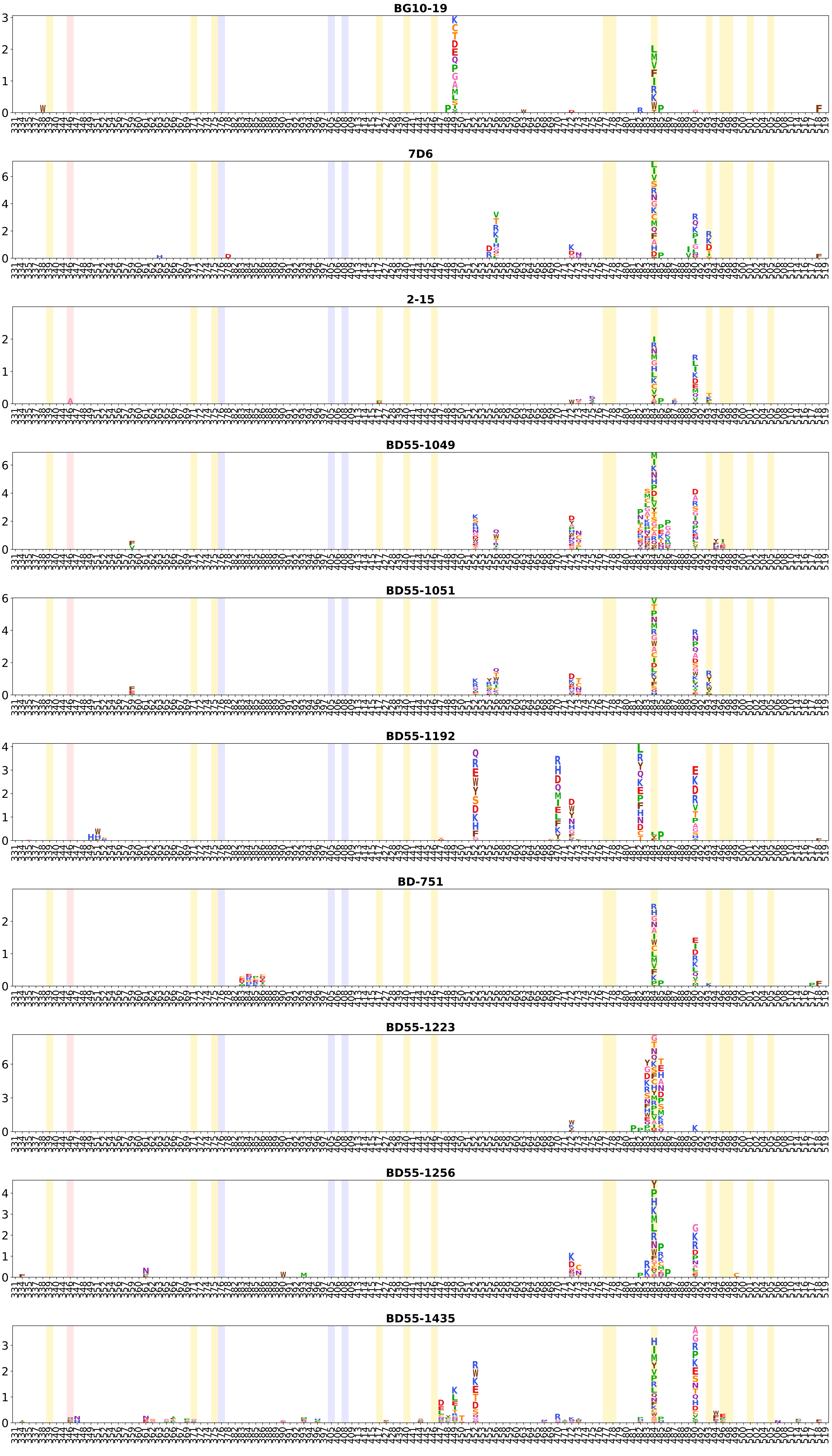
BG10-19
7D6
2-15
BD55-1049
BD55-1051
BD55-1192
BD-751
BD55-1223
BD55-1256
BD55-1435

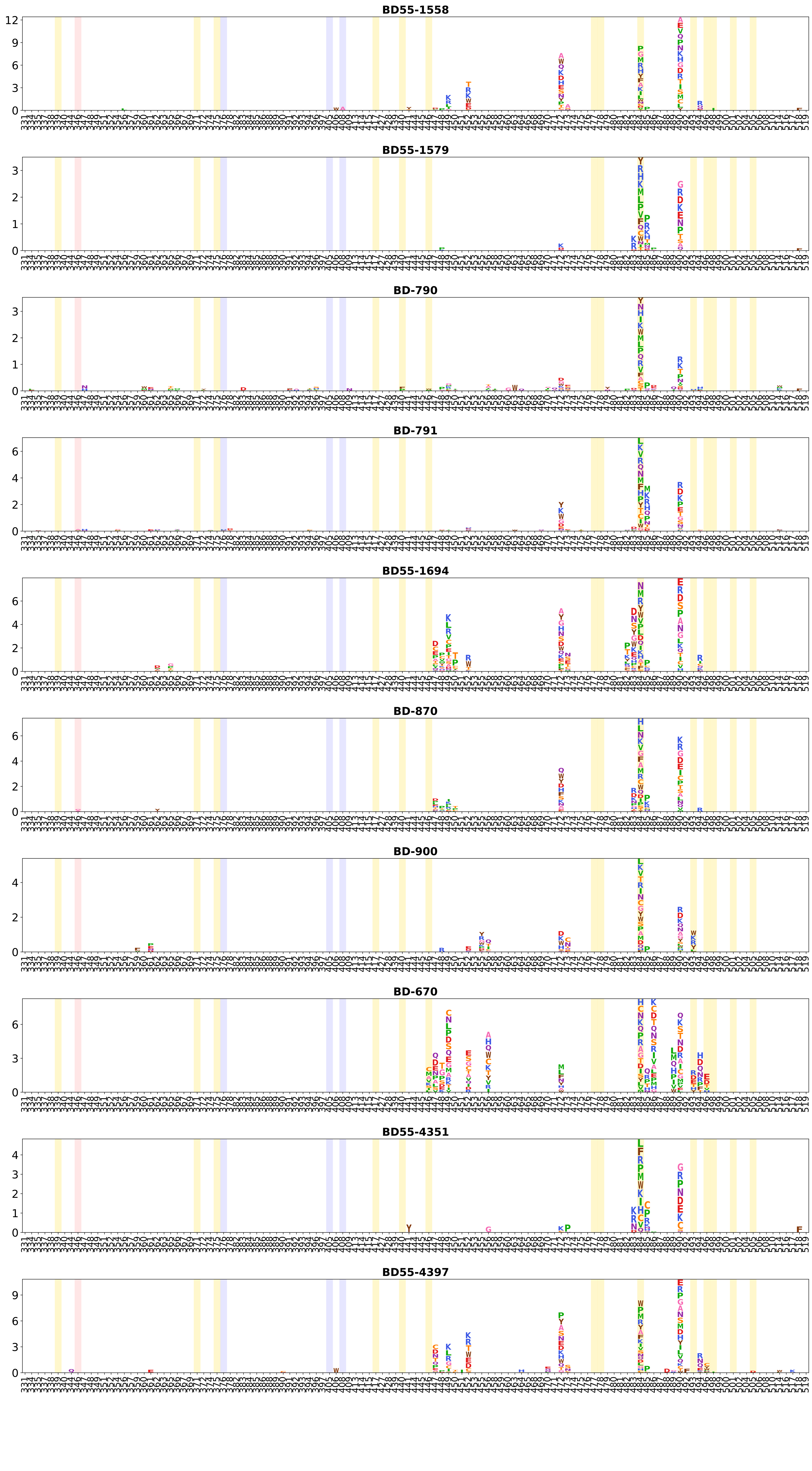
BD55-1558
BD55-1579
BD-790
BD-791
BD55-1694
BD-870
BD-900
BD-670
BD55-4351
BD55-4397

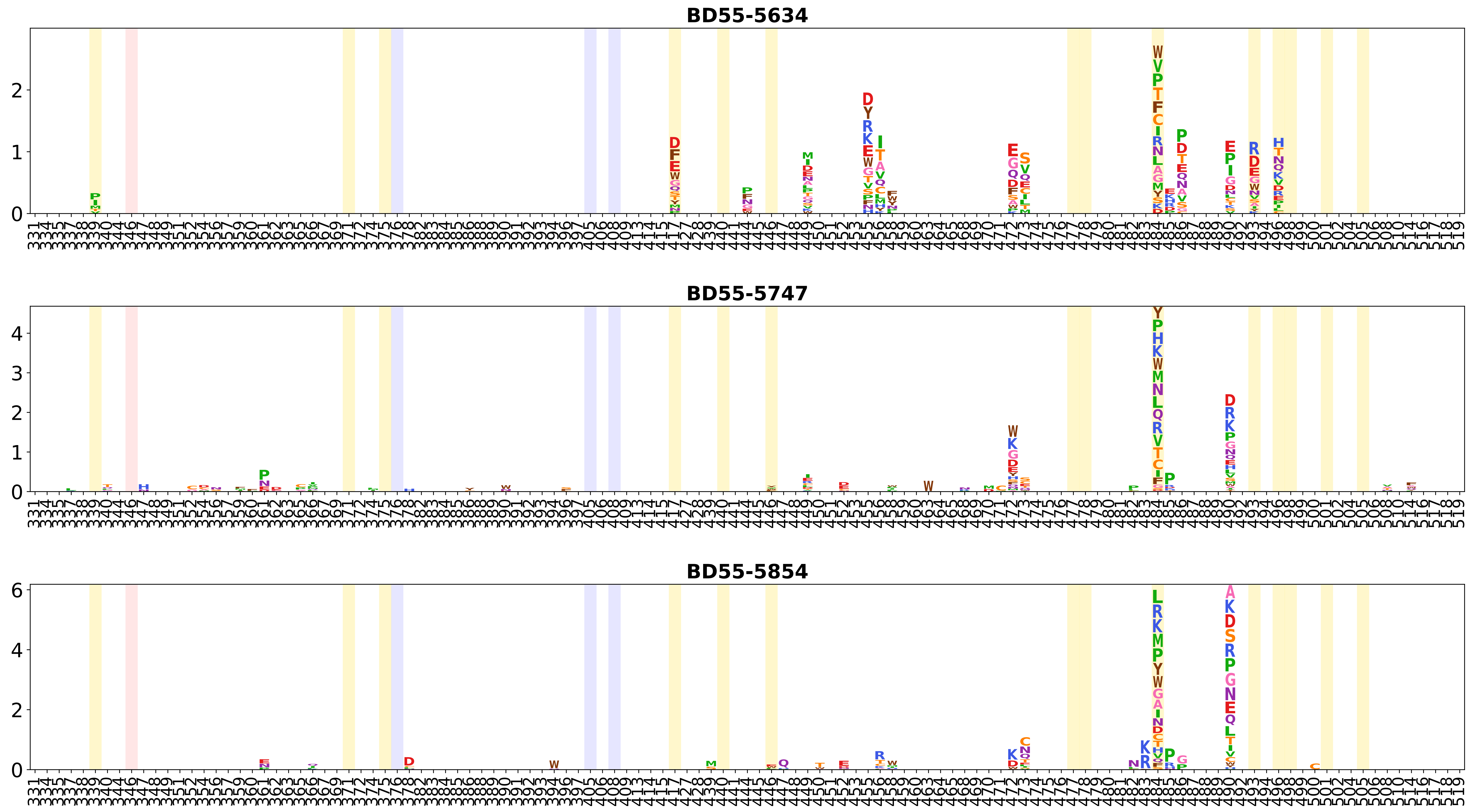
BD55-5634
BD55-5747
BD55-5854

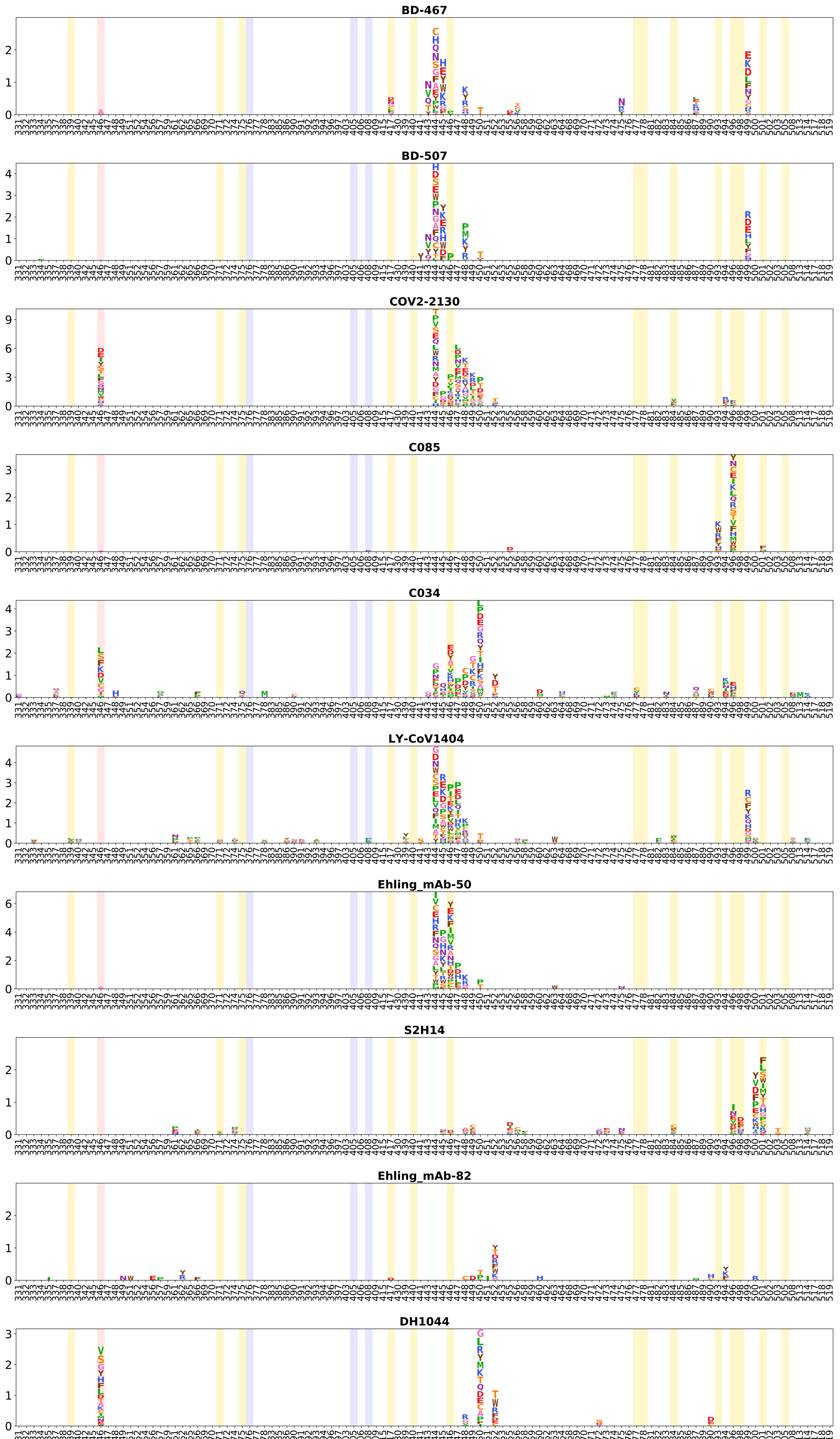
BD-467
BD-507
COV2-2130
C085
C034
LY-CoV1404
Ehling_mAb-50
S2H14
Ehling_mAb-82
DH1044

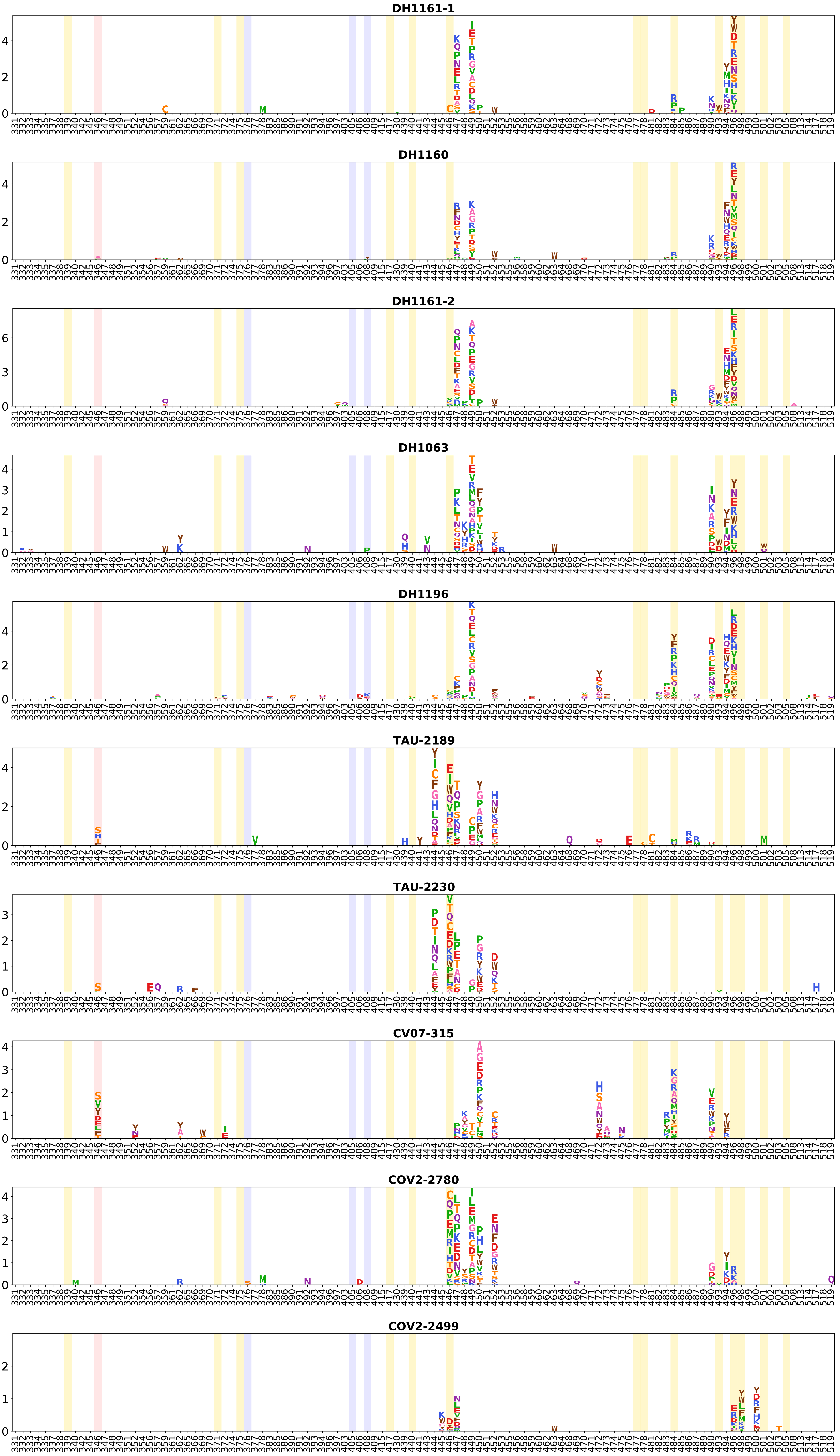
DH1161-1
DH1160
DH1161-2
DH1063
DH1196
TAU-2189
TAU-2230
CV07-315
COV2-2780
COV2-2499

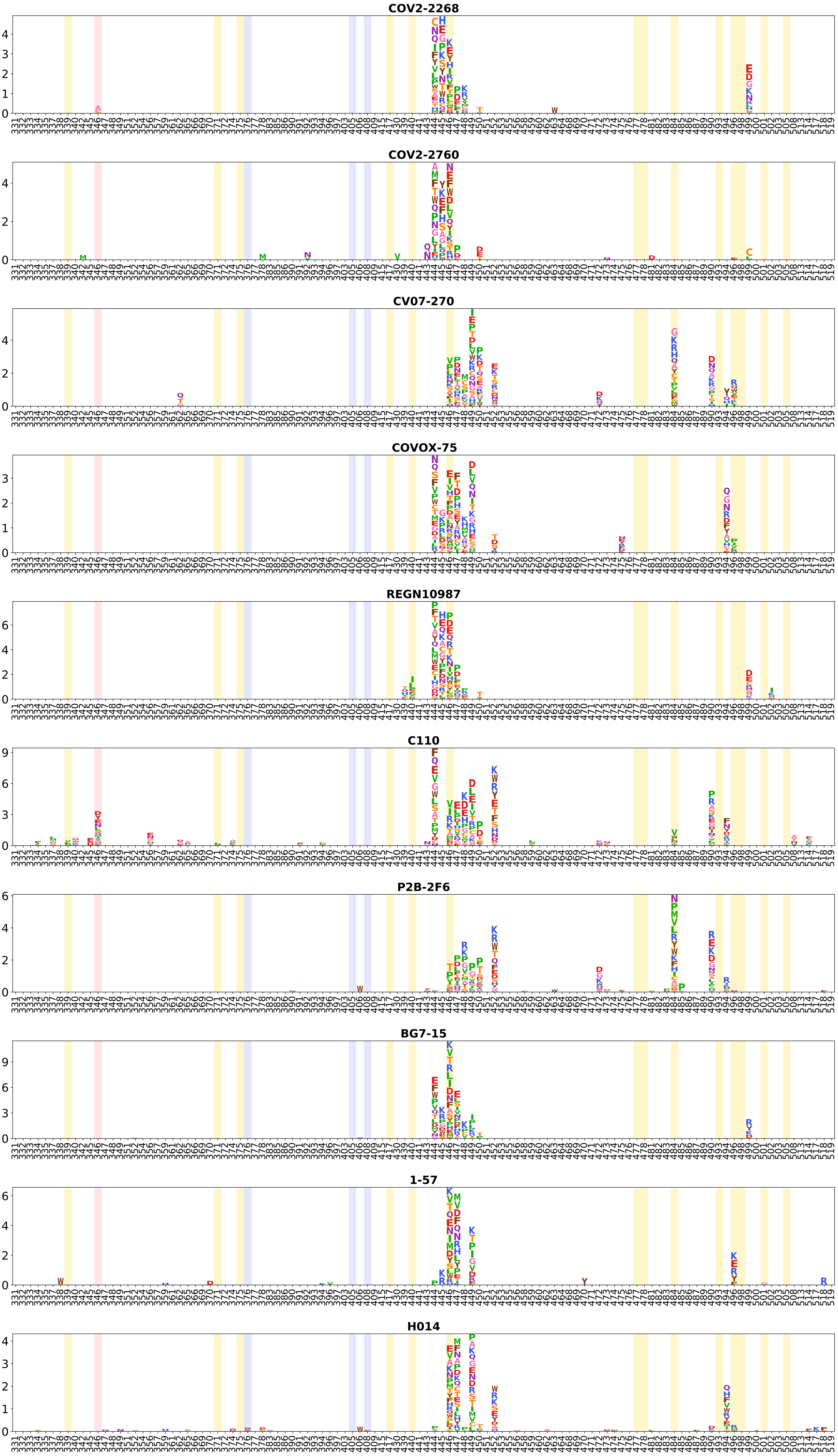
COV2-2268
COV2-2760
CV07-270
COVOX-75
REGN10987
C110
P2B-2F6
BG7-15
1-57
H014

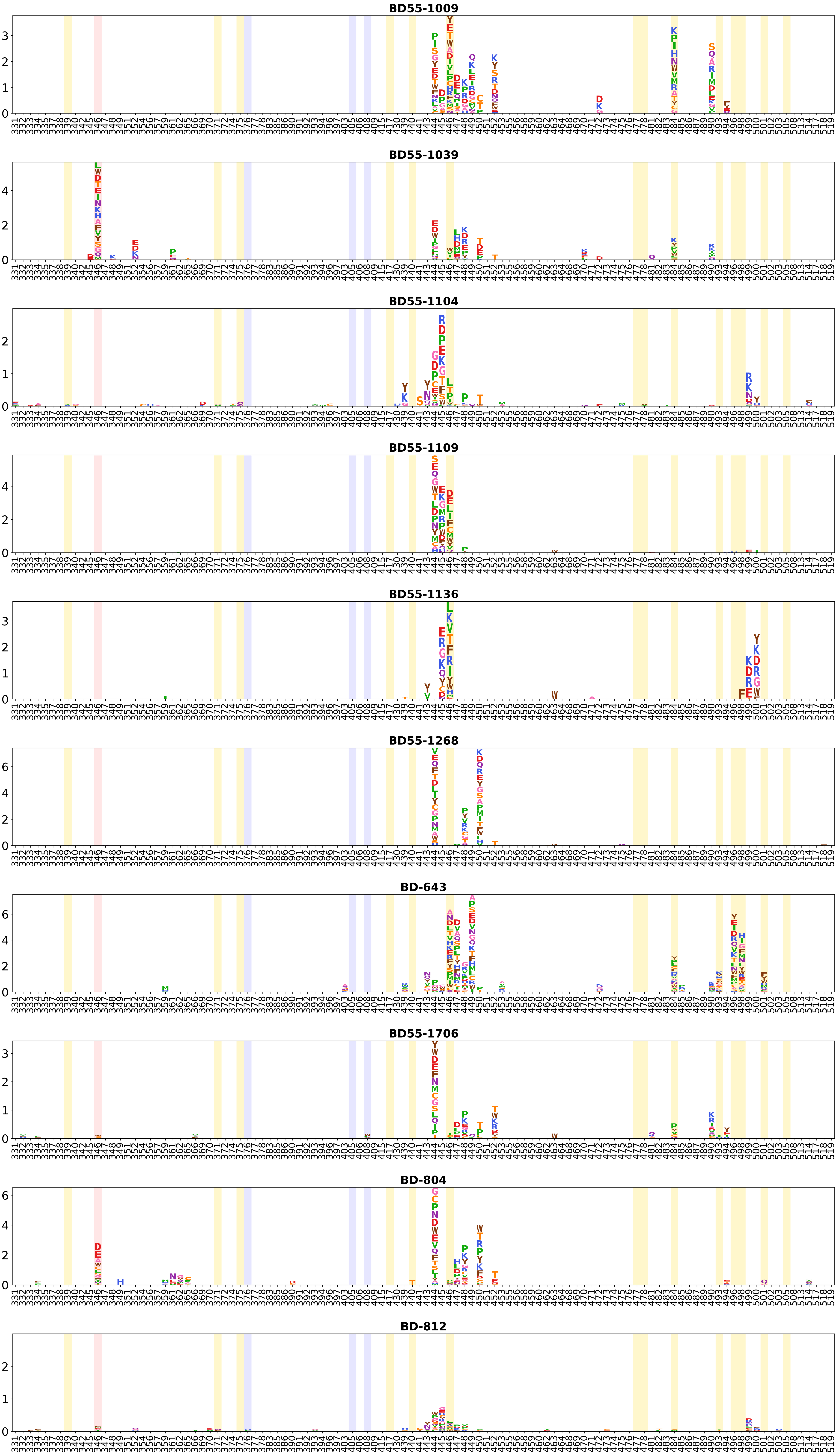
BD55-1009
BD55-1039
BD55-1104
BD55-1109
BD55-1136
BD55-1268
BD-643
BD55-1706
BD-804
BD-812

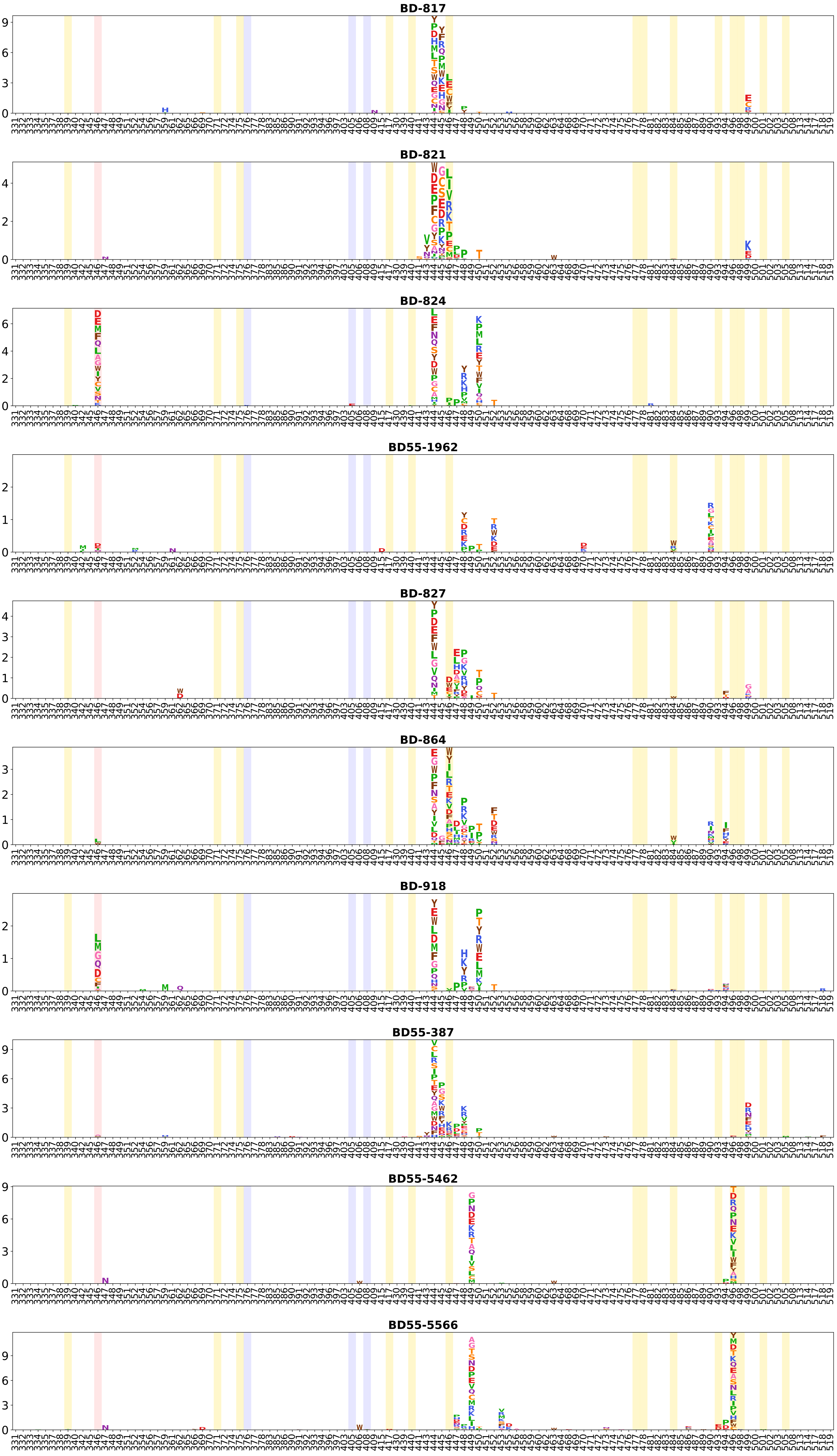
BD-817
BD-821
BD-824
BD55-1962
BD-827
BD-864
BD-918
BD55-387
BD55-5462
BD55-5566

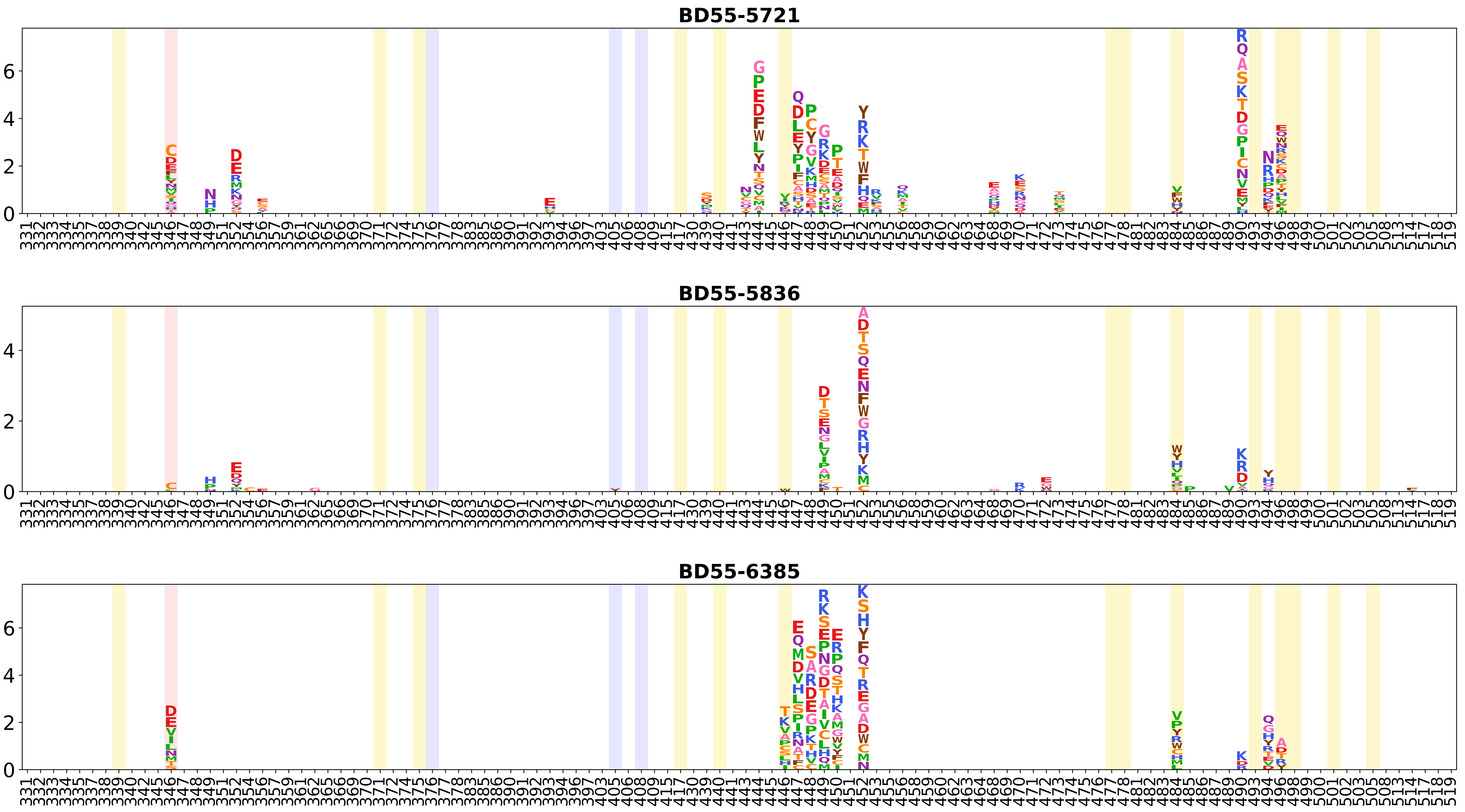
BD55-5721
BD55-5836
BD55-6385

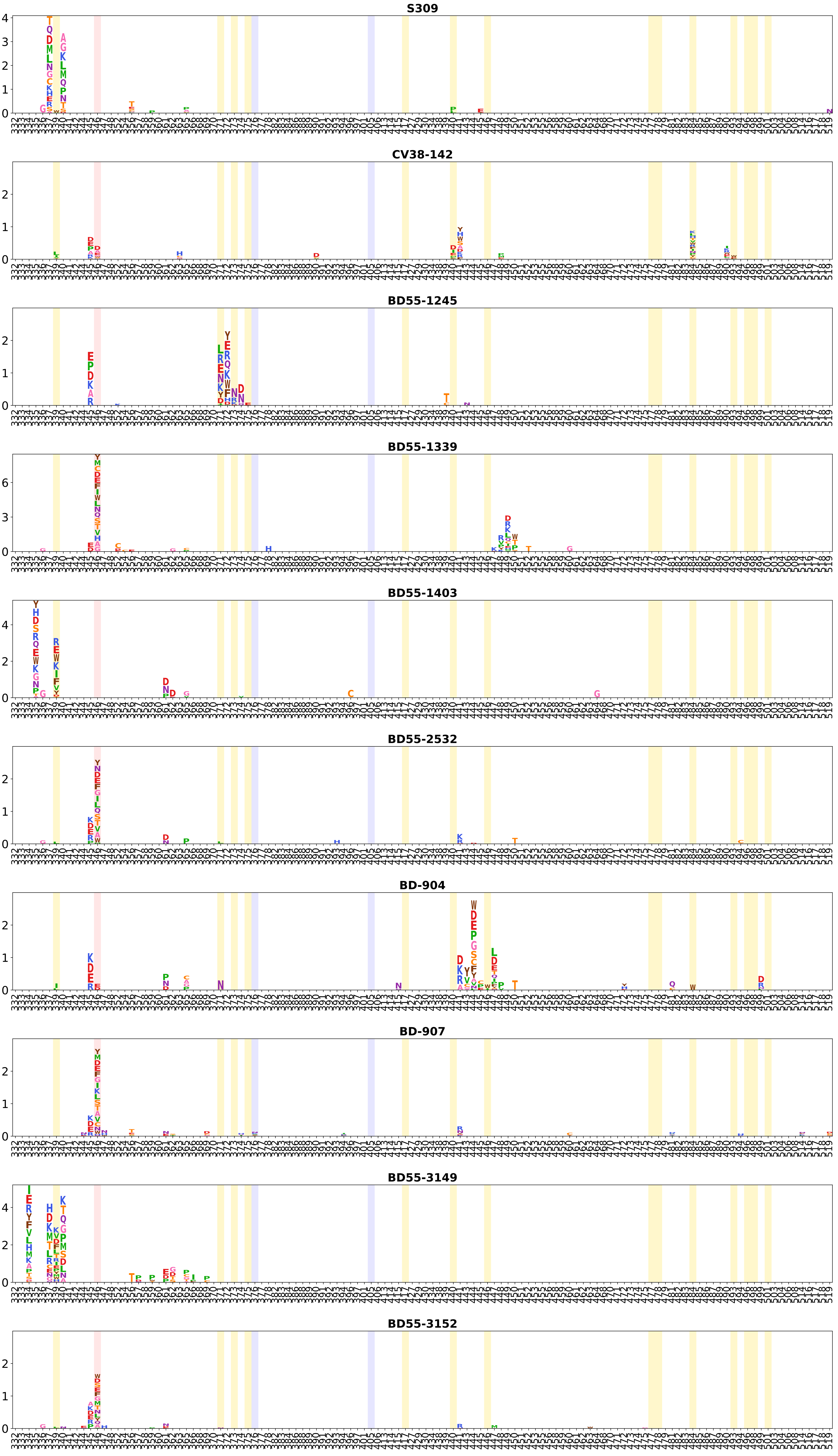
S309
CV38-142
BD55-1245
BD55-1339
BD55-1403
BD55-2532
BD-904
BD-907
BD55-3149
BD55-3152

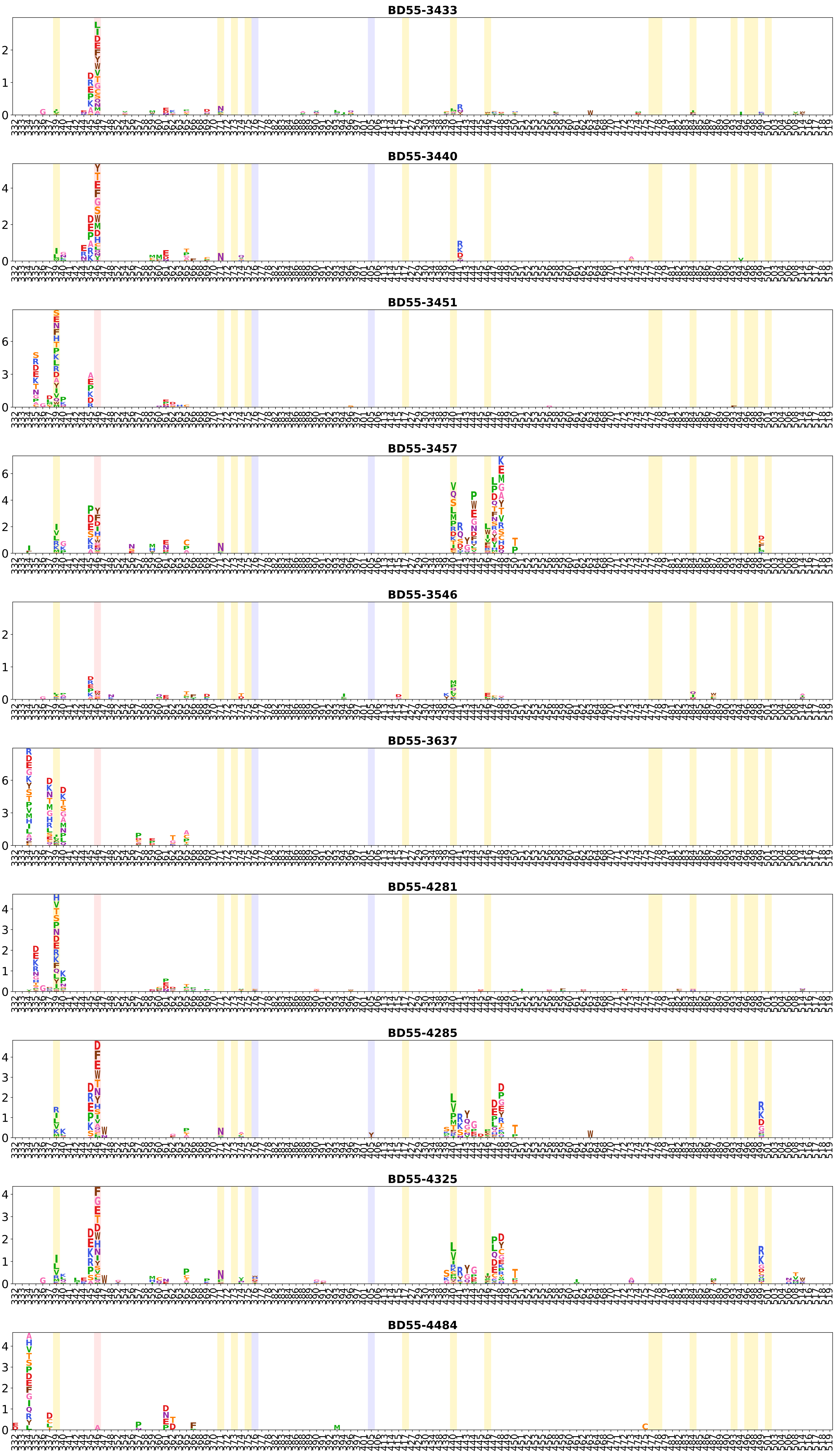
BD55-3433
BD55-3440
BD55-3451
BD55-3457
BD55-3546
BD55-3637
BD55-4281
BD55-4285
BD55-4325
BD55-4484

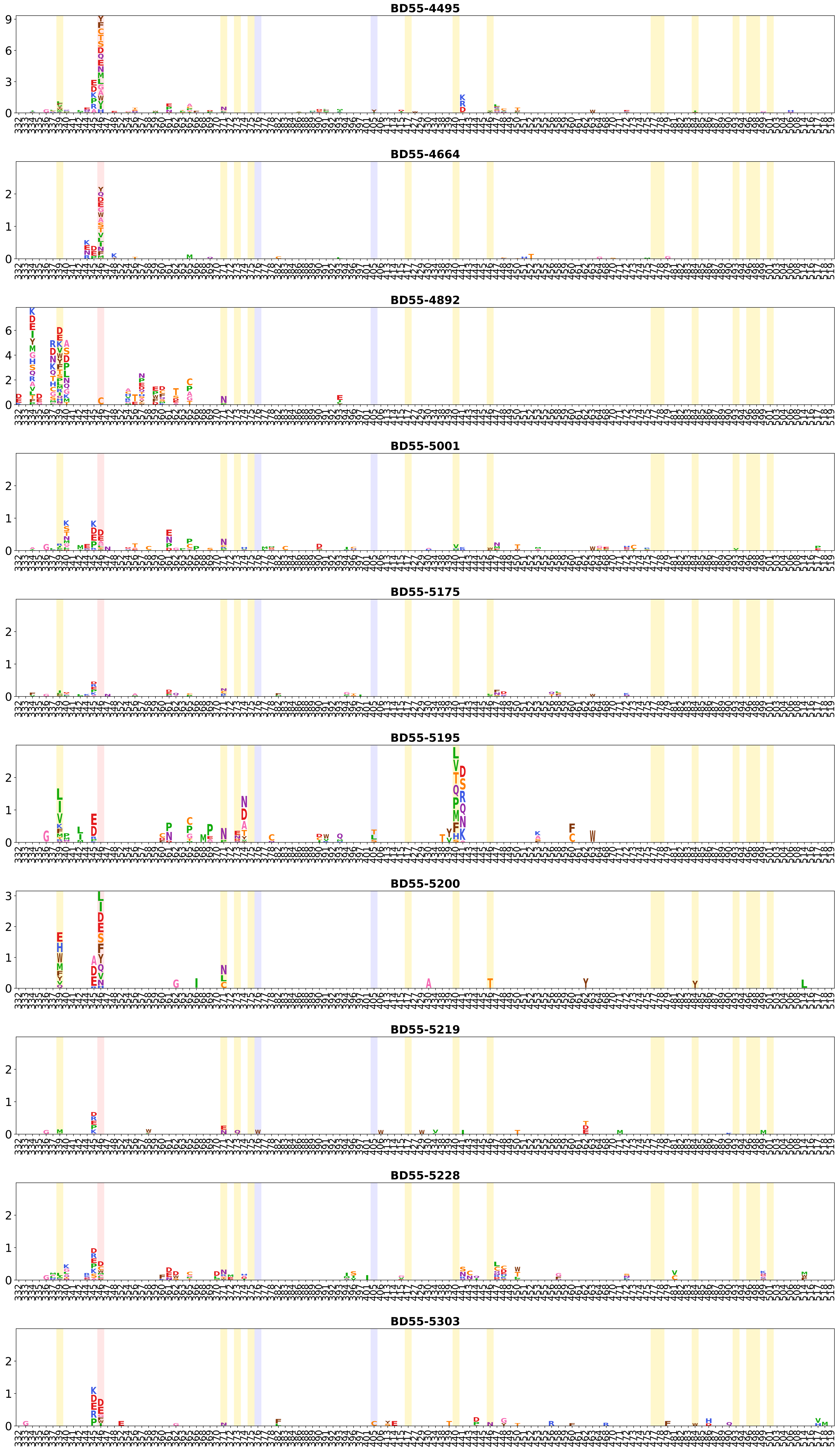
BD55-4495
BD55-4664
BD55-4892
BD55-5001
BD55-5175
BD55-5195
BD55-5200
BD55-5219
BD55-5228
BD55-5303

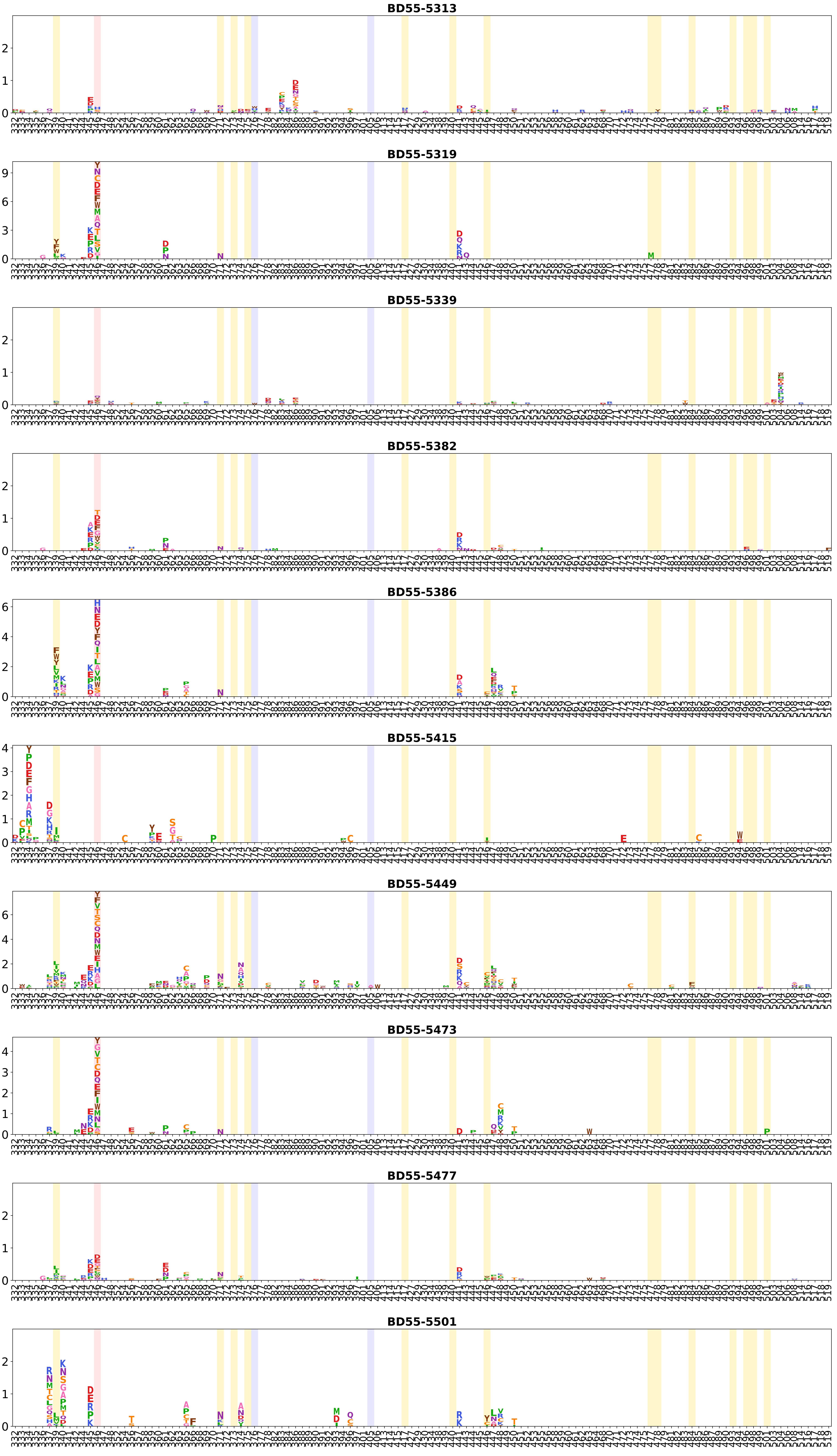
BD55-5313
BD55-5319
BD55-5339
BD55-5382
BD55-5386
BD55-5415
BD55-5449
BD55-5473
BD55-5477
BD55-5501

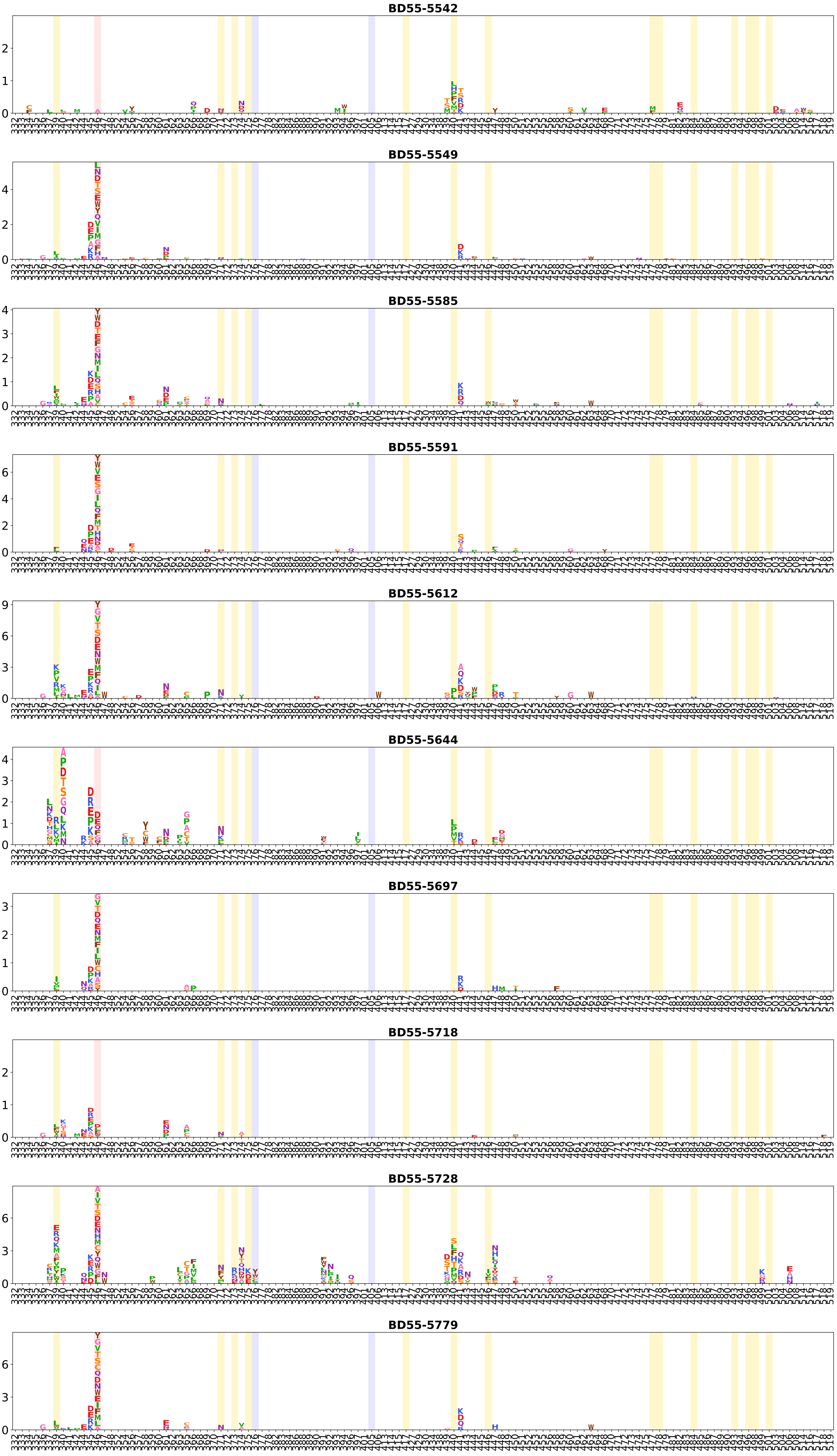
BD55-5542
BD55-5549
BD55-5585
BD55-5591
BD55-5612
BD55-5644
BD55-5697
BD55-5718
BD55-5728
BD55-5779

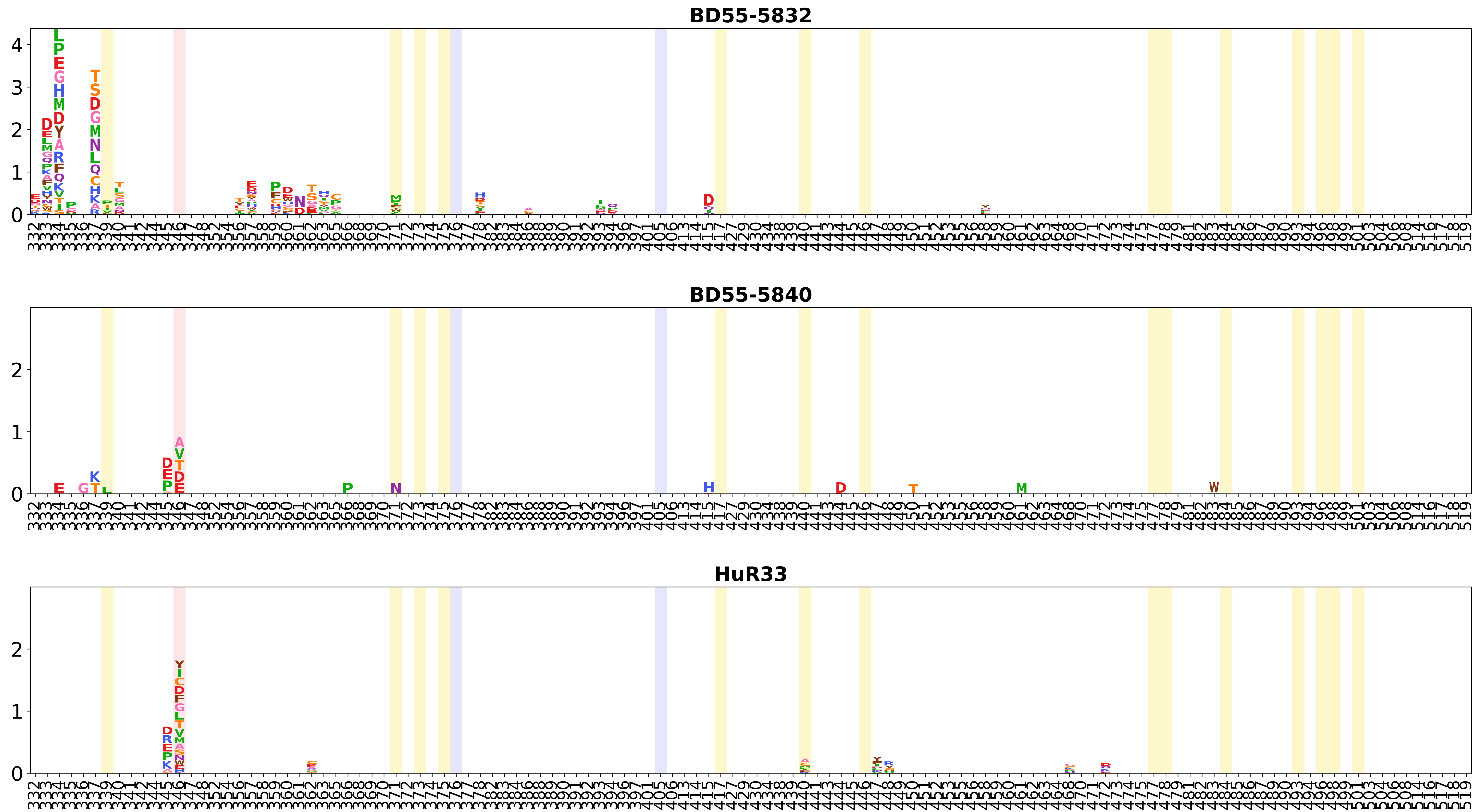
BD55-5832
BD55-5840
HuR33

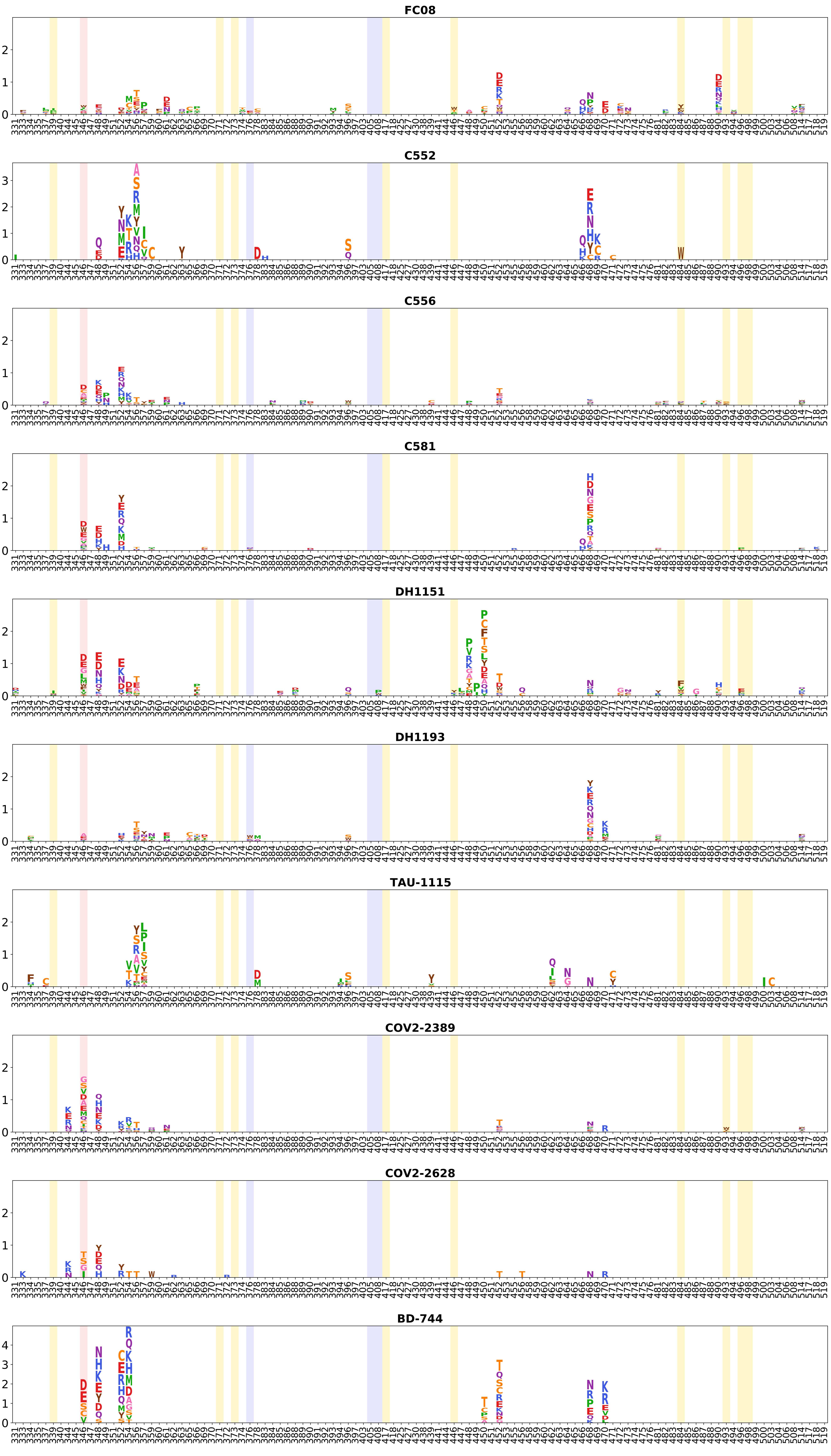
FC08
C552
C556
C581
DH1151
DH1193
TAU-1115
COV2-2389
COV2-2628
BD-744

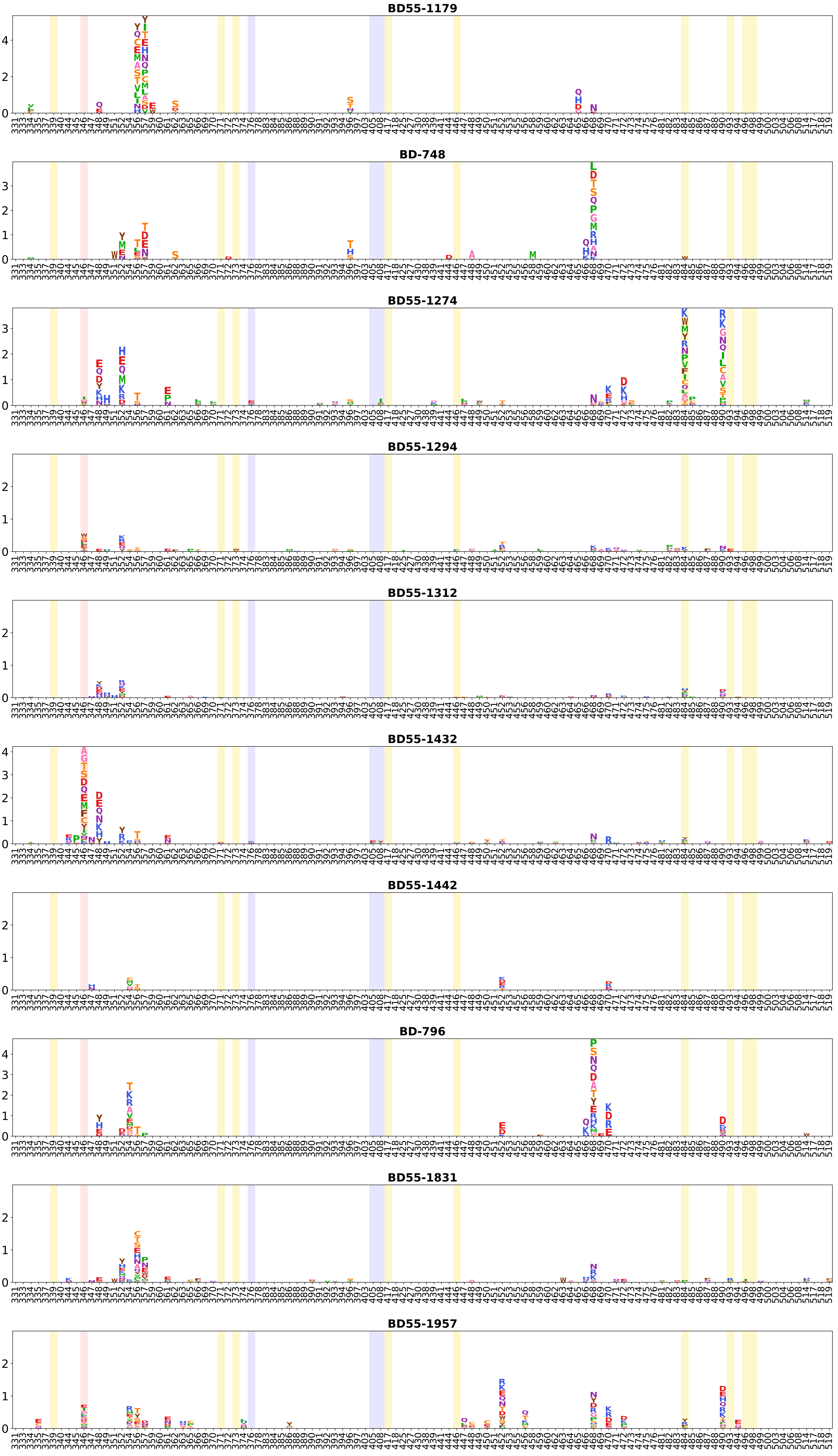
BD55-1179
BD-748
BD55-1274
BD55-1294
BD55-1312
BD55-1432
BD55-1442
BD-796
BD55-1831
BD55-1957

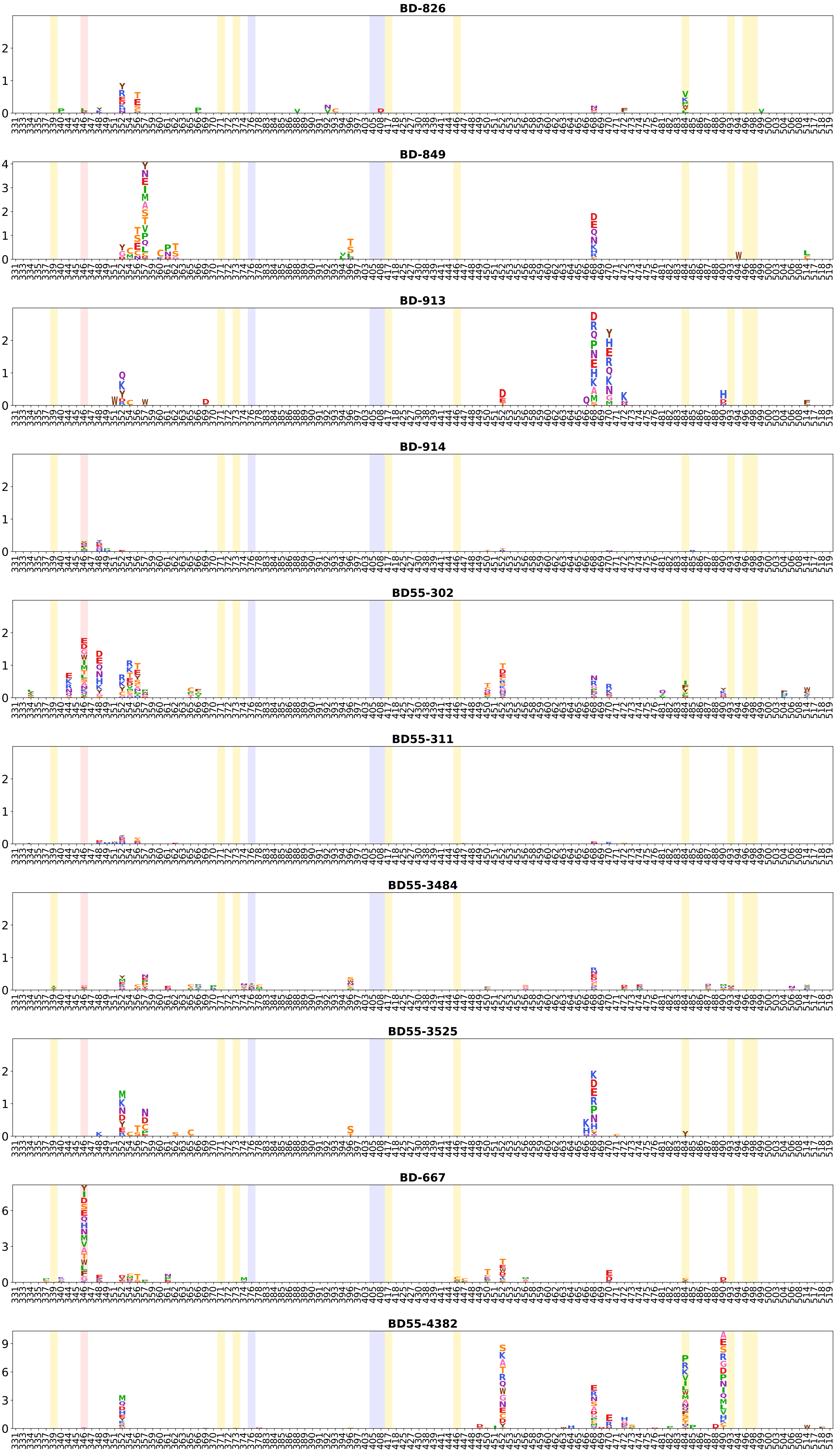
BD-826
BD-849
BD-913
BD-914
BD55-302
BD55-311
BD55-3484
BD55-3525
BD-667
BD55-4382

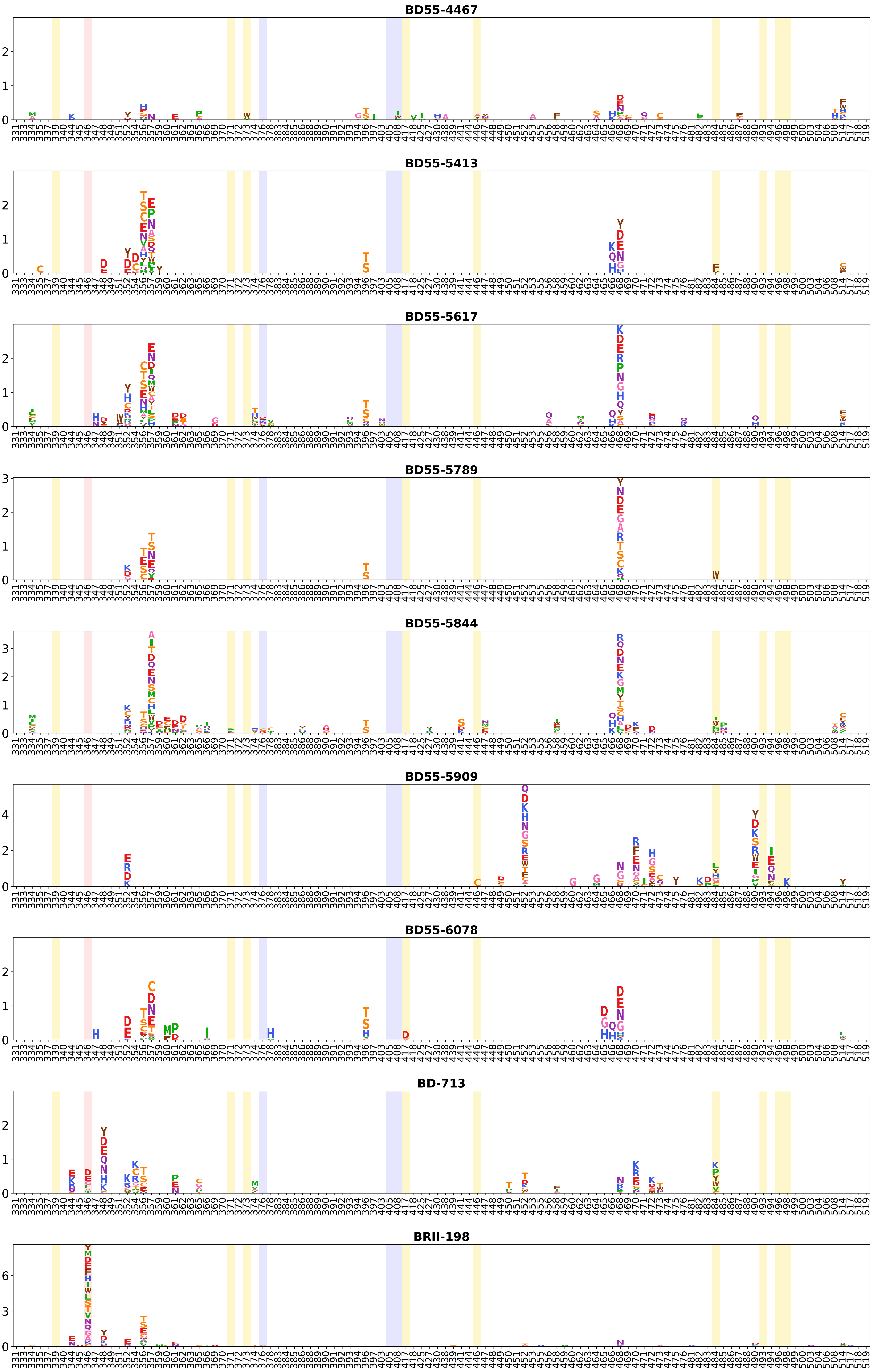
BD55-4467
BD55-5413
BD55-5617
BD55-5789
BD55-5844
BD55-5909
BD55-6078
BD-713
BRII-198

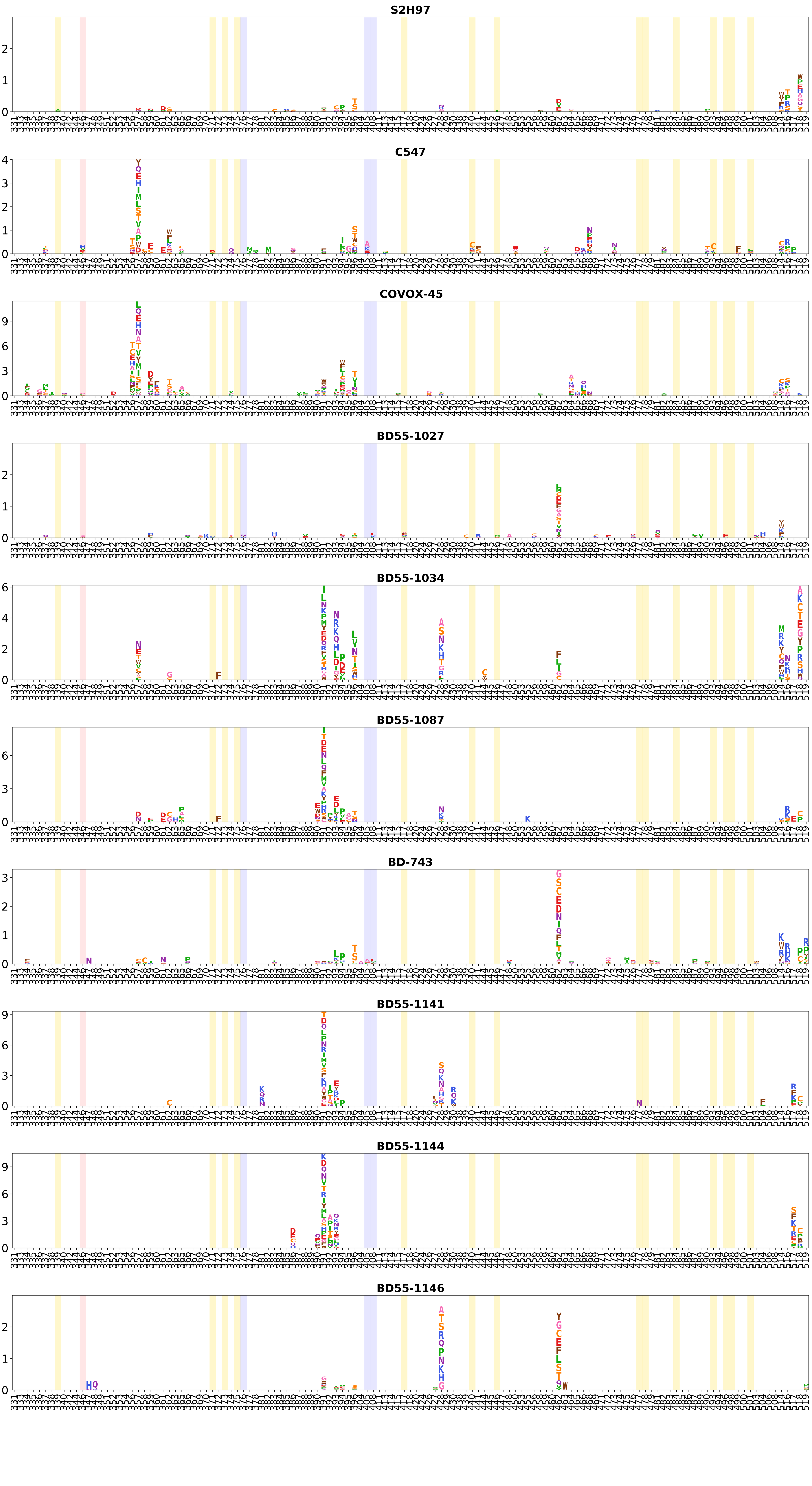
S2H97
C547
COVOX-45
BD55-1027
BD55-1034
BD55-1087
BD-743
BD55-1141
BD55-1144
BD55-1146

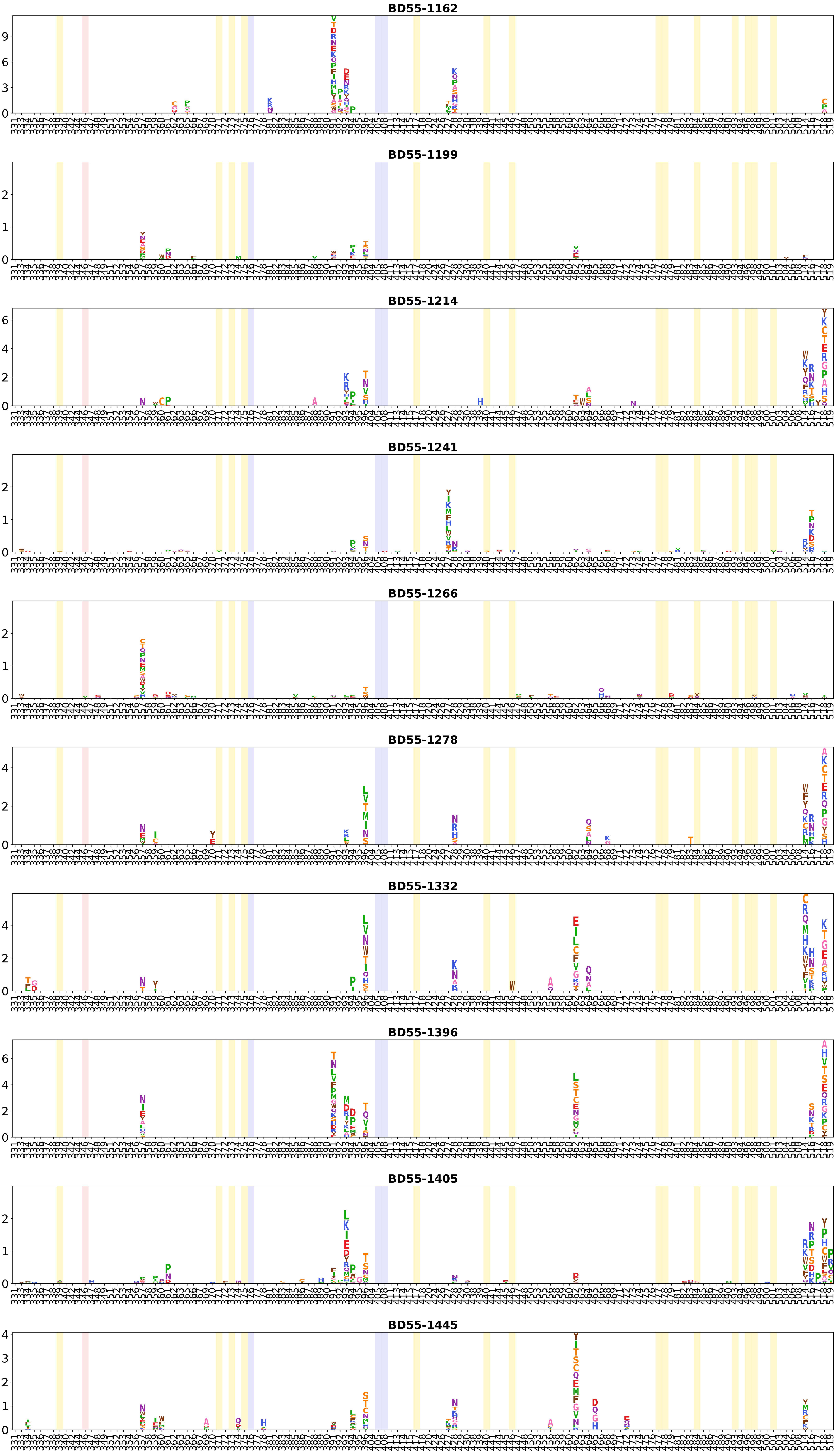
BD55-1162
BD55-1199
BD55-1214
BD55-1241
BD55-1266
BD55-1278
BD55-1332
BD55-1396
BD55-1405
BD55-1445

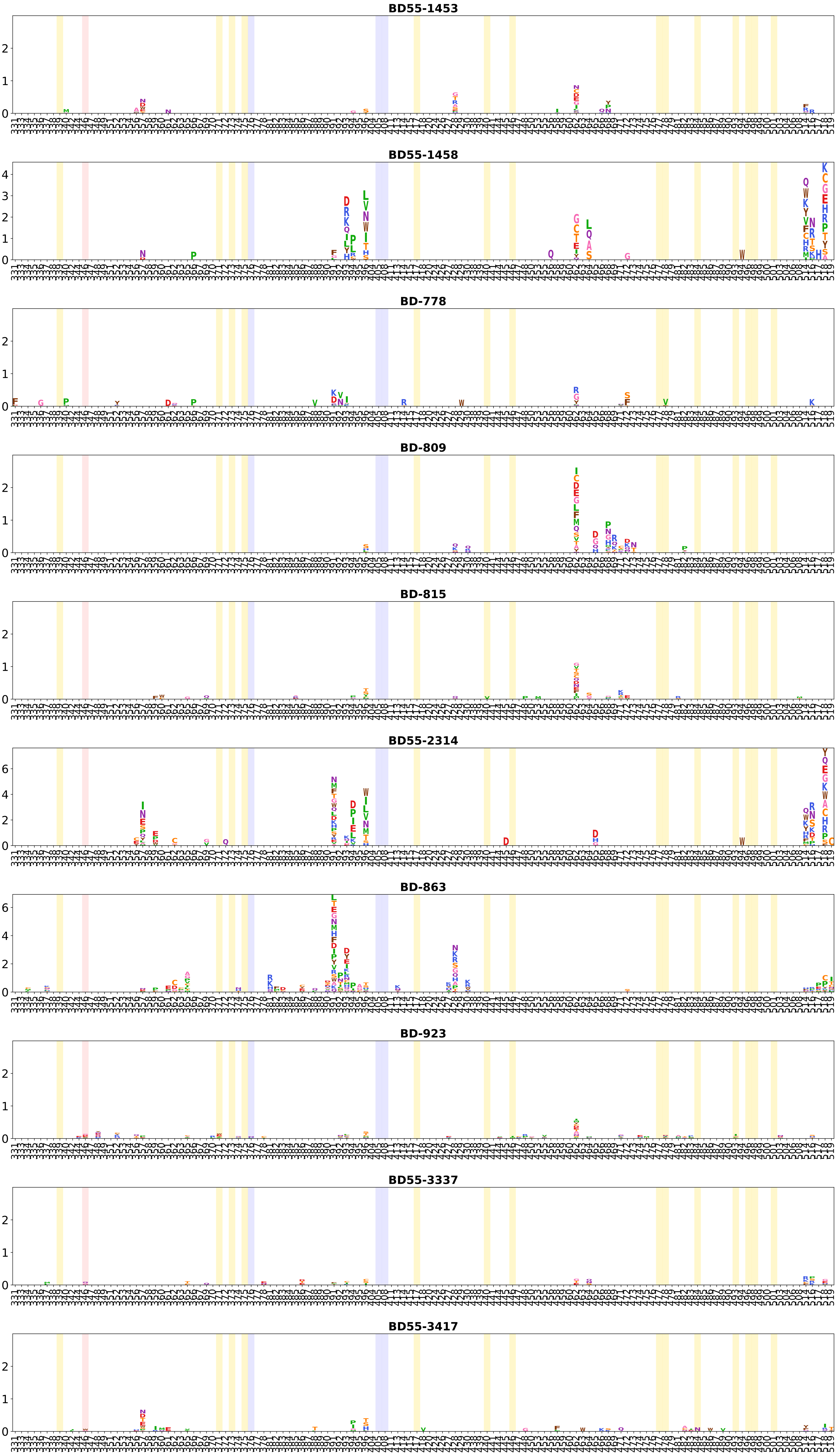
BD55-1453
BD55-1458
BD-778
BD-809
BD-815
BD55-2314
BD-863
BD-923
BD55-3337
BD55-3417

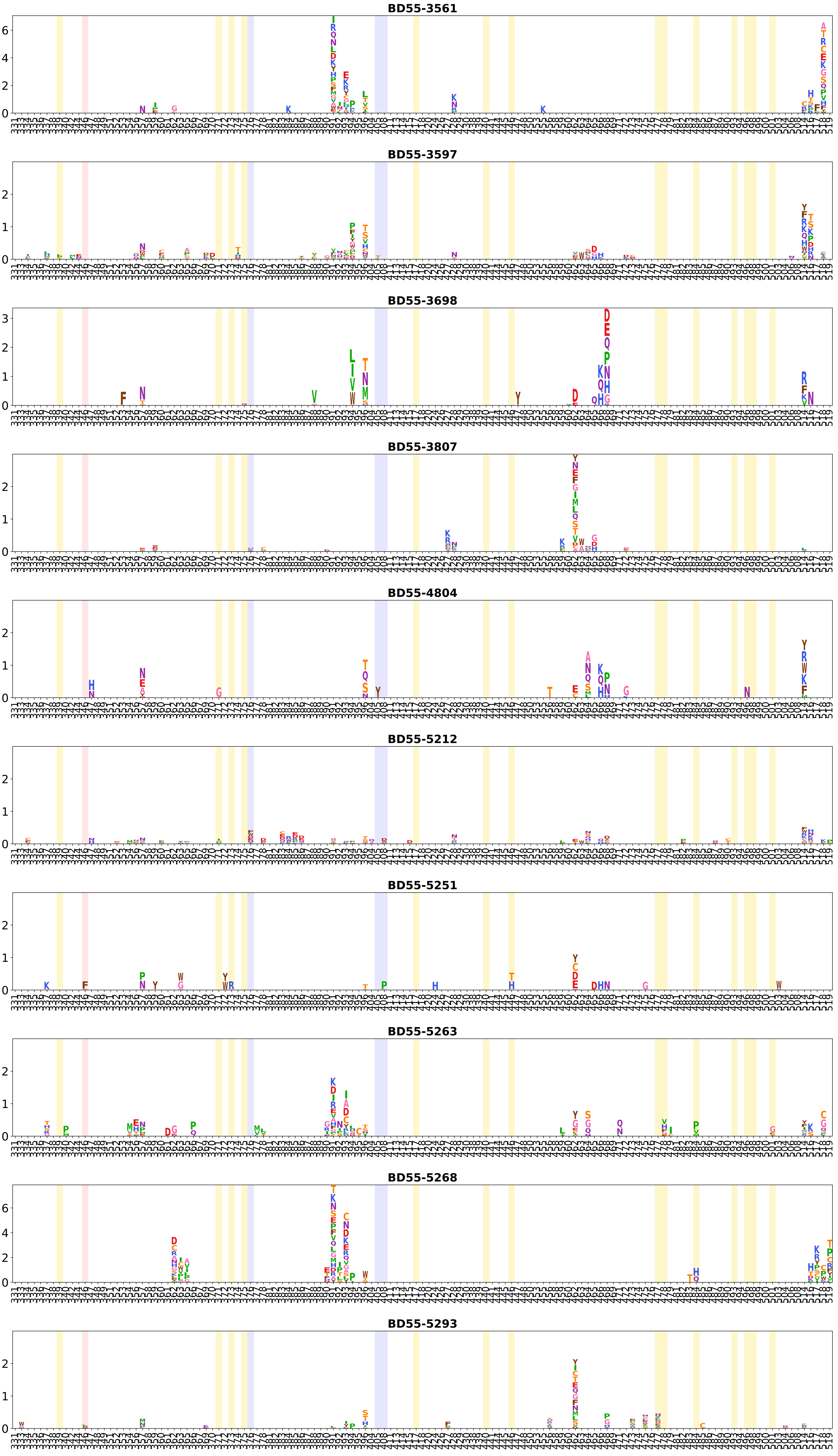
BD55-3561
BD55-3597
BD55-3698
BD55-3807
BD55-4804
BD55-5212
BD55-5251
BD55-5263
BD55-5268
BD55-5293

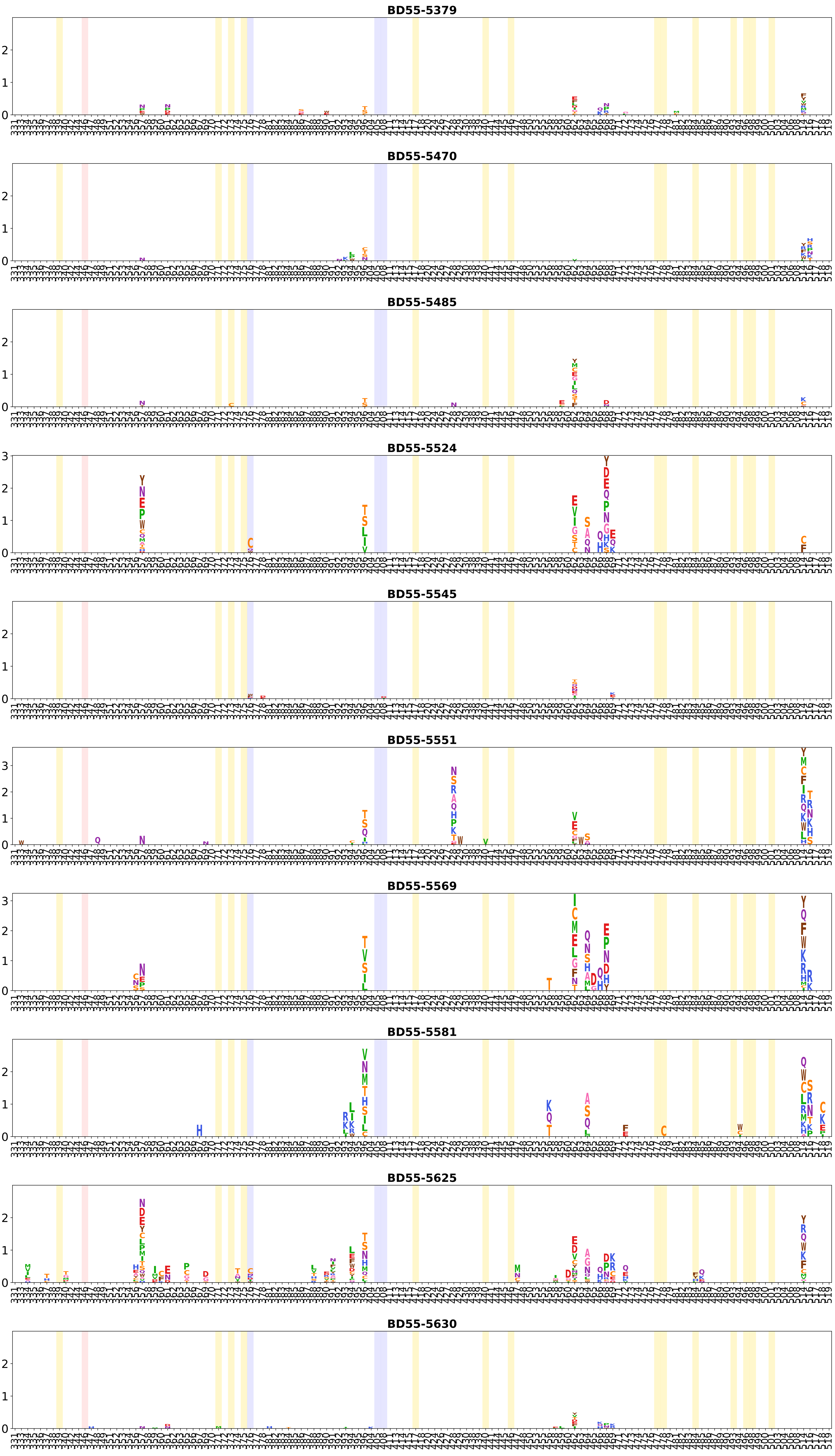
BD55-5379
BD55-5470
BD55-5485
BD55-5524
BD55-5545
BD55-5551
BD55-5569
BD55-5581
BD55-5625
BD55-5630

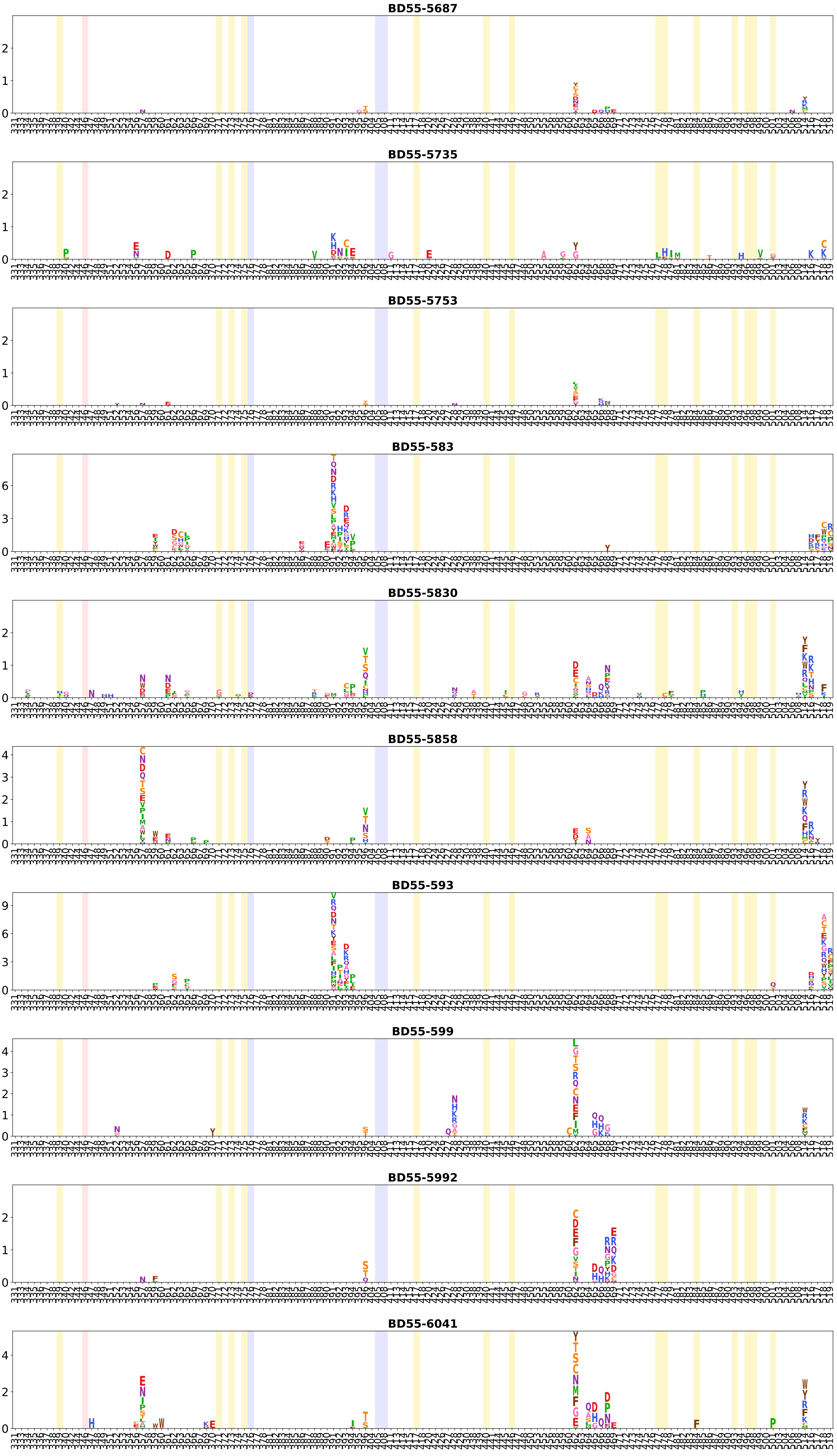
BD55-5687
BD55-5735
BD55-5753
BD55-583
BD55-5830
BD55-5858
BD55-593
BD55-599
BD55-5992
BD55-6041

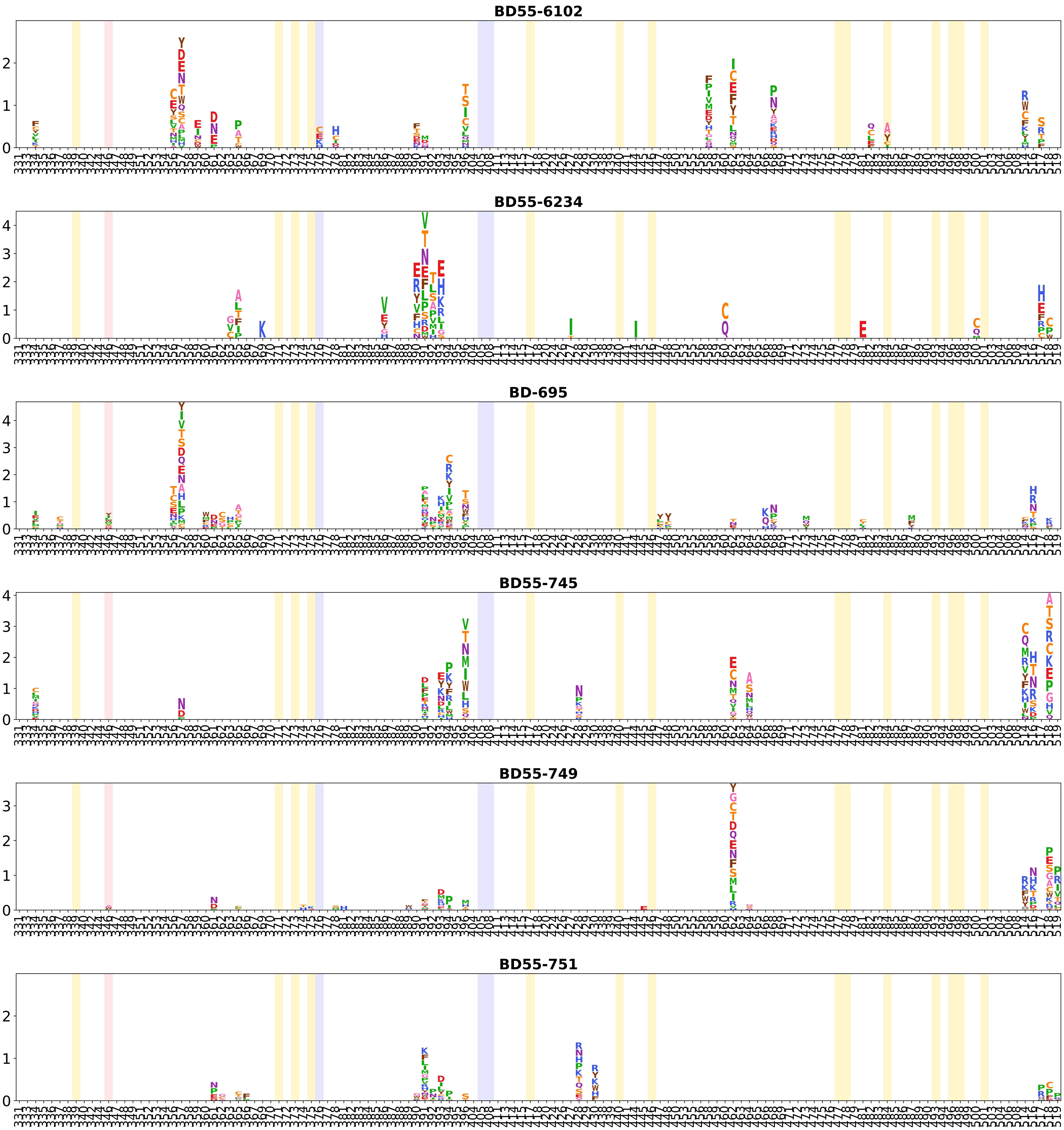
BD55-6102
BD55-6234
BD-695
BD55-745
BD55-749
BD55-751

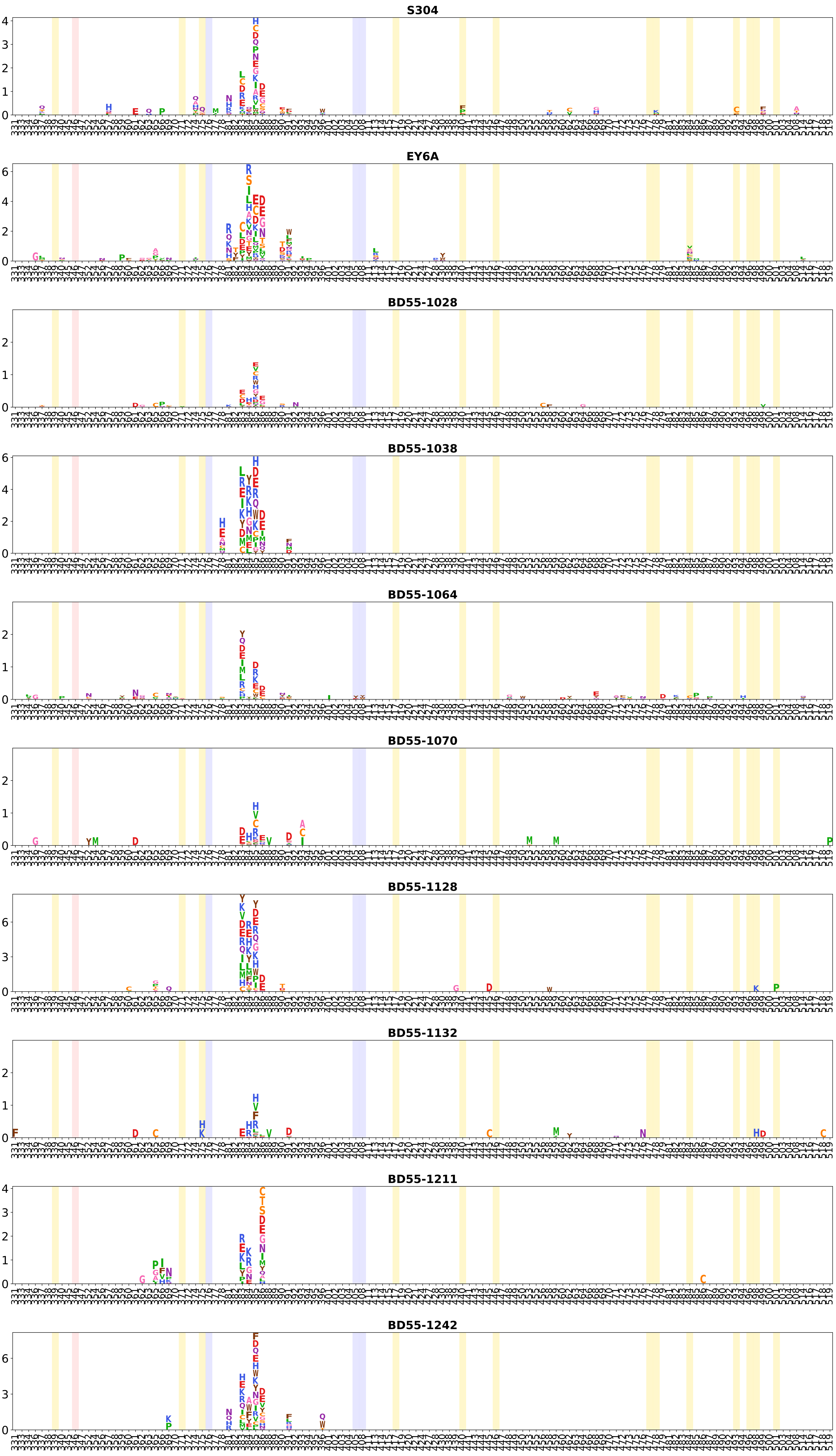
S304
EY6A
BD55-1028
BD55-1038
BD55-1064
BD55-1070
BD55-1128
BD55-1132
BD55-1211
BD55-1242

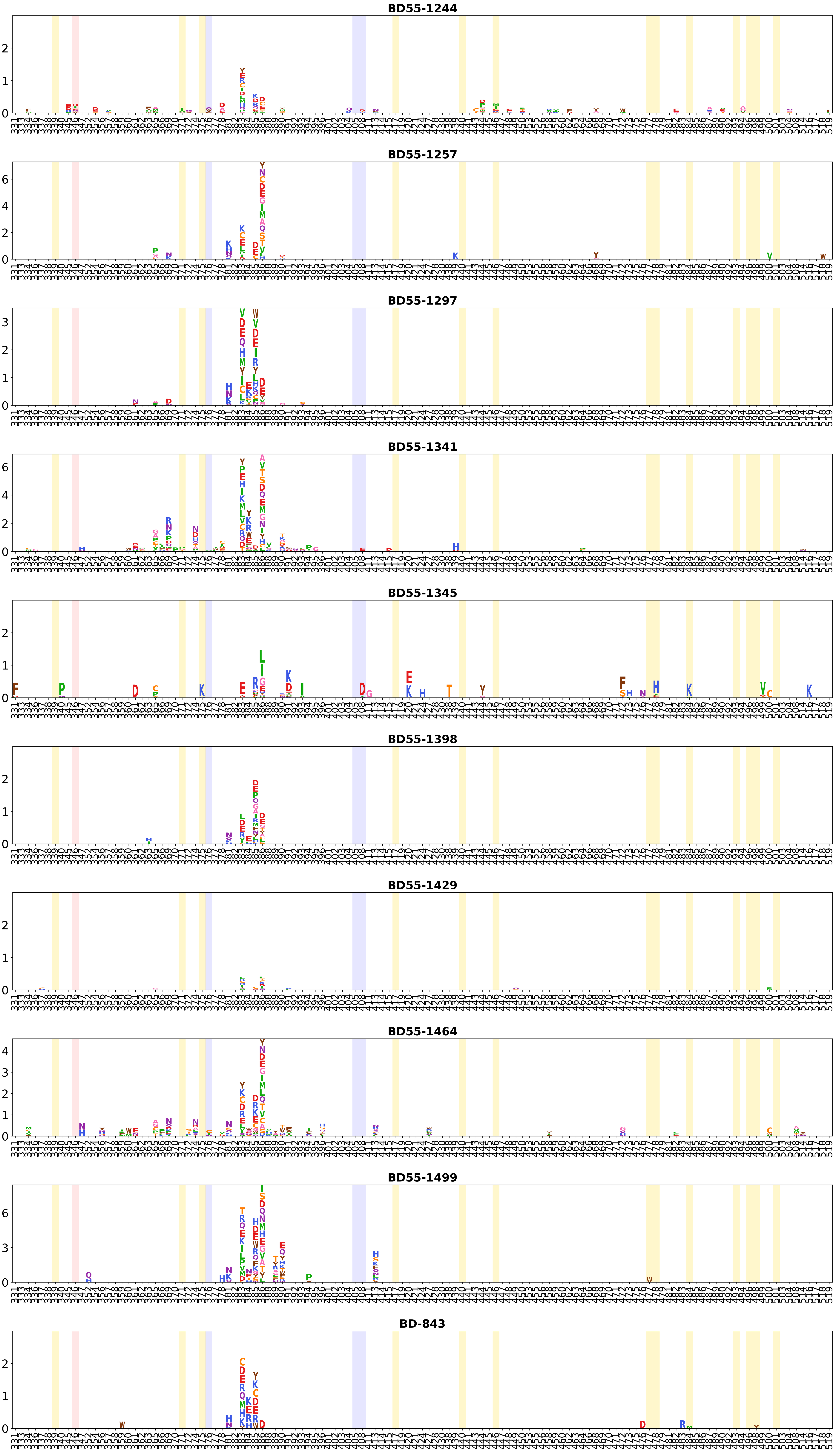
BD55-1244
BD55-1257
BD55-1297
BD55-1341
BD55-1345
BD55-1398
BD55-1429
BD55-1464
BD55-1499
BD-843

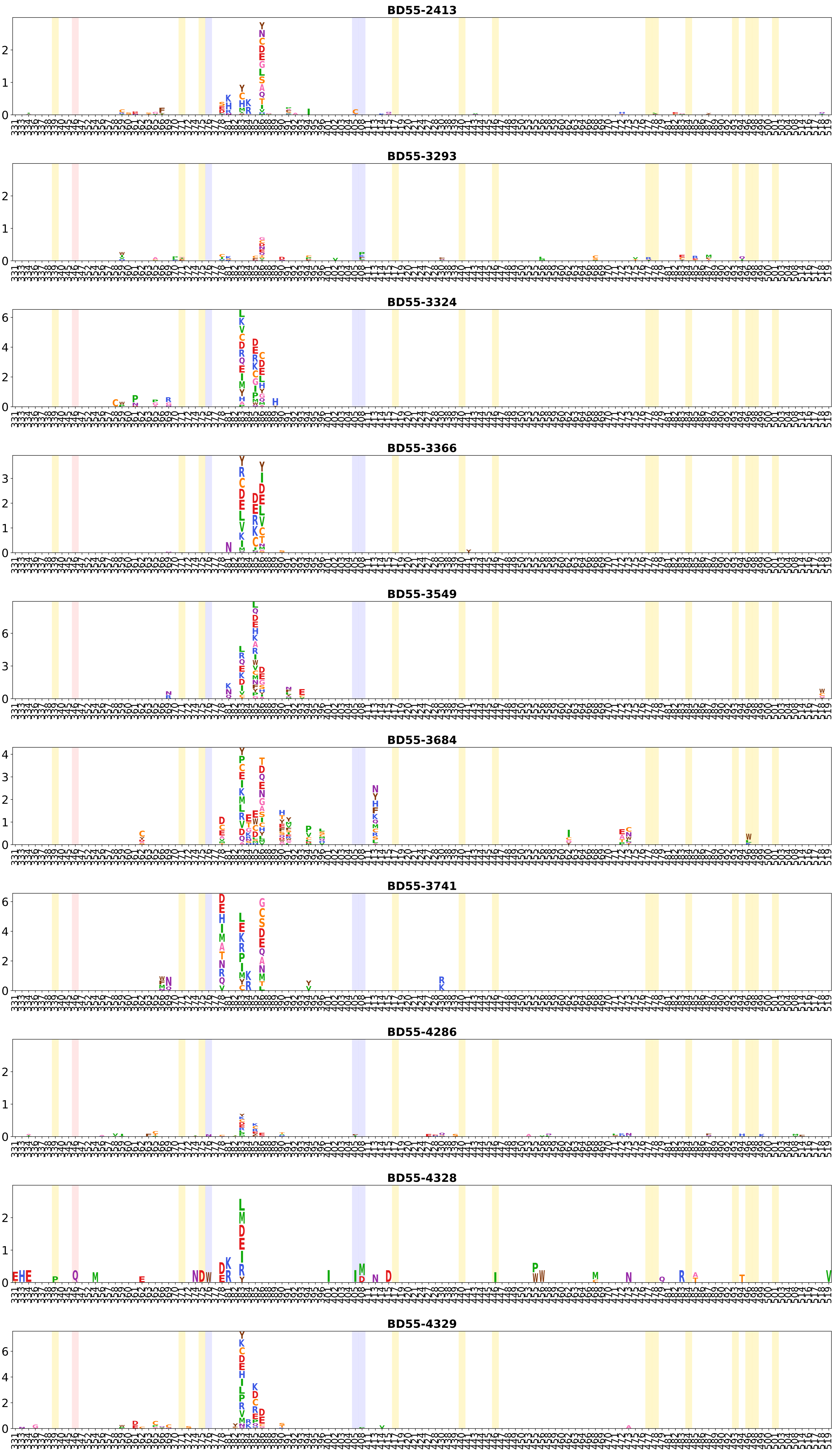
BD55-2413
BD55-3293
BD55-3324
BD55-3366
BD55-3549
BD55-3684
BD55-3741
BD55-4286
BD55-4328
BD55-4329

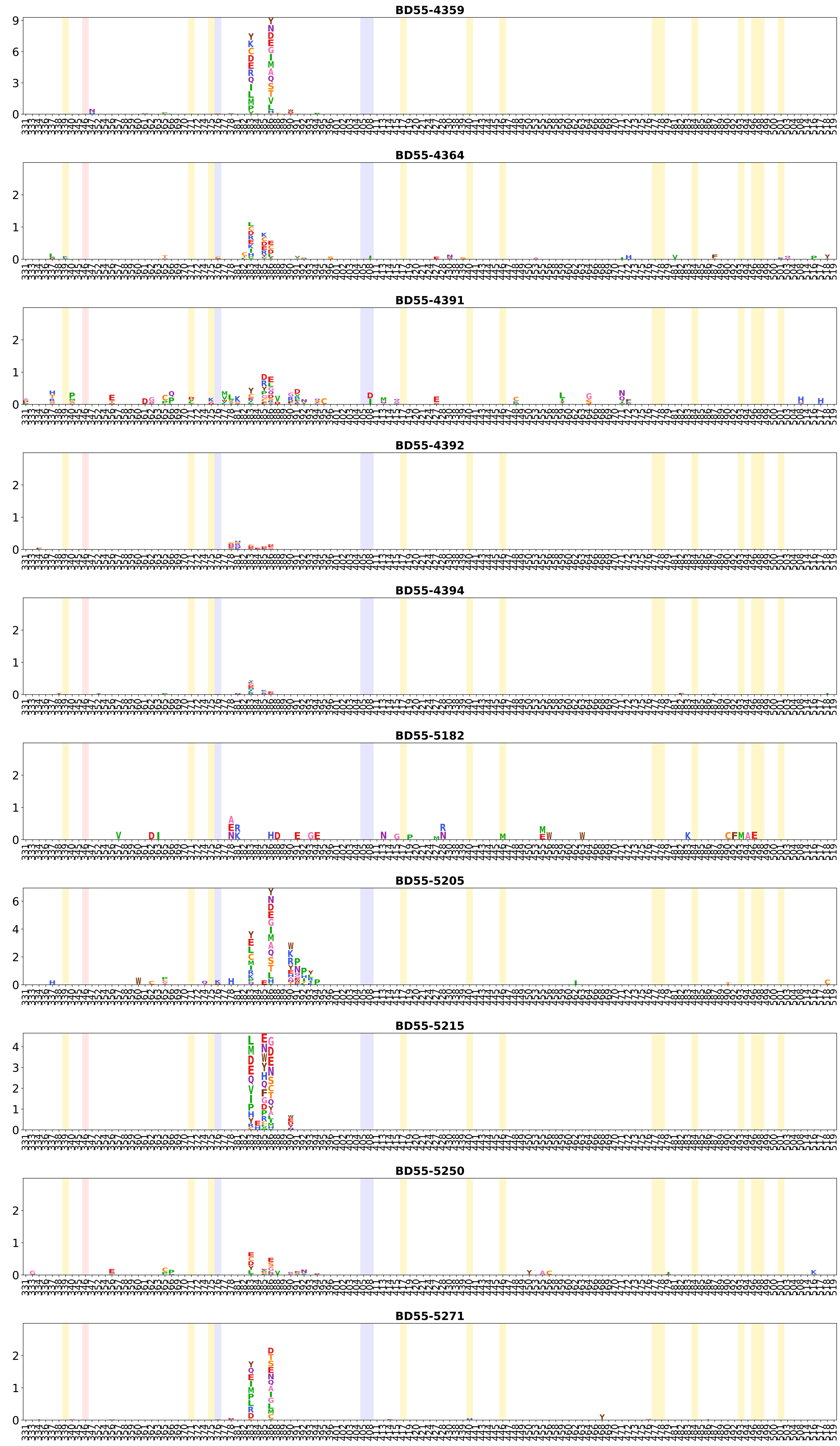
BD55-4359
BD55-4364
BD55-4391
BD55-4392
BD55-4394
BD55-5182
BD55-5205
BD55-5215
BD55-5250
BD55-5271

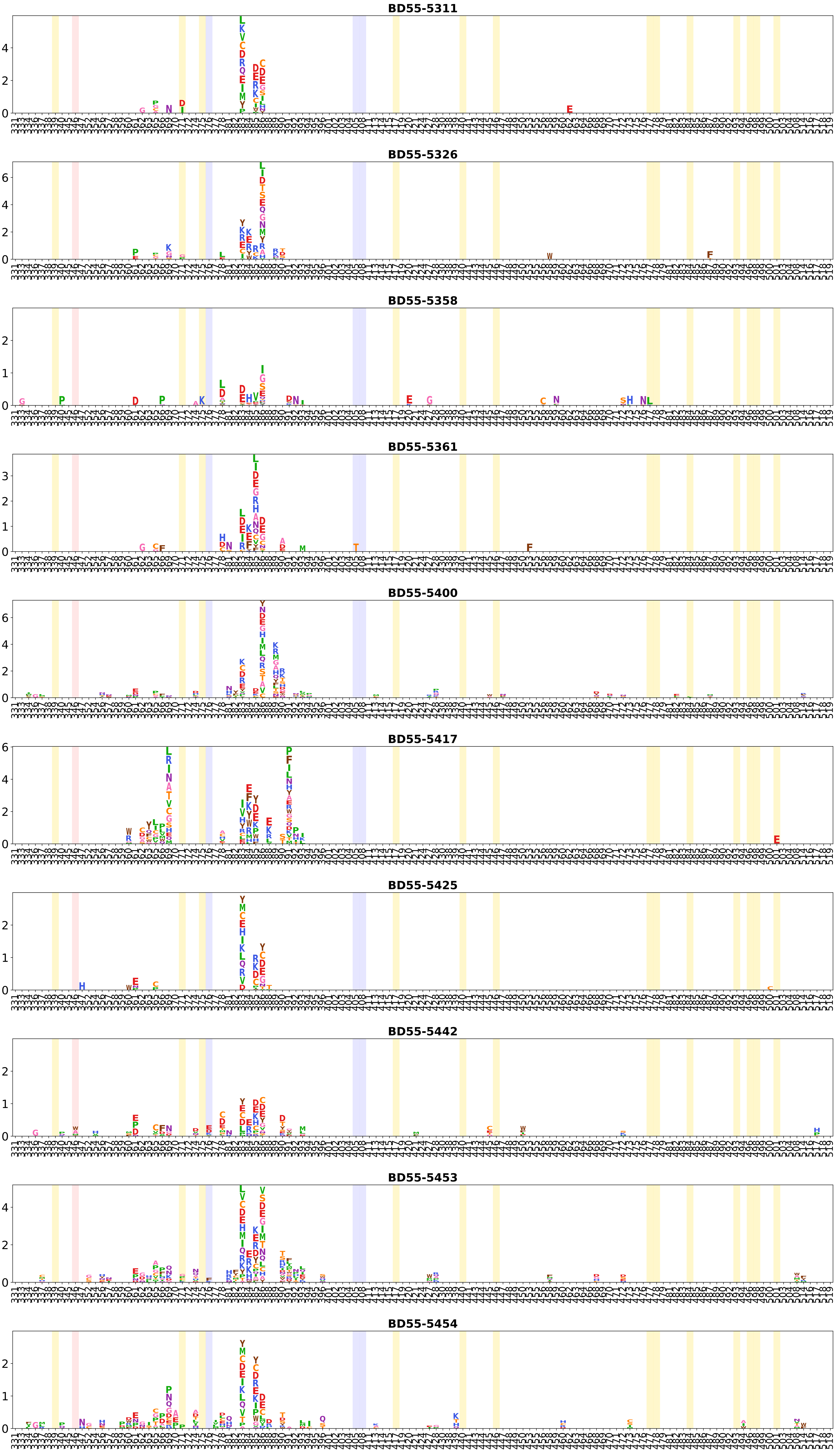
BD55-5311
BD55-5326
BD55-5358
BD55-5361
BD55-5400
BD55-5417
BD55-5425
BD55-5442
BD55-5453
BD55-5454

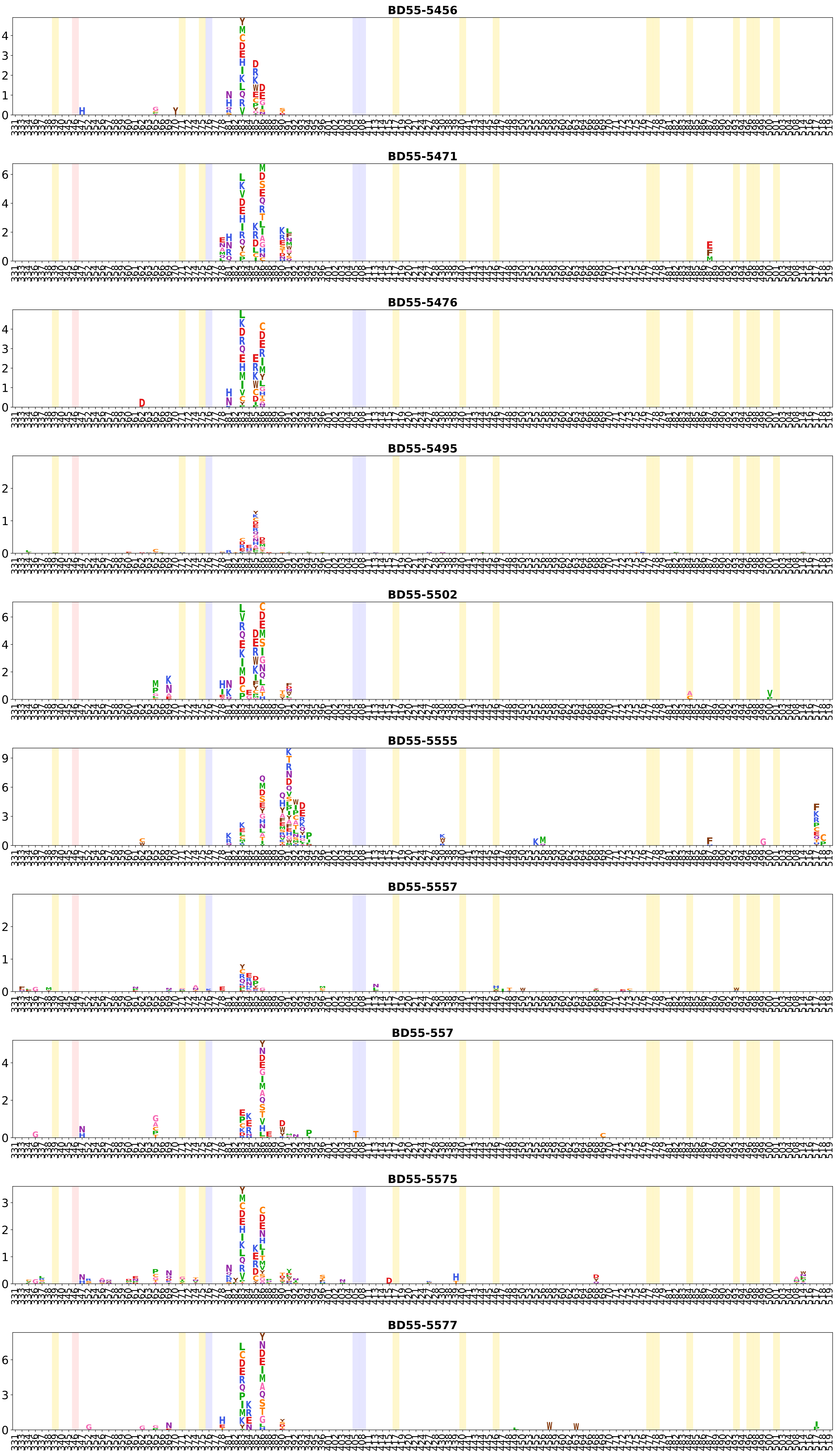
BD55-5456
BD55-5471
BD55-5476
BD55-5495
BD55-5502
BD55-5555
BD55-5557
BD55-557
BD55-5575
BD55-5577

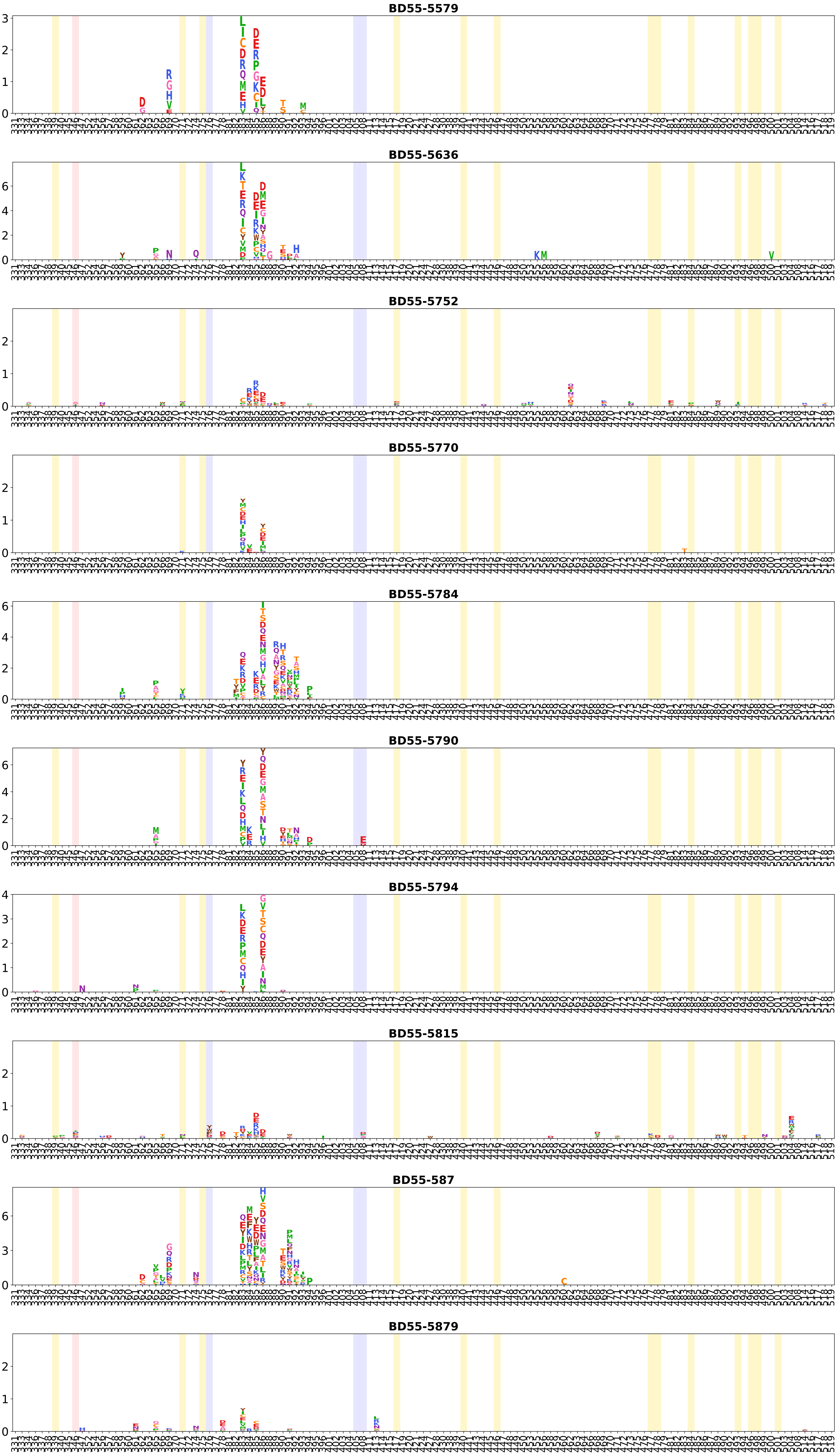
BD55-5579
BD55-5636
BD55-5752
BD55-5770
BD55-5784
BD55-5790
BD55-5794
BD55-5815
BD55-587
BD55-5879

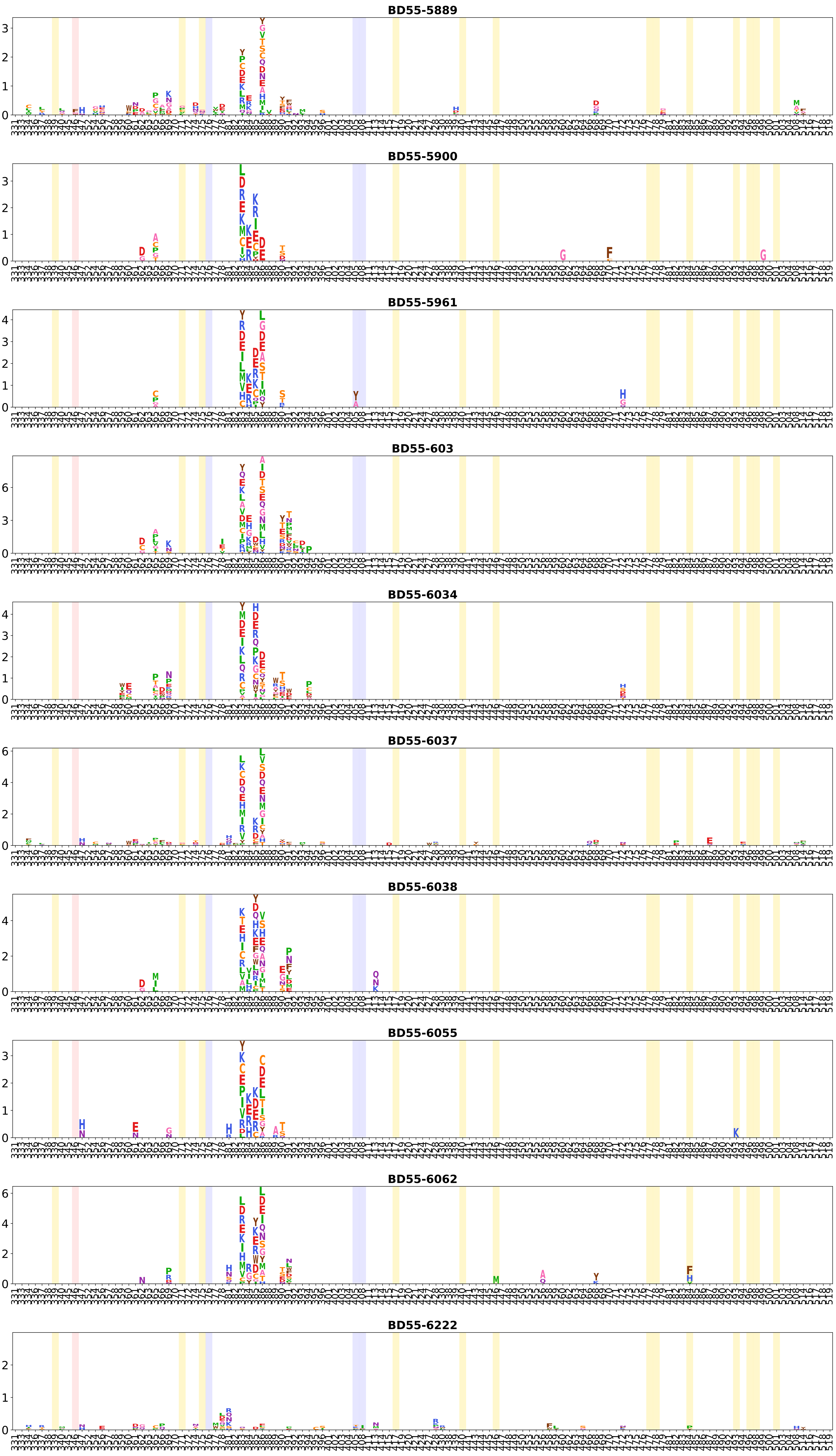
BD55-5889
BD55-5900
BD55-5961
BD55-603
BD55-6034
BD55-6037
BD55-6038
BD55-6055
BD55-6062
BD55-6222

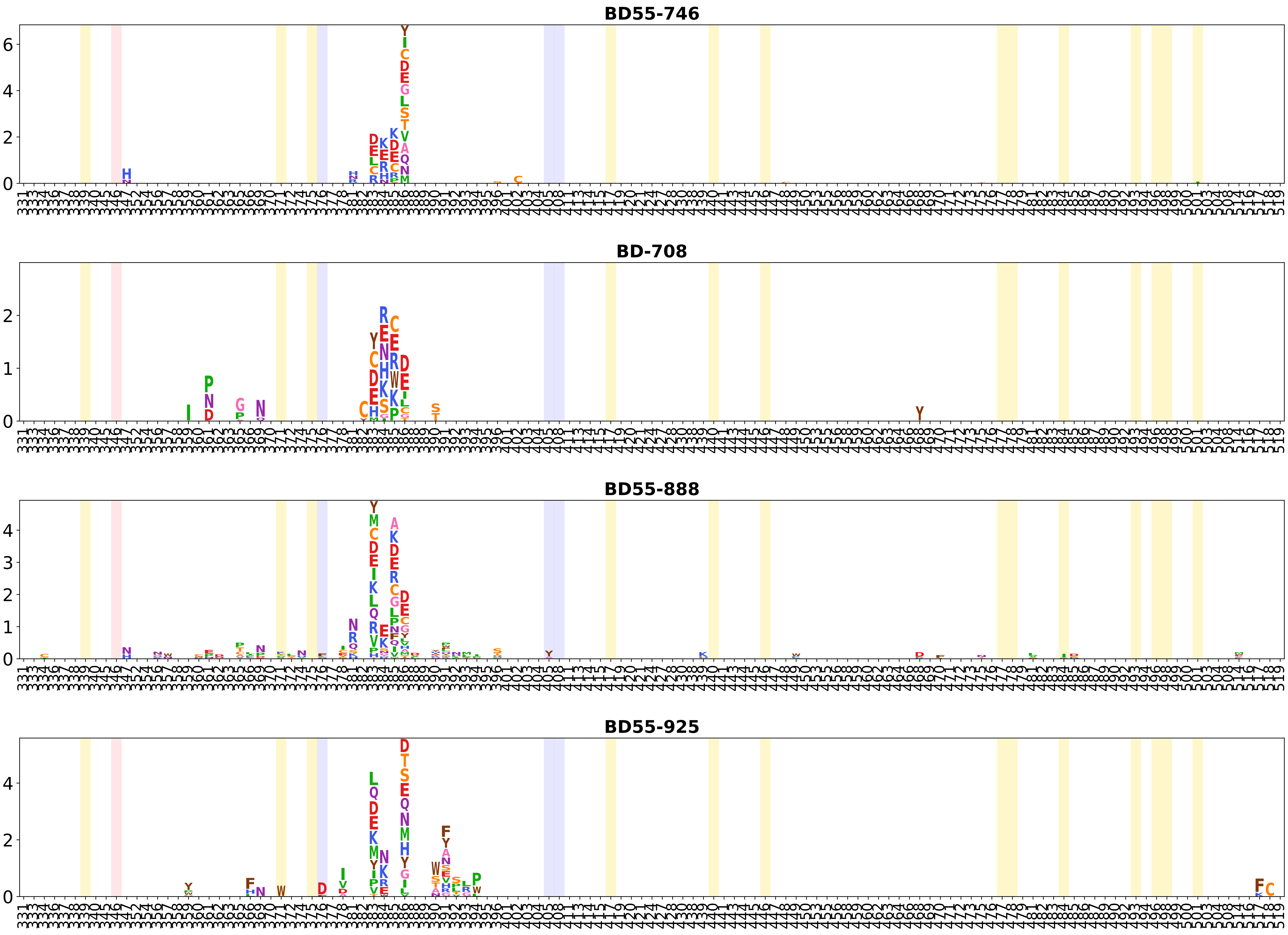
BD55-746
BD-708
BD55-888
BD55-925

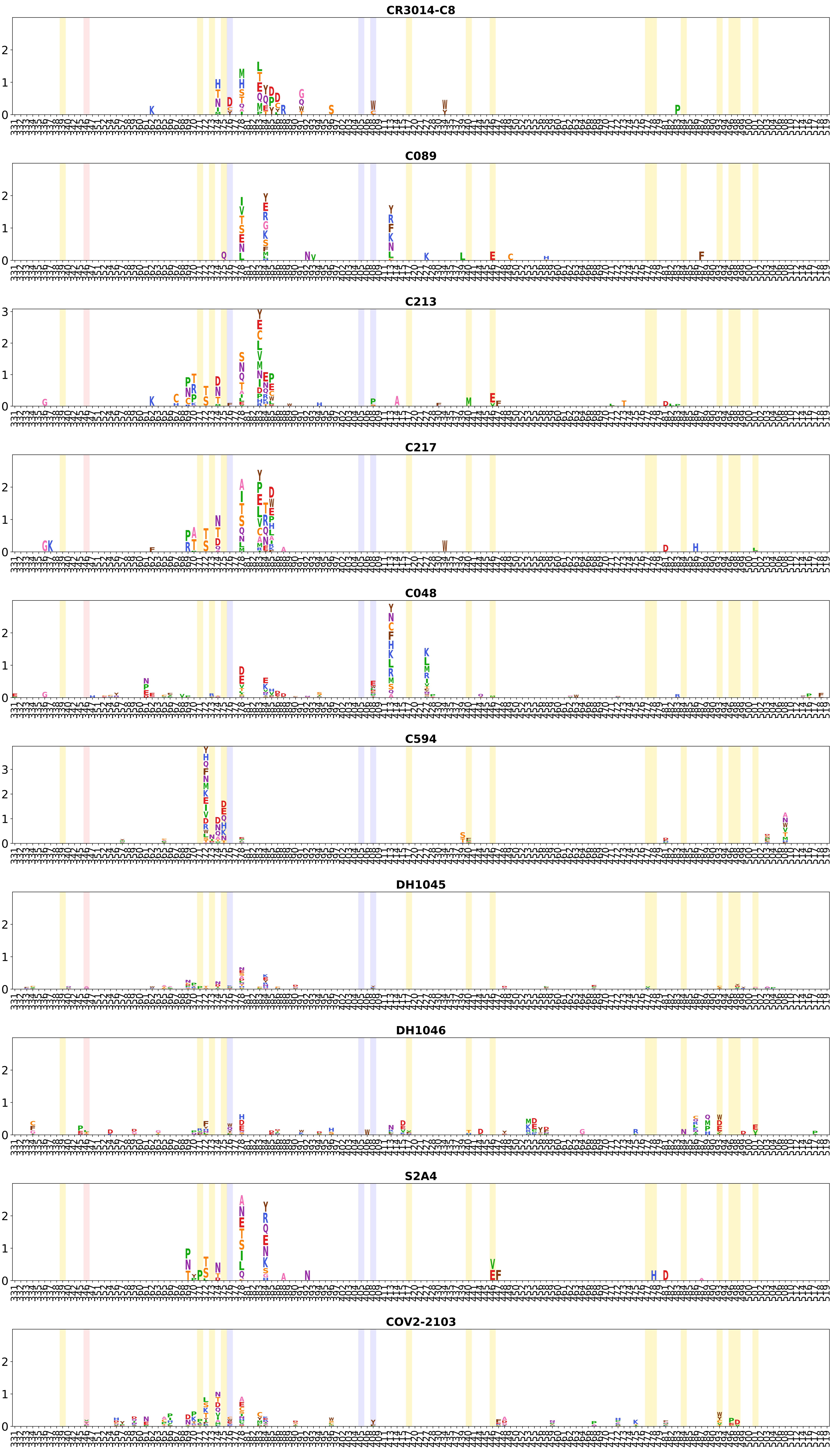
CR3014-C8
C089
C213
C217
C048
C594
DH1045
DH1046
S2A4
COV2-2103

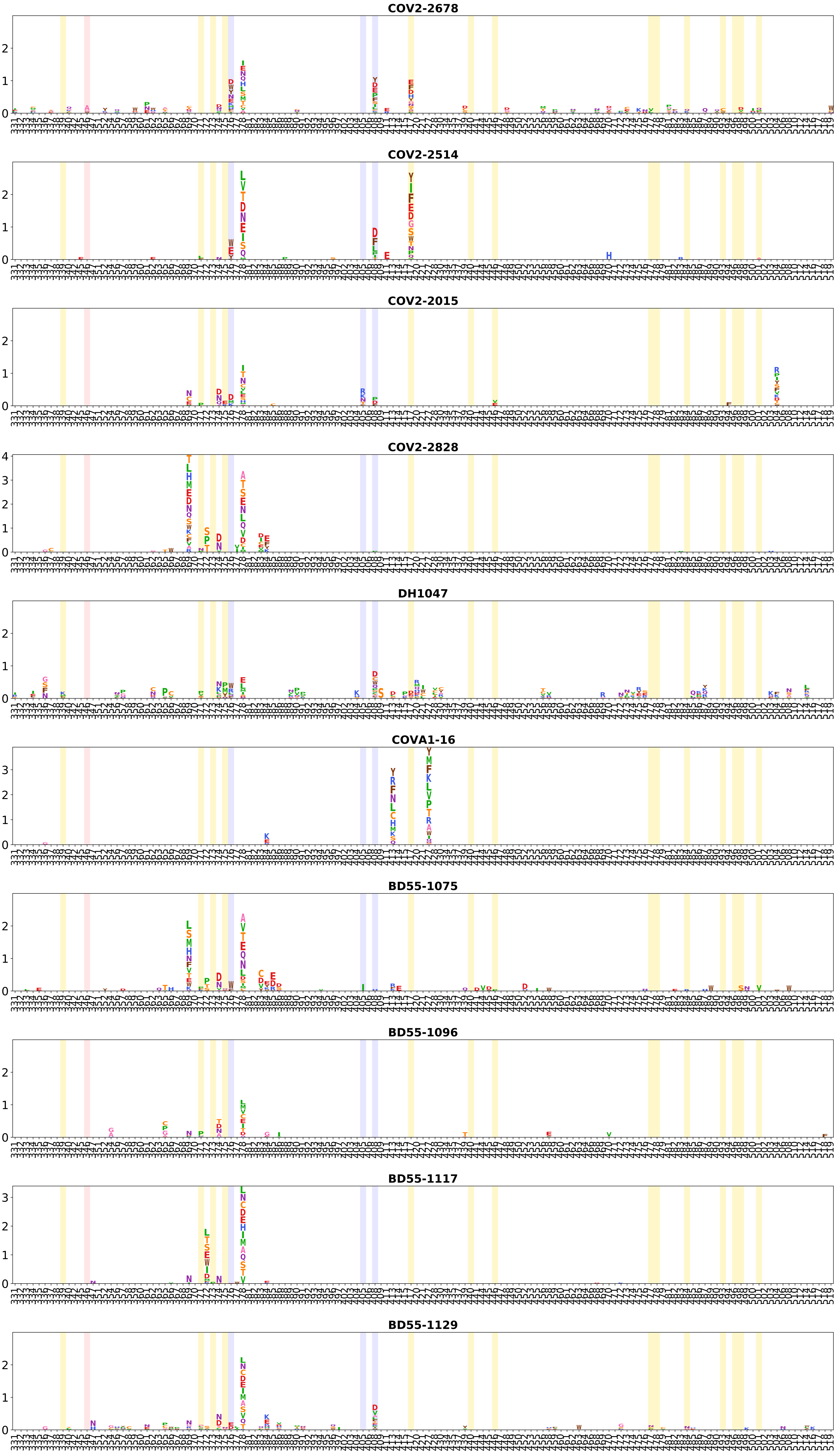
COV2-2678
COV2-2514
COV2-2015
COV2-2828
DH1047
COVA1-16
BD55-1075
BD55-1096
BD55-1117
BD55-1129

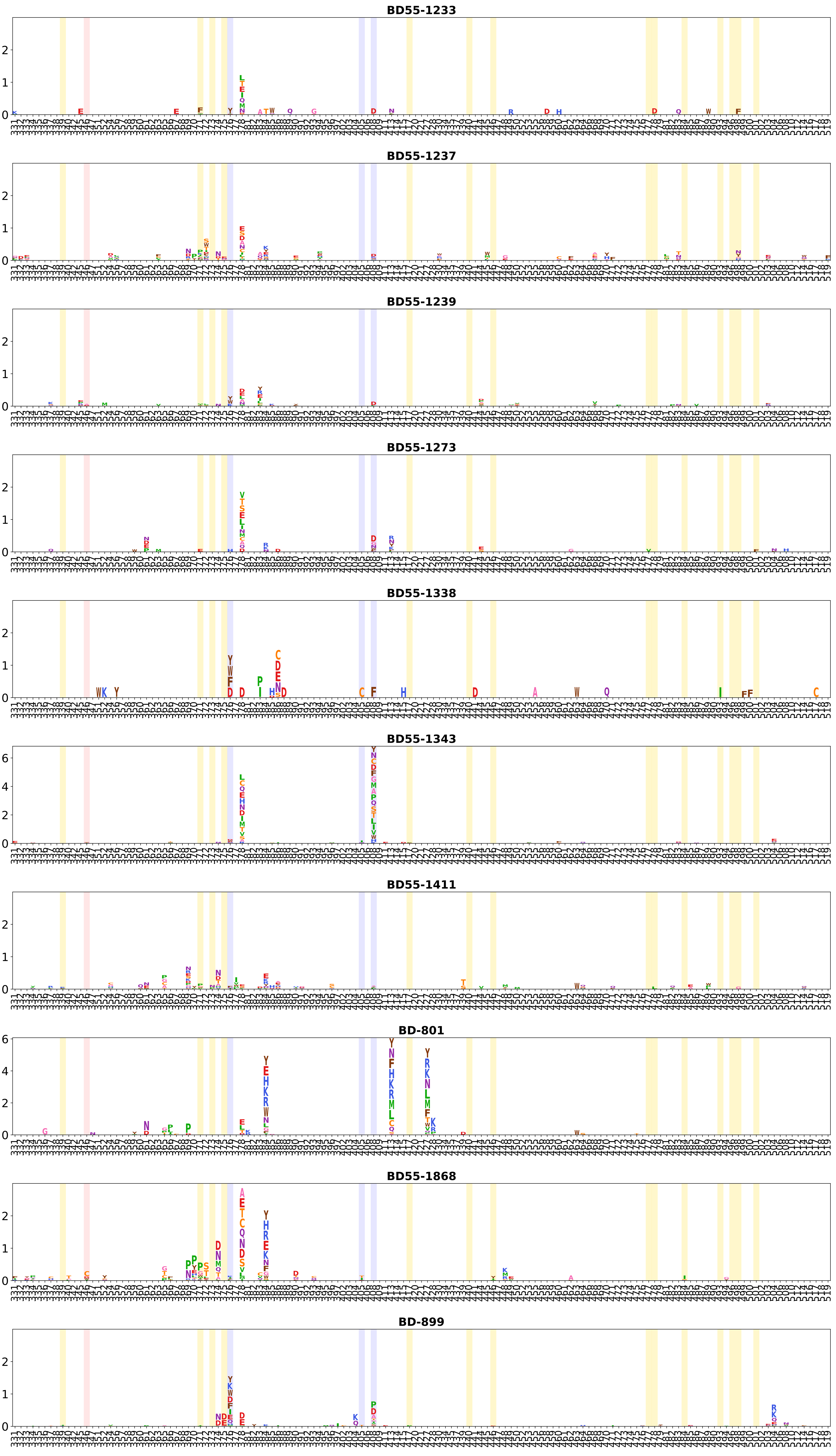
BD55-1233
BD55-1237
BD55-1239
BD55-1273
BD55-1338
BD55-1343
BD55-1411
BD-801
BD55-1868
BD-899

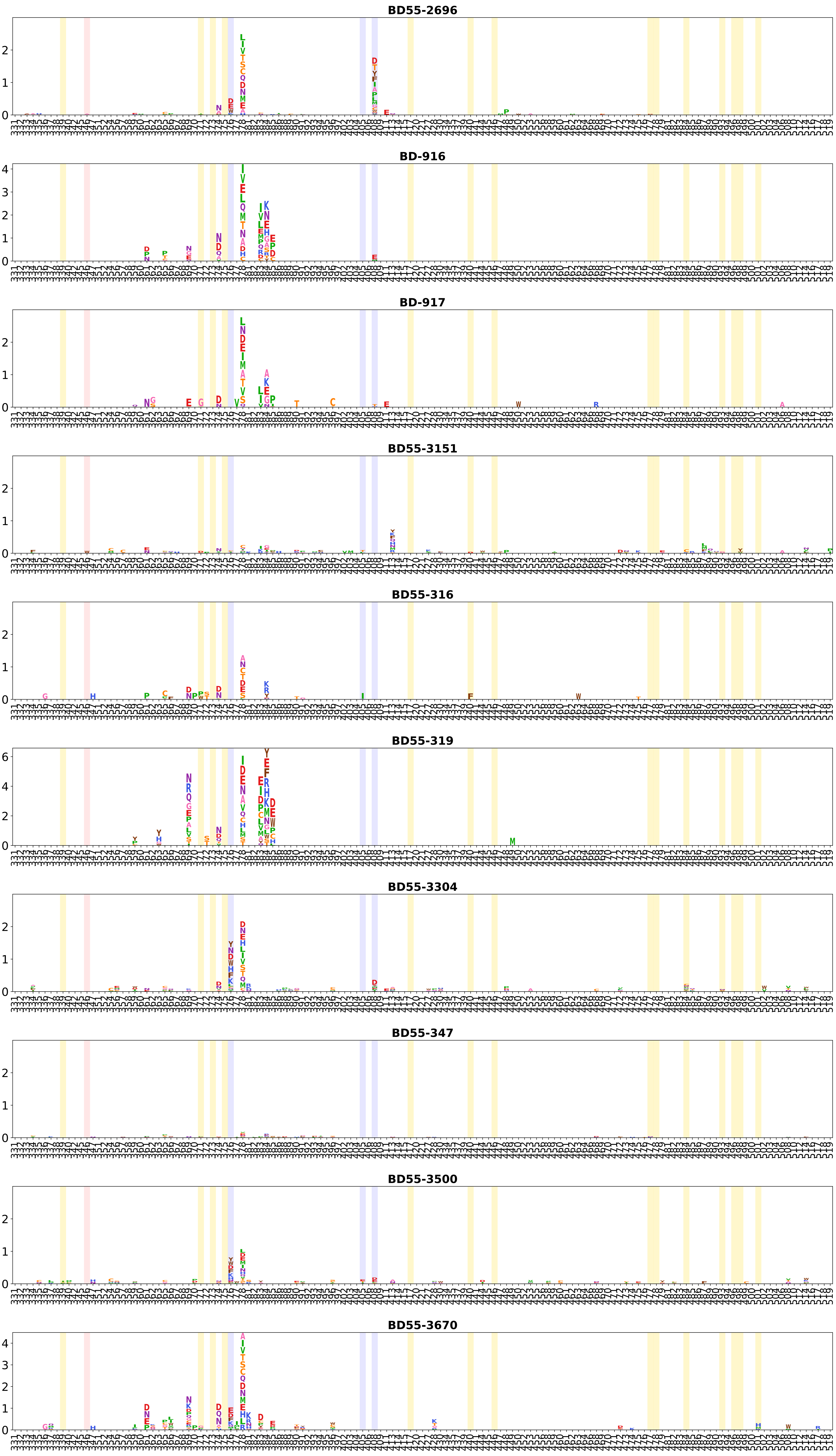
BD55-2696
BD-916
BD-917
BD55-3151
BD55-316
BD55-319
BD55-3304
BD55-347
BD55-3500
BD55-3670

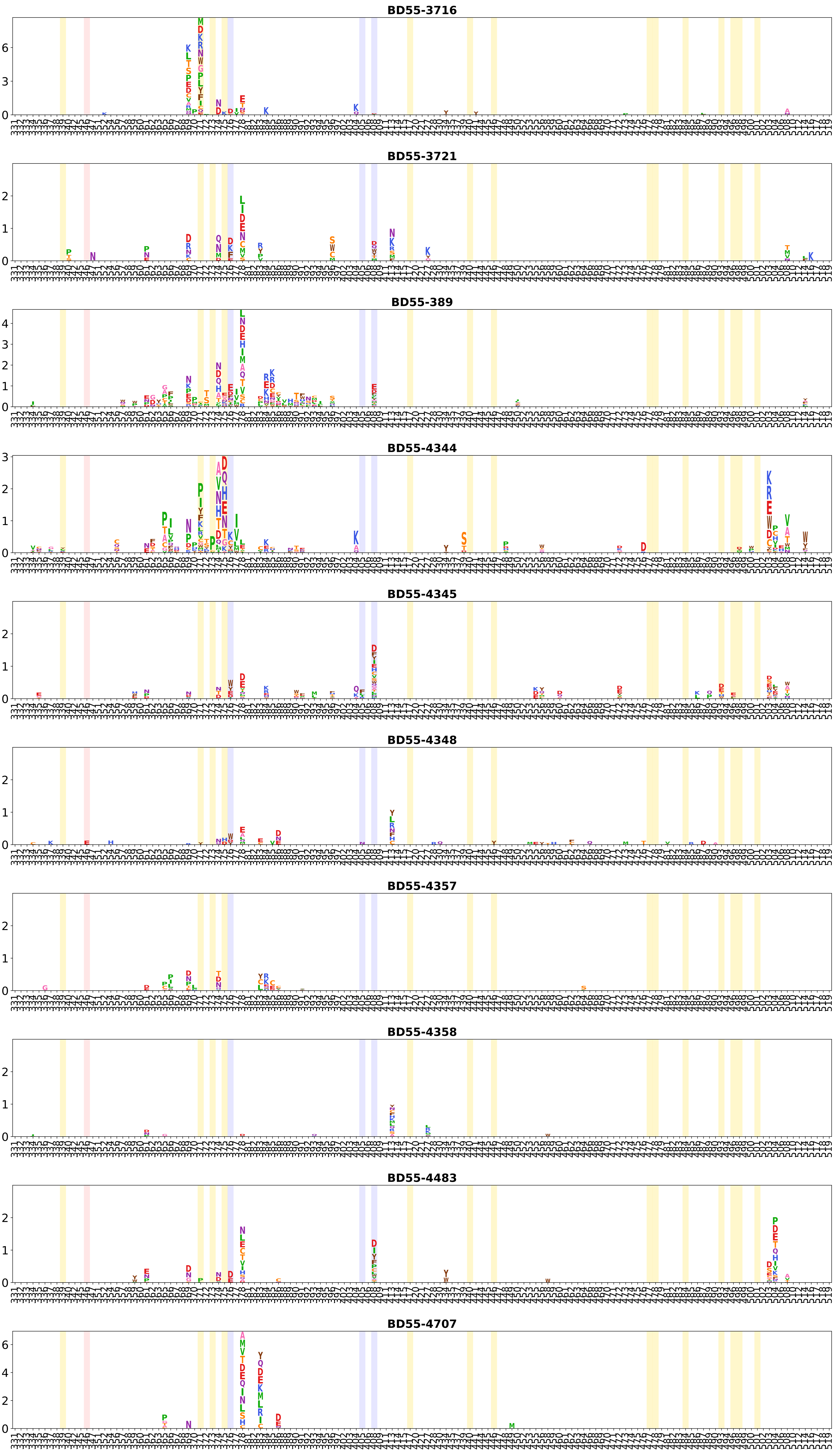

BD55-3716
BD55-3721
BD55-389
BD55-4344
BD55-4345
BD55-4348
BD55-4357
BD55-4358
BD55-4483
BD55-4707

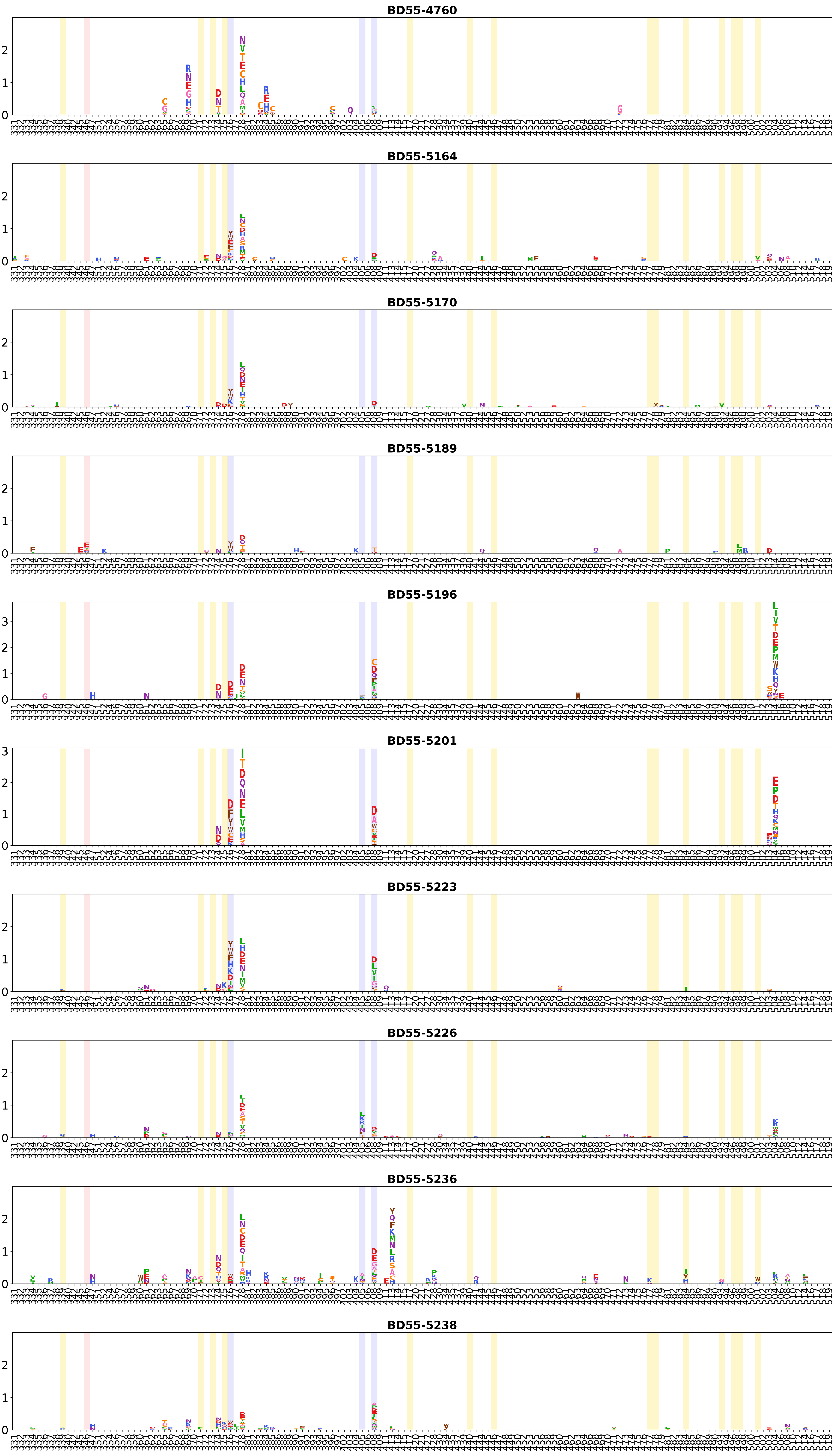
BD55-4760
BD55-5164
BD55-5170
BD55-5189
BD55-5196
BD55-5201
BD55-5223
BD55-5226
BD55-5236
BD55-5238

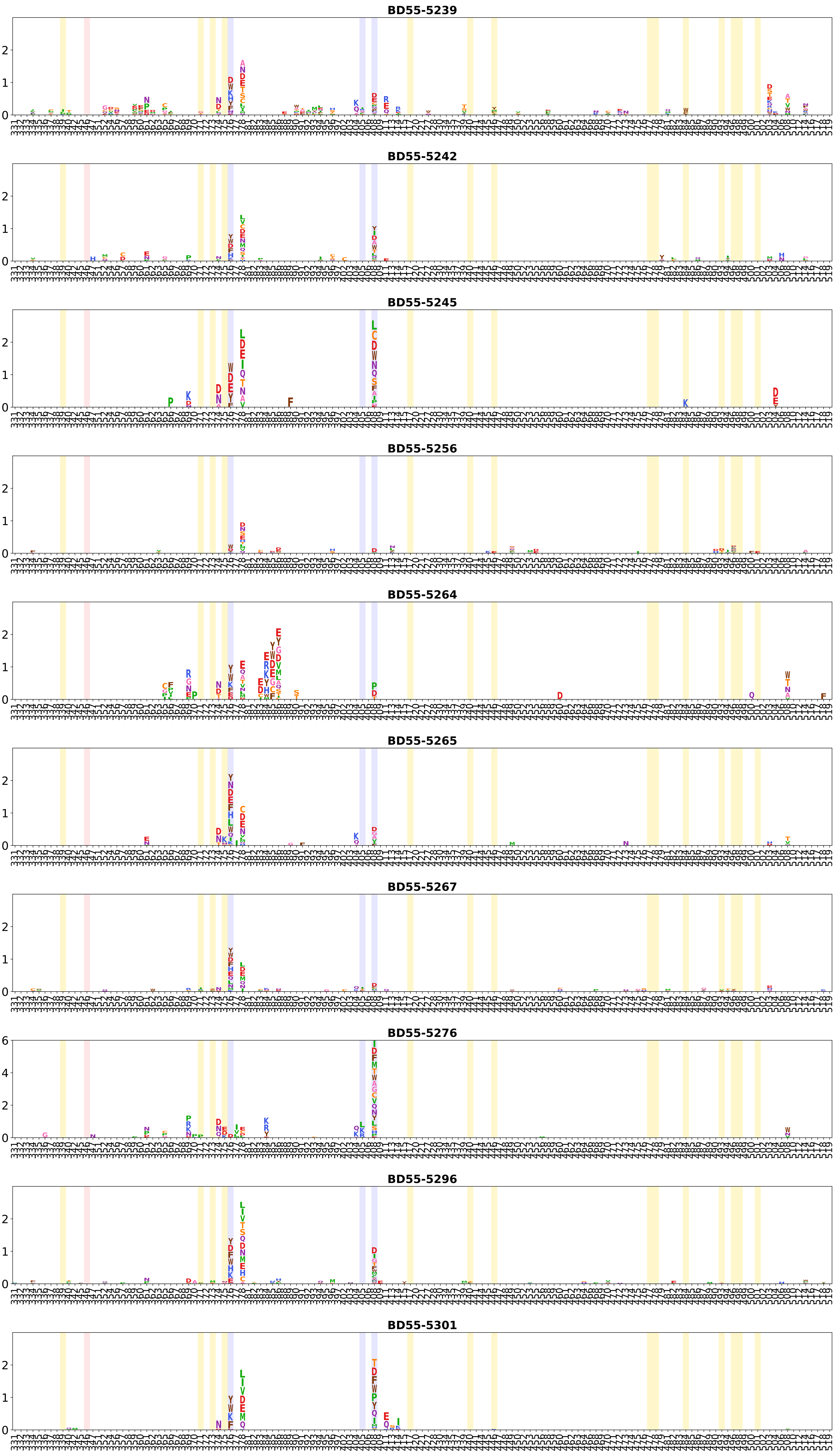
BD55-5239
BD55-5242
BD55-5245
BD55-5256
BD55-5264
BD55-5265
BD55-5267
BD55-5276
BD55-5296
BD55-5301

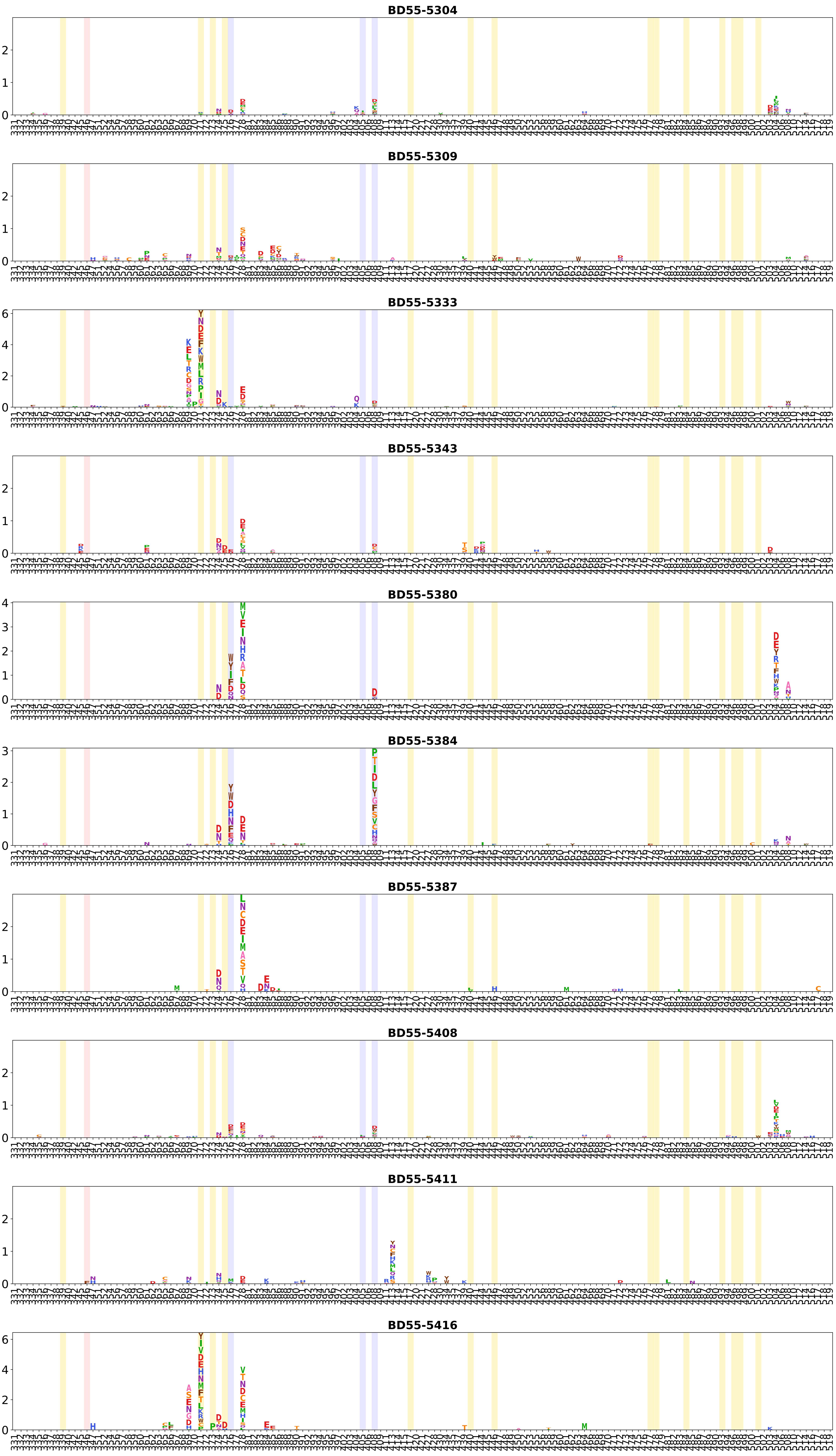
BD55-5304
BD55-5309
BD55-5333
BD55-5343
BD55-5380
BD55-5384
BD55-5387
BD55-5408
BD55-5411
BD55-5416

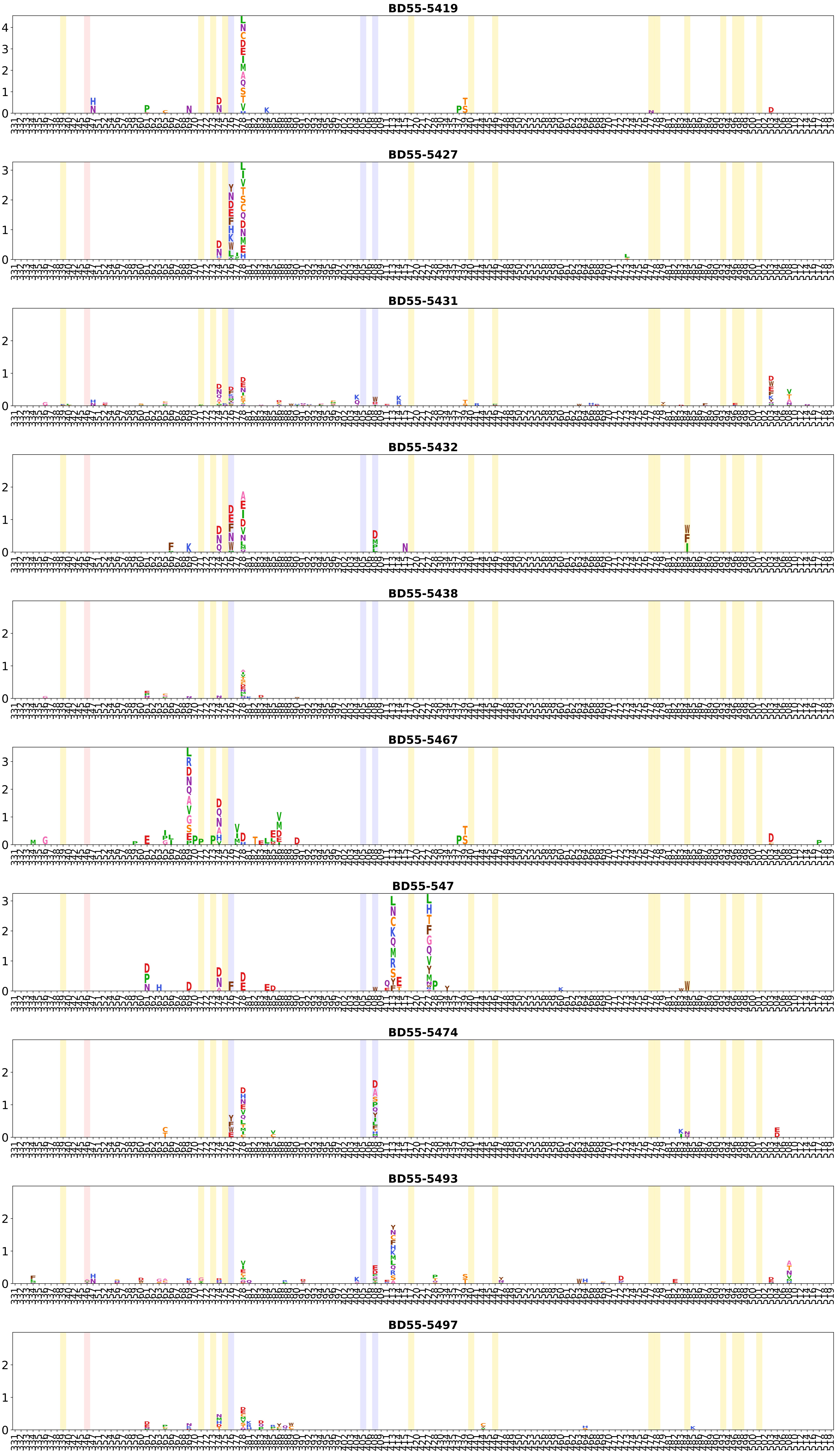
BD55-5419
BD55-5427
BD55-5431
BD55-5432
BD55-5438
BD55-5467
BD55-547
BD55-5474
BD55-5493
BD55-5497

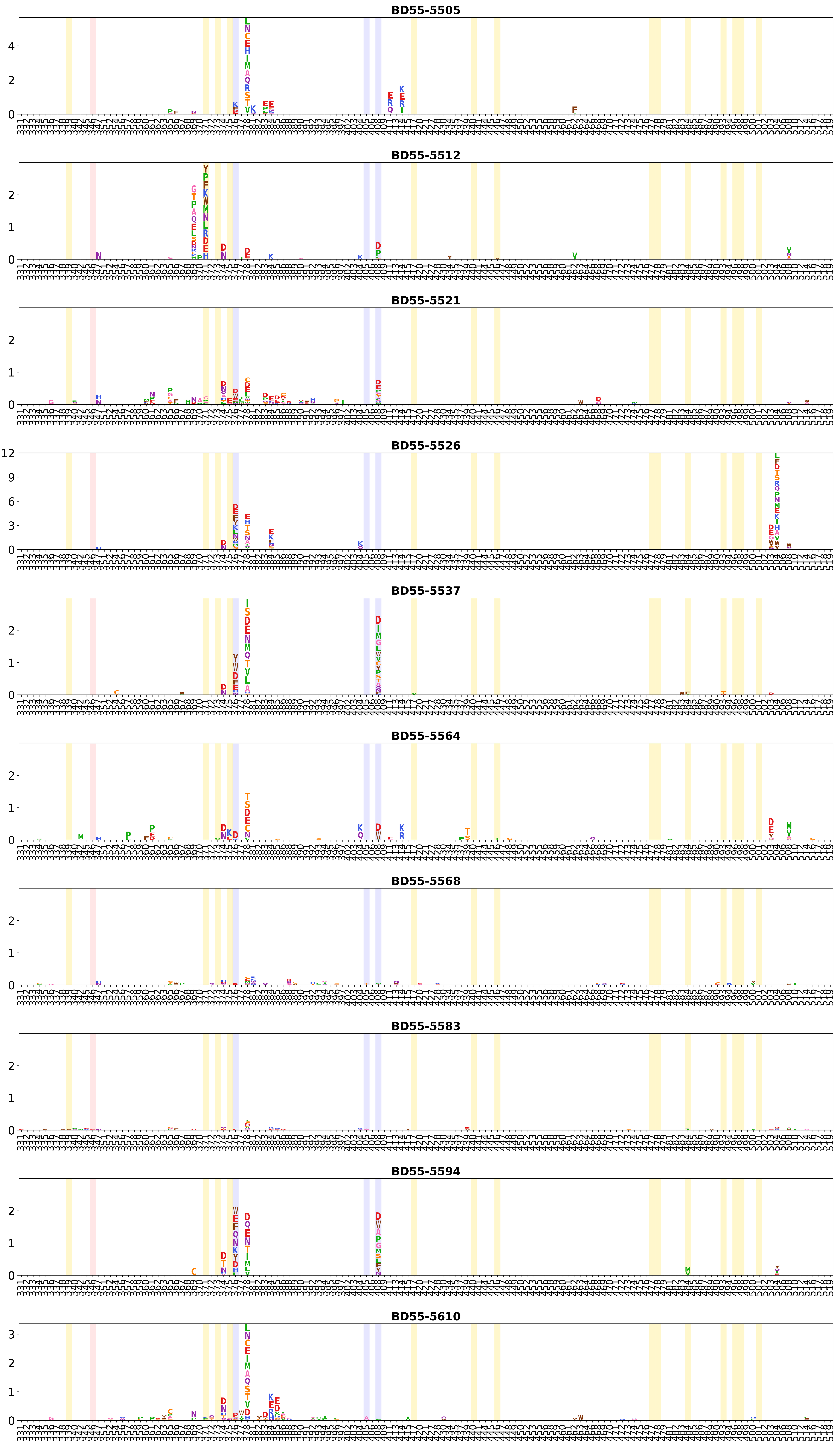
BD55-5505
BD55-5512
BD55-5521
BD55-5526
BD55-5537
BD55-5564
BD55-5568
BD55-5583
BD55-5594
BD55-5610

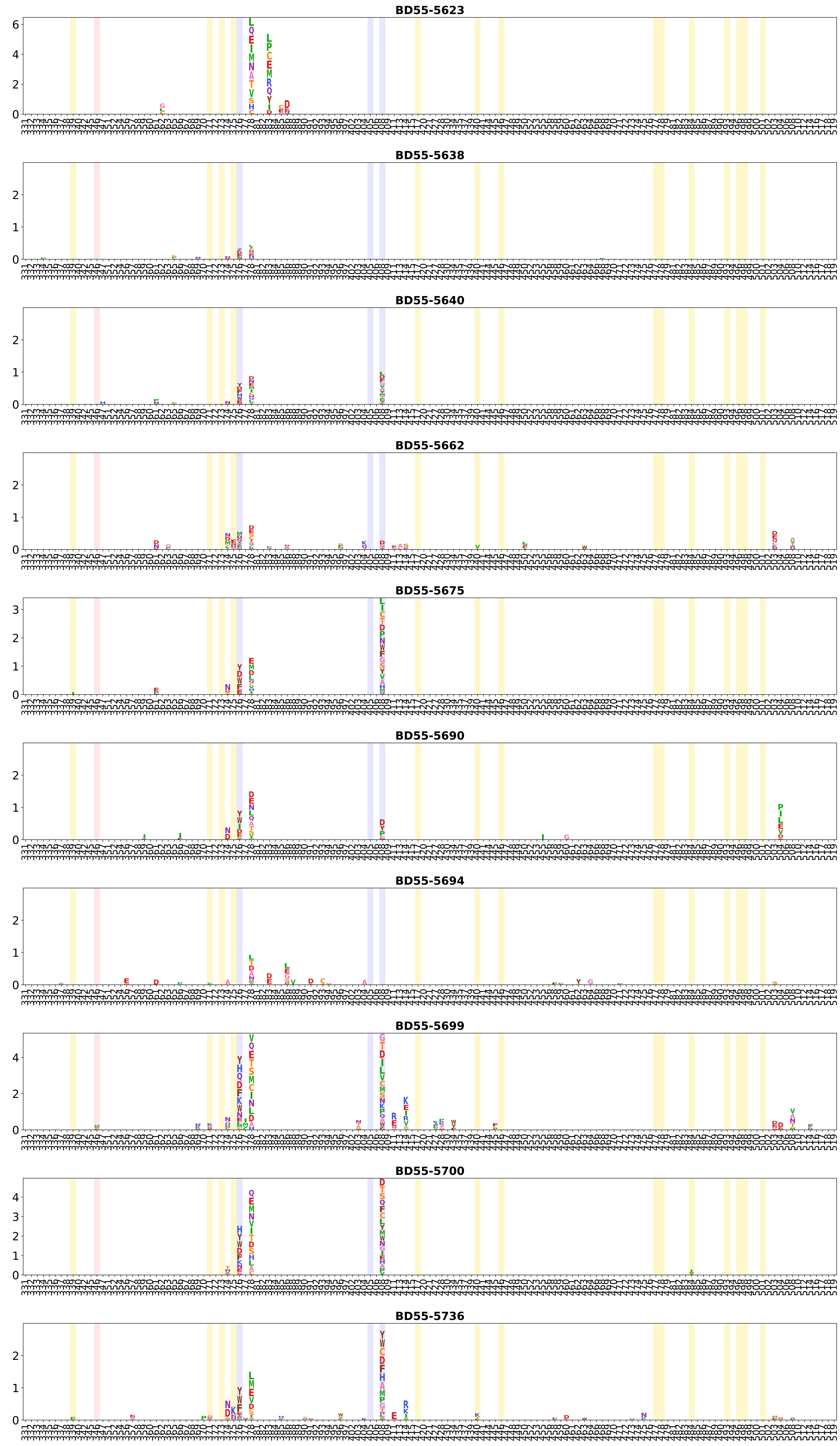
BD55-5623
BD55-5638
BD55-5640
BD55-5662
BD55-5675
BD55-5690
BD55-5694
BD55-5699
BD55-5700
BD55-5736

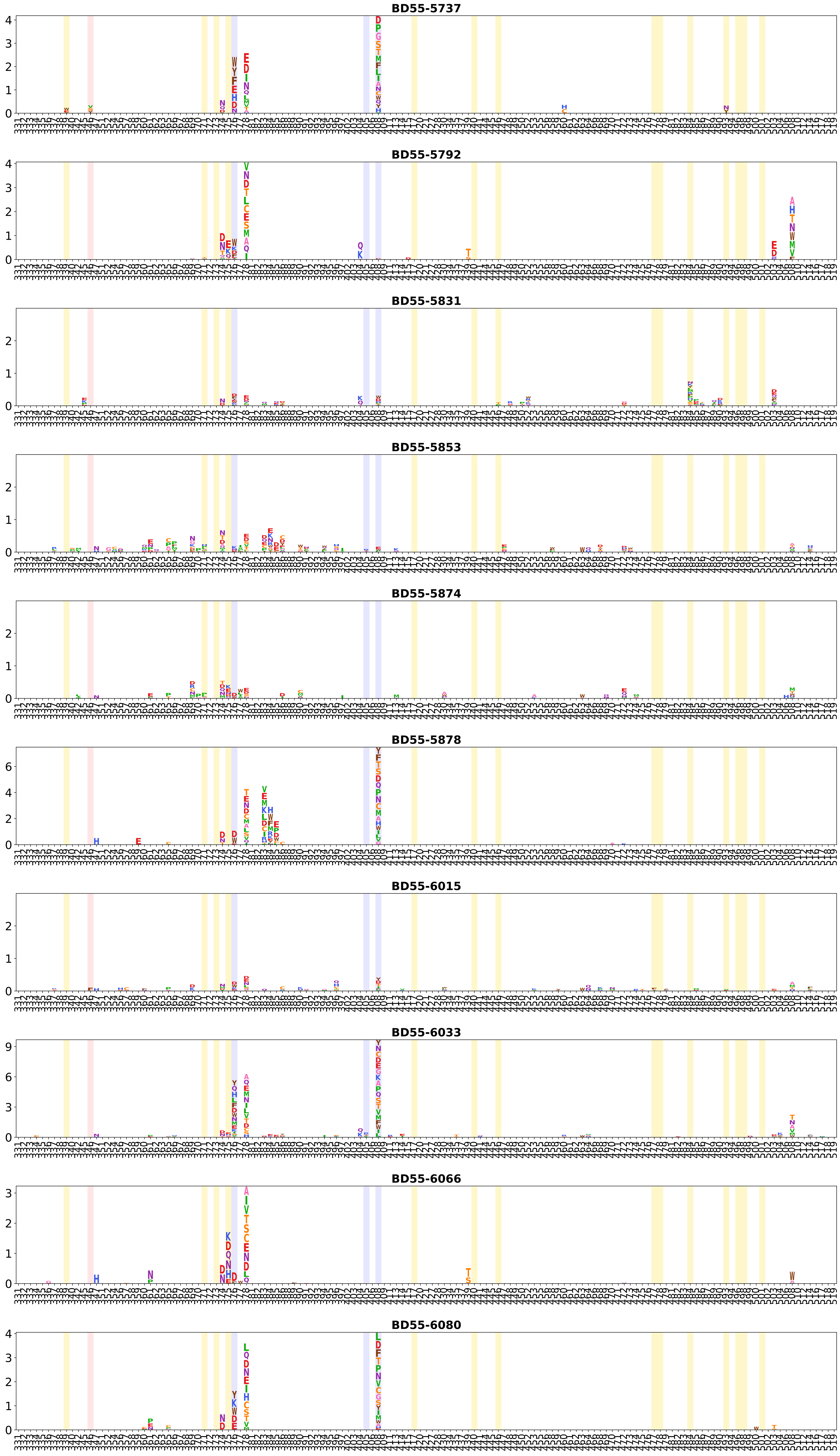
BD55-5737
BD55-5792
BD55-5831
BD55-5853
BD55-5874
BD55-5878
BD55-6015
BD55-6033
BD55-6066
BD55-6080

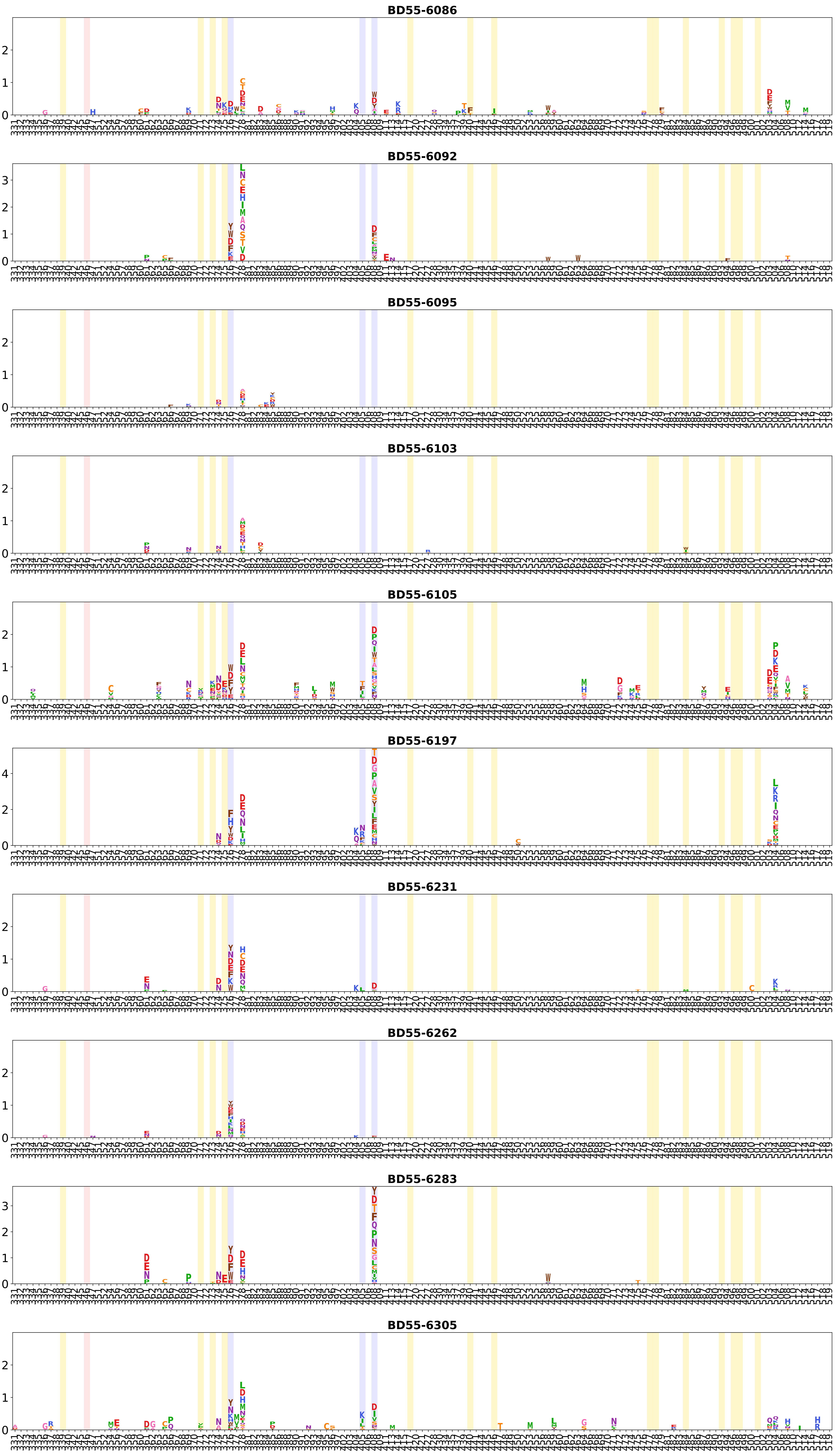
BD55-6086
BD55-6092
BD55-6095
BD55-6103
BD55-6105
BD55-6197
BD55-6231
BD55-6262
BD55-6283
BD55-6305

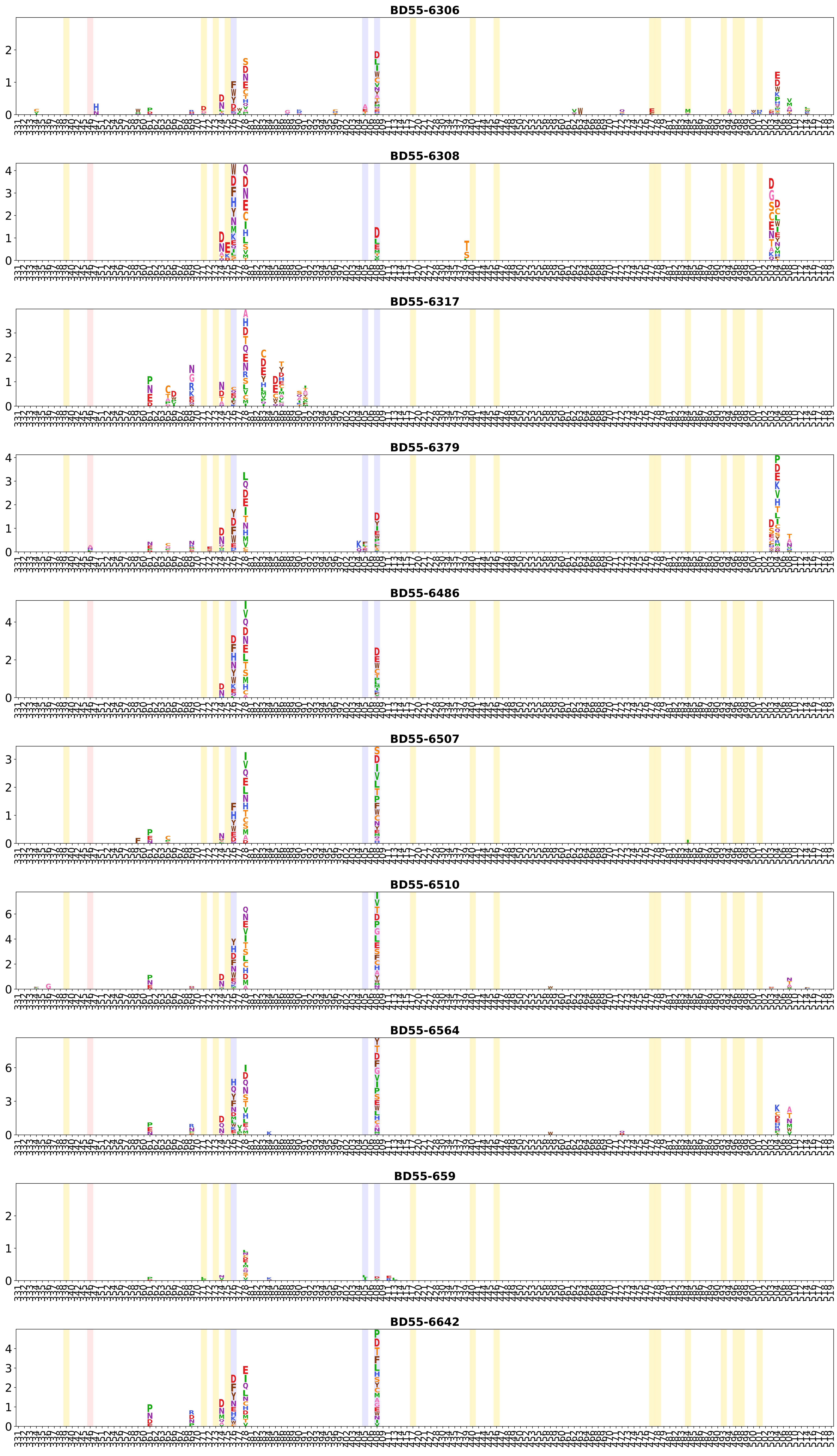
BD55-6306
BD55-6308
BD55-6317
BD55-6379
BD55-6486
BD55-6507
BD55-6510
BD55-6564
BD55-659
BD55-6642

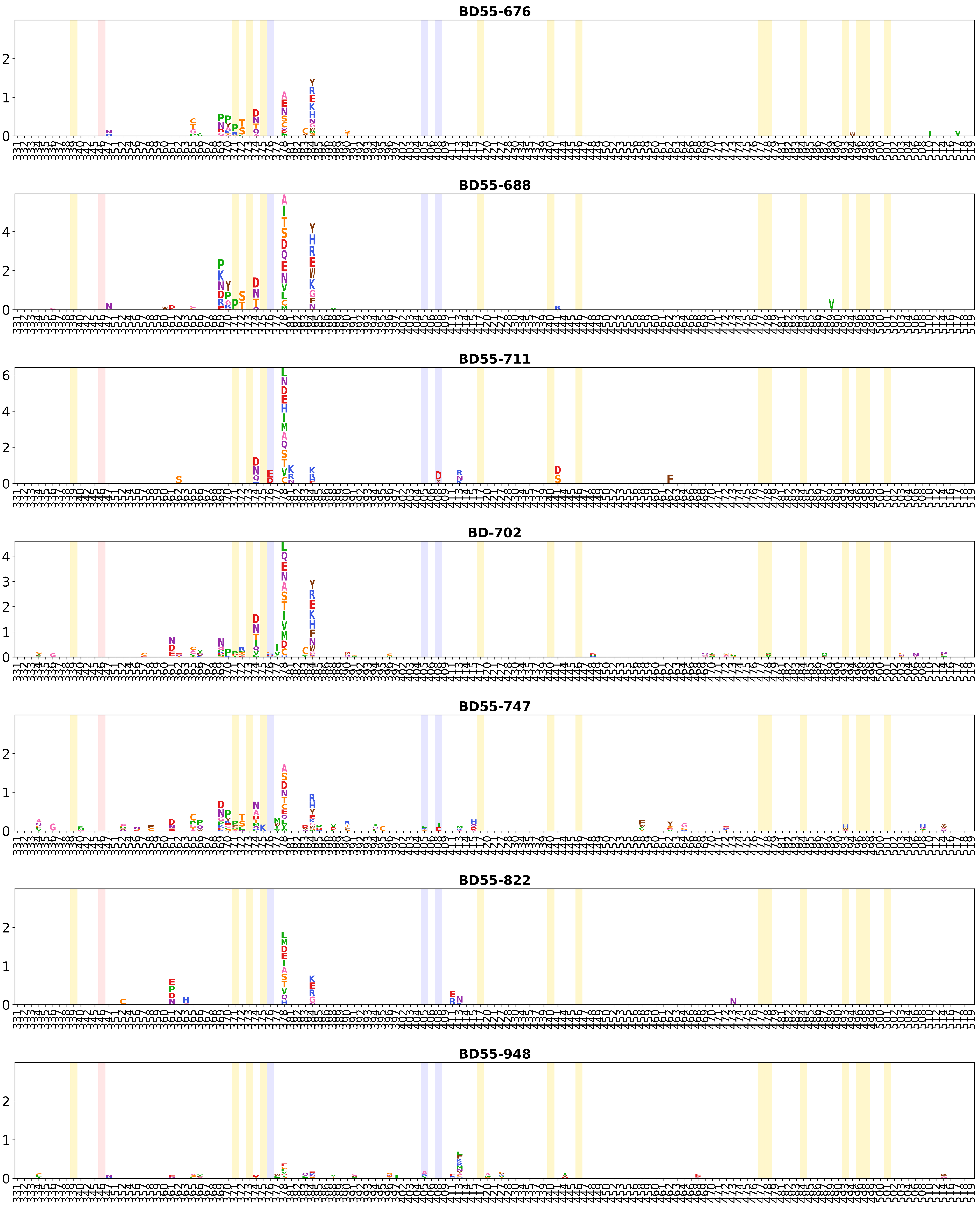
BD55-676
BD55-688
BD55-711
BD-702
BD55-747
BD55-822
BD55-948

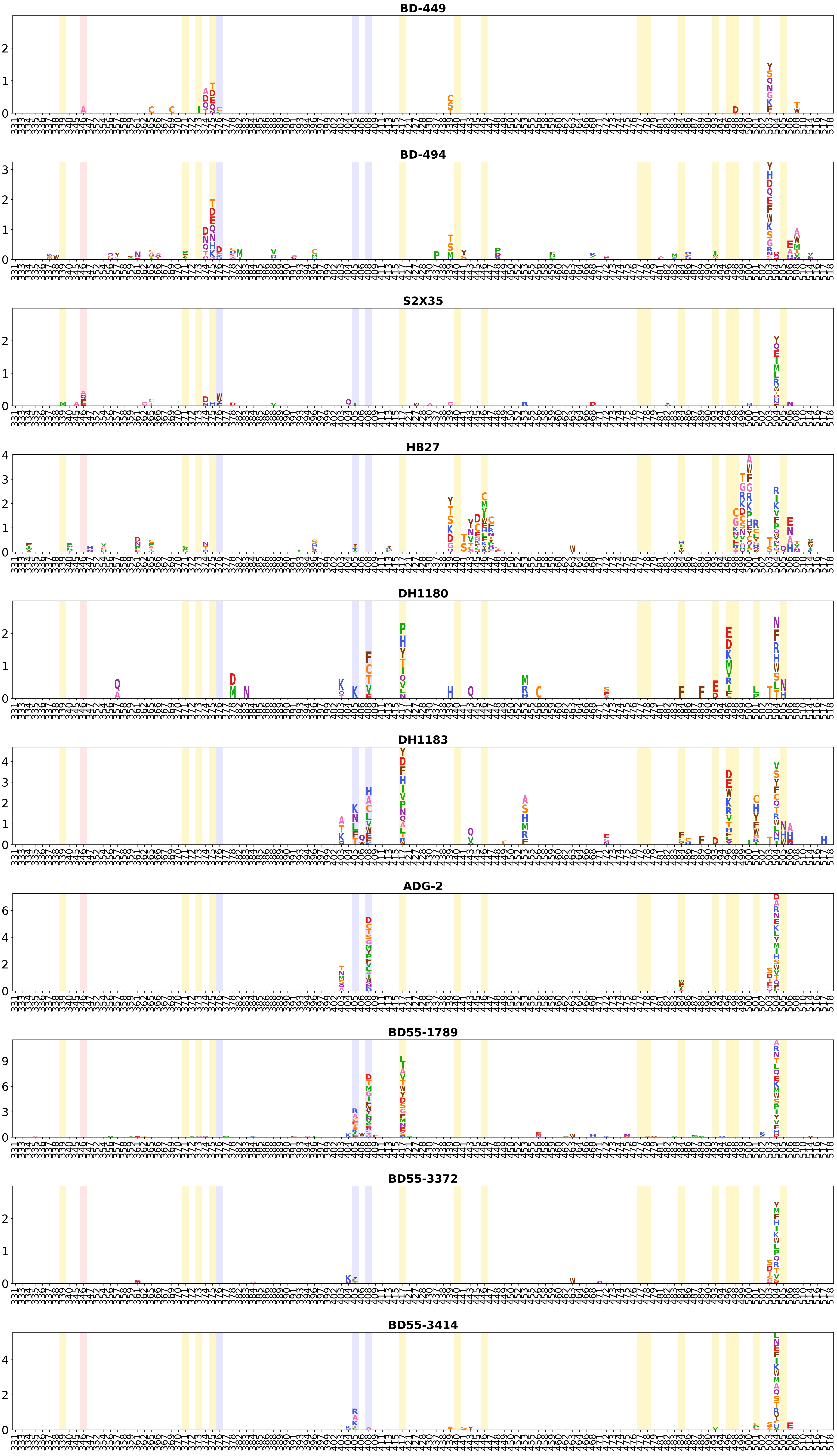
BD-449
BD-494
S2X35
HB27
DH1180
DH1183
ADG-2
BD55-1789
BD55-3372
BD55-3414

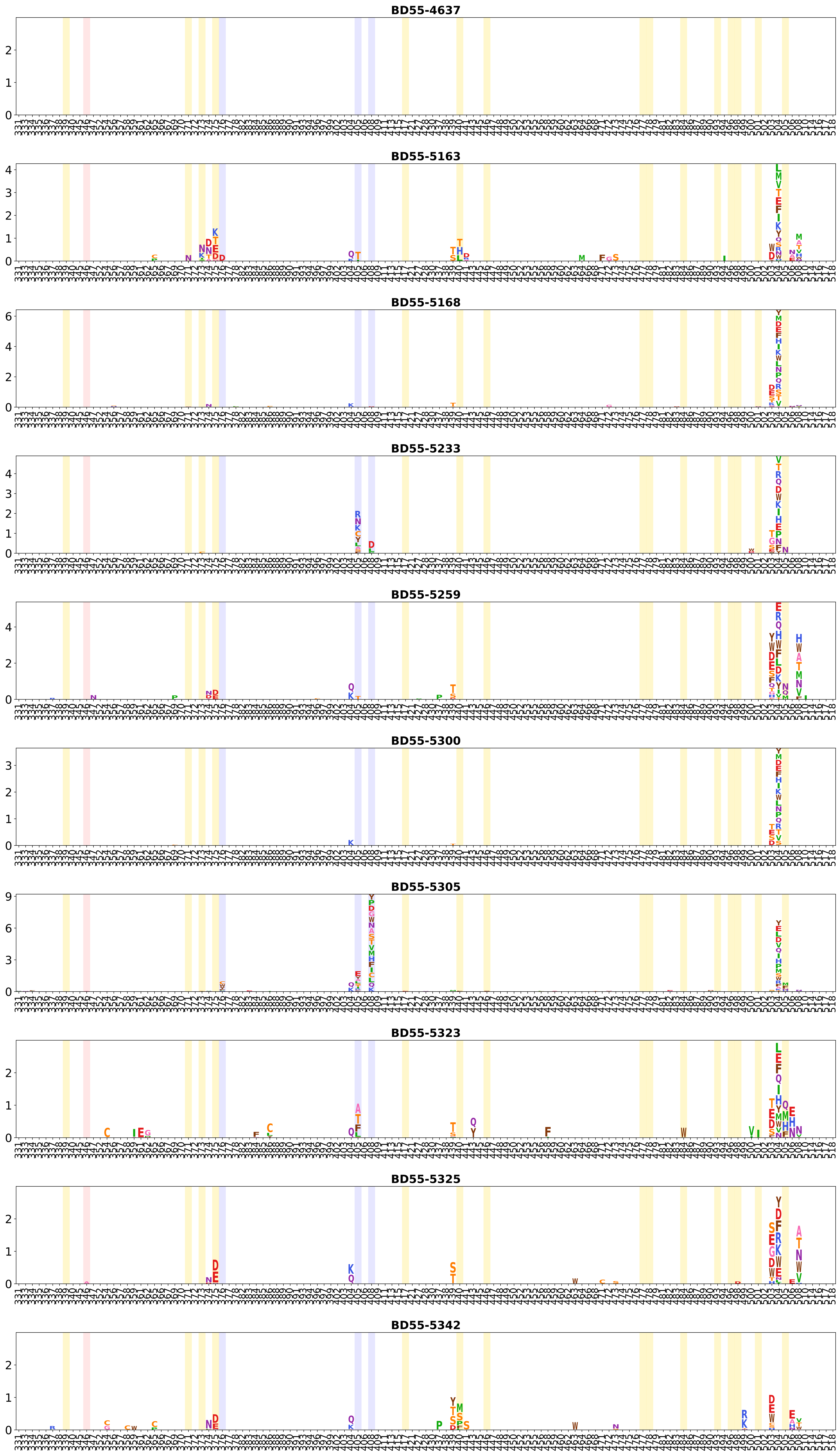
BD55-4637
BD55-5163
BD55-5168
BD55-5233
BD55-5259
BD55-5300
BD55-5305
BD55-5323
BD55-5325
BD55-5342

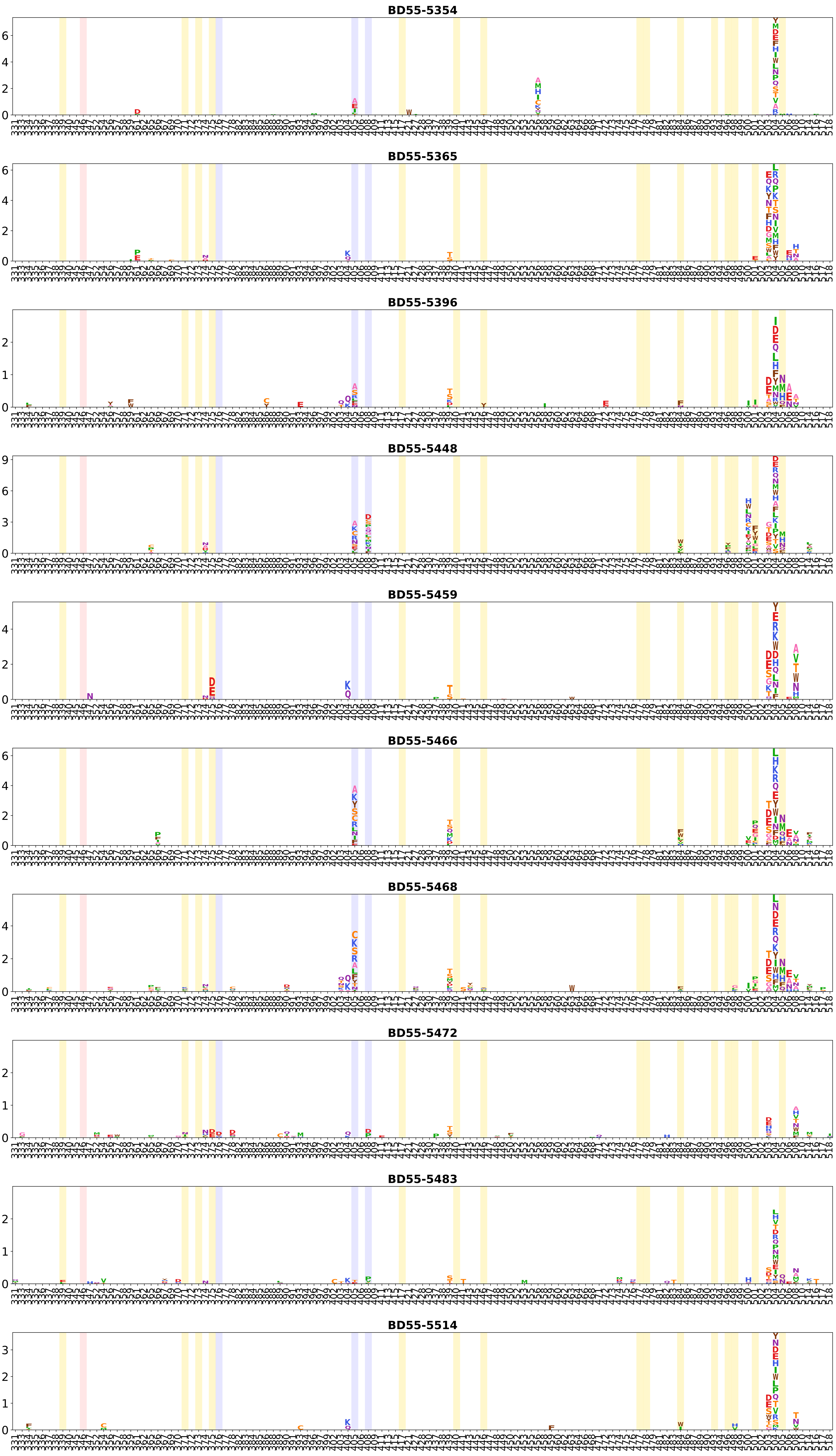
BD55-5354
BD55-5365
BD55-5396
BD55-5448
BD55-5459
BD55-5466
BD55-5468
BD55-5472
BD55-5483
BD55-5514

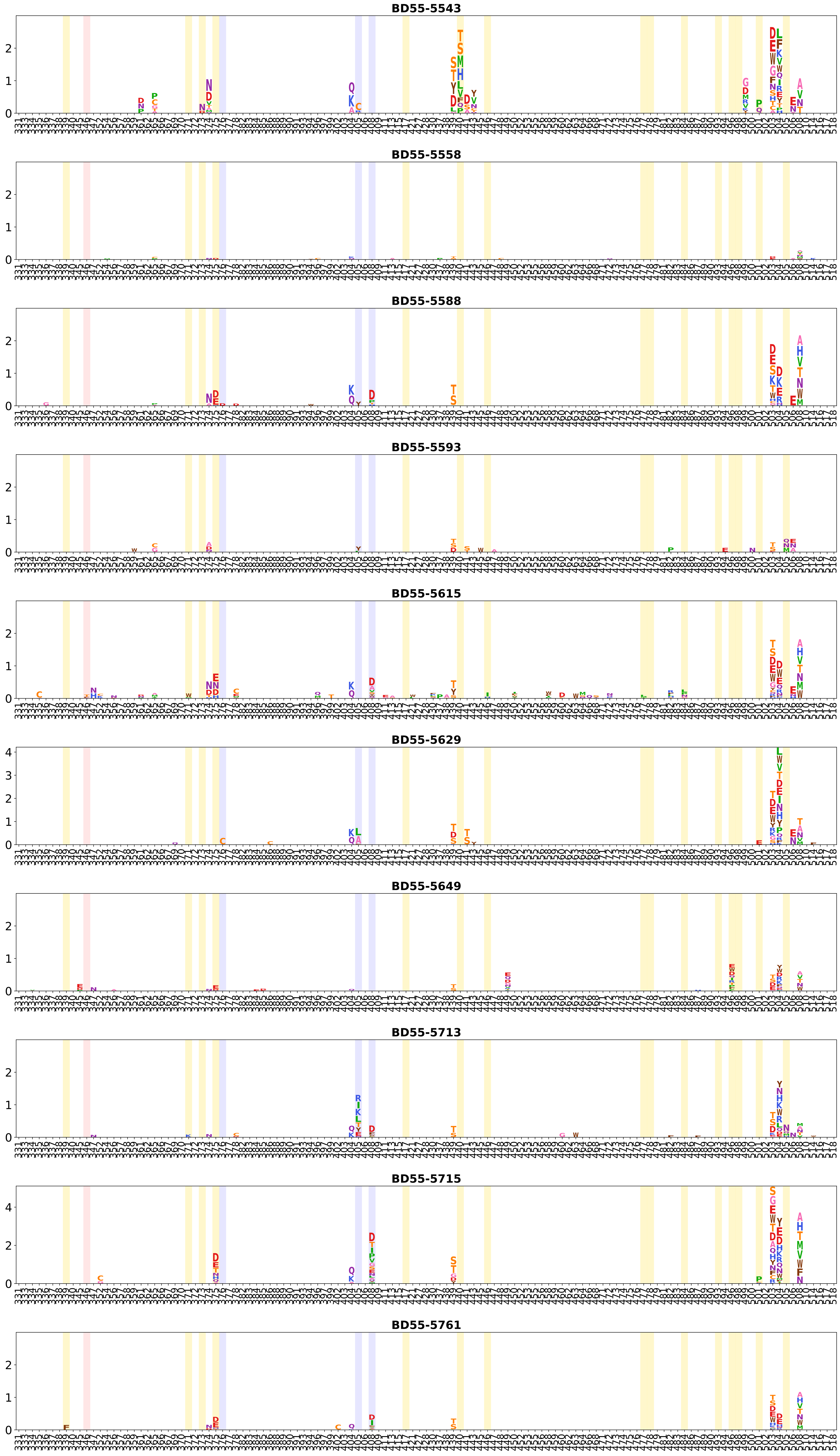
BD55-5543
BD55-5558
BD55-5588
BD55-5593
BD55-5615
BD55-5629
BD55-5649
BD55-5713
BD55-5715
BD55-5761

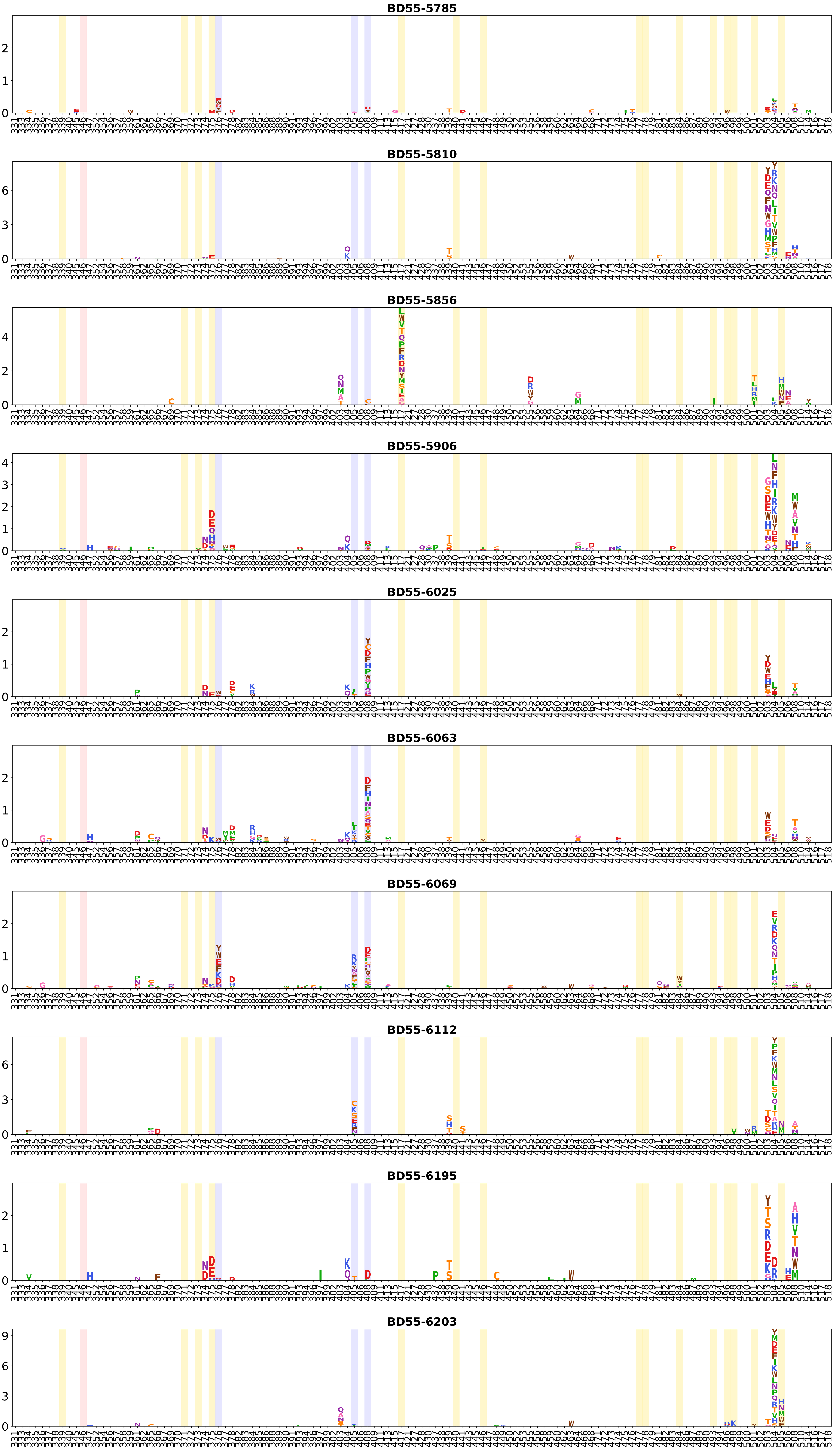
BD55-5785
BD55-5810
BD55-5856
BD55-5906
BD55-6025
BD55-6063
BD55-6069
BD55-6112
BD55-6195
BD55-6203

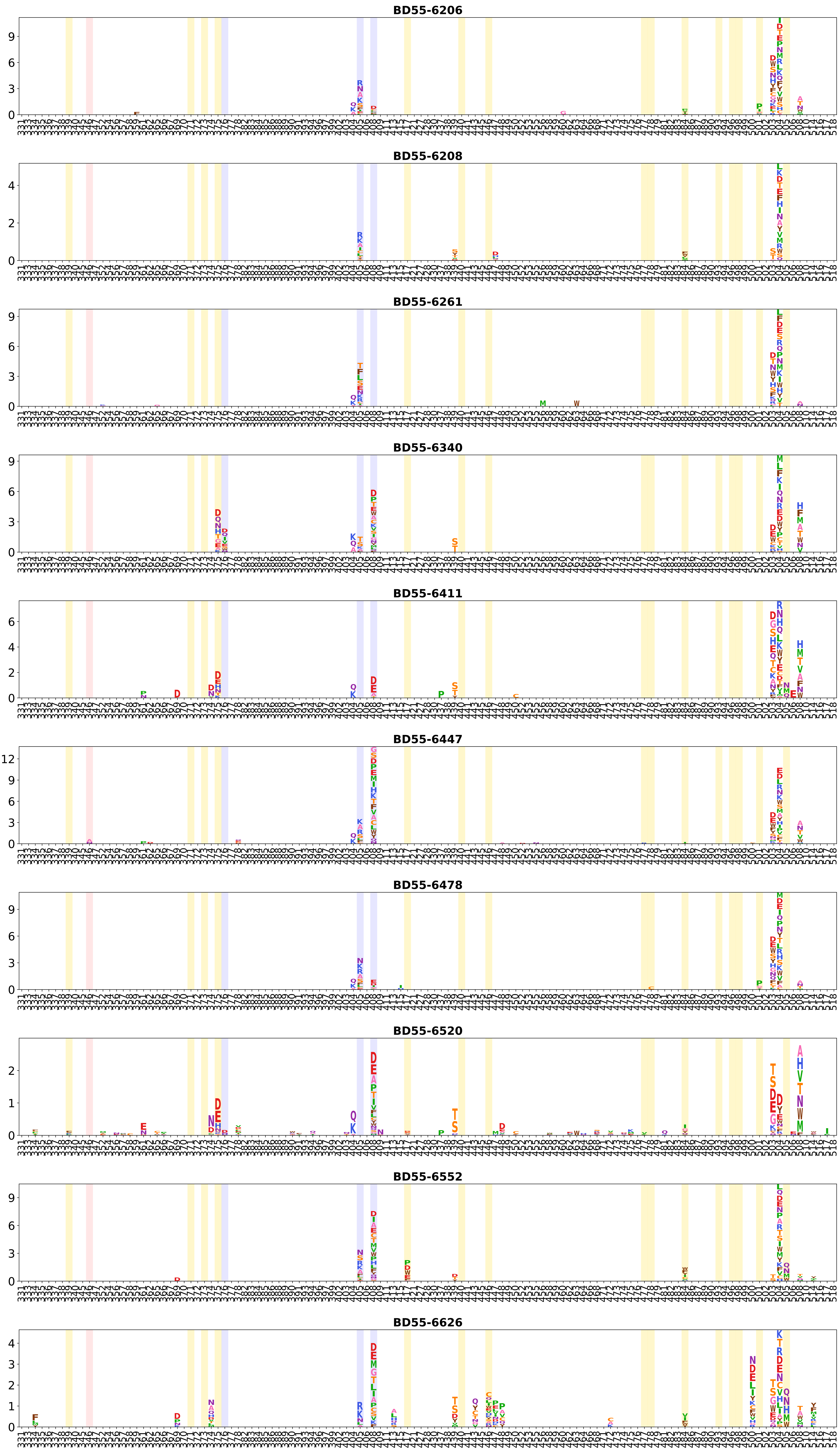

BD55-6206
BD55-6208
BD55-6261
BD55-6340
BD55-6411
BD55-6447
BD55-6478
BD55-6520
BD55-6552
BD55-6626

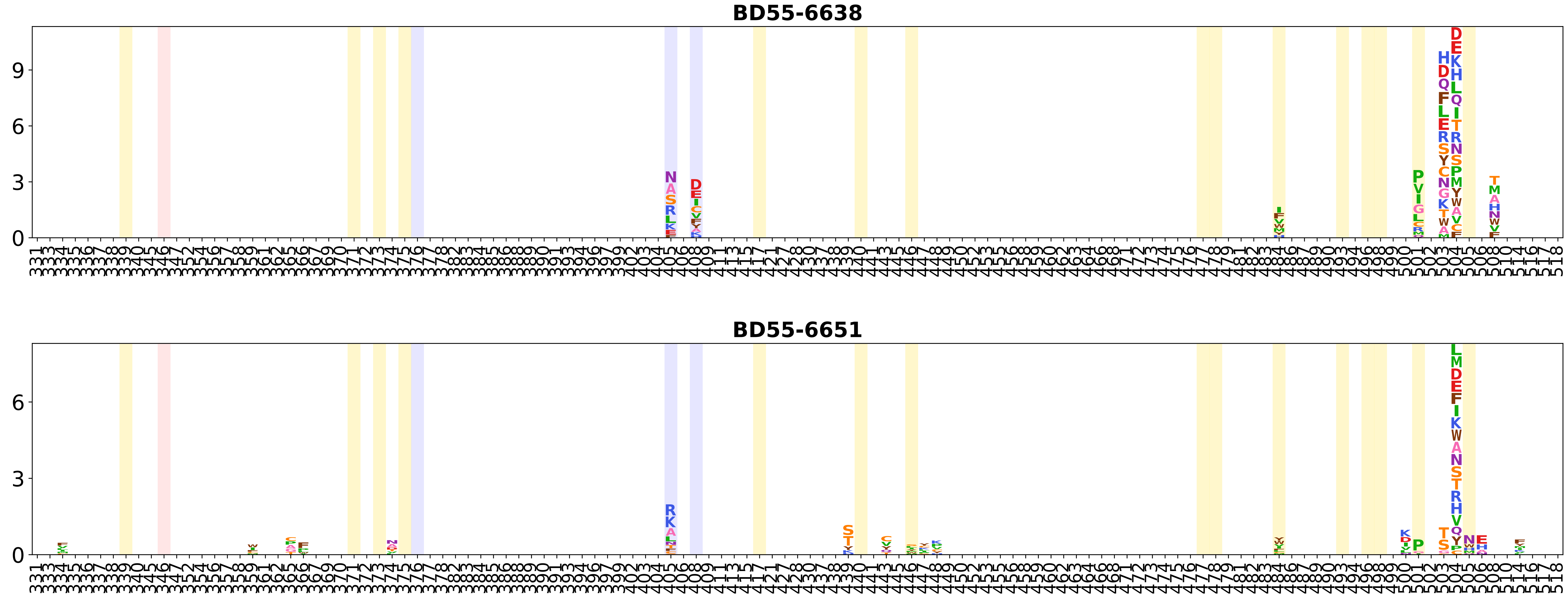
BD55-6638
BD55-6651
